# Integrative analysis reveals therapeutic potential of pyrvinium pamoate in Merkel cell carcinoma

**DOI:** 10.1101/2023.11.01.565218

**Authors:** Jiawen Yang, James T. Lim, Paul Victor, Marcelo G. Corona, Chen Chen, Hunain Khawaja, Qiong Pan, Gillian D. Paine-Murrieta, Rick G. Schnellmann, Denise J. Roe, Prafulla C. Gokhale, James A. DeCaprio, Megha Padi

## Abstract

Merkel Cell Carcinoma (MCC) is an aggressive neuroendocrine cutaneous malignancy arising from either ultraviolet-induced mutagenesis or Merkel cell polyomavirus (MCPyV) integration. Despite extensive research, our understanding of the molecular mechanisms driving the transition from normal cells to MCC remains limited. To address this knowledge gap, we assessed the impact of inducible MCPyV T antigens on normal human fibroblasts by performing RNA sequencing. Our data uncovered changes in expression and regulation of Wnt signaling pathway members. Building on this observation, we bioinformatically evaluated various Wnt pathway perturbagens for their ability to reverse the MCC gene expression signature and identified pyrvinium pamoate, an FDA-approved anthelminthic drug known for its anti-tumor activity in other cancers. Leveraging transcriptomic, network, and molecular analyses, we found that pyrvinium targets multiple MCC vulnerabilities. Pyrvinium not only reverses the neuroendocrine features of MCC by modulating canonical and non-canonical Wnt signaling but also inhibits cancer cell growth by activating p53-mediated apoptosis, disrupting mitochondrial function, and inducing endoplasmic reticulum stress. Finally, we demonstrated that pyrvinium reduces tumor growth in an MCC mouse xenograft model. These findings offer a new understanding of the role of Wnt signaling in MCC and highlight the utility of pyrvinium as a potential treatment for MCC.

## Introduction

Merkel cell carcinoma (MCC) is a rare yet highly aggressive neuroendocrine skin cancer, displaying variable incidence rates of approximately 0.3-1.6 per million population across different geographic regions. With metastasis occurring in more than one third of MCC patients, the disease’s morbidity rate is alarmingly high, resulting in an estimated 5-year overall survival rate of 51% for local, 35% for nodal involvement and 14% for metastatic MCC (1–5). The current standard of care for MCC involves surgical intervention followed by adjuvant radiation therapy to target the primary tumor or the draining lymph node basin (1). Historically, chemotherapeutic regimens employing combinations of platinum with drugs like etoposide, taxanes, and anthracyclines have been utilized for metastatic MCC cases not amenable to surgery. However, the response rates to chemotherapy in metastasis MCC ranged from 20-61%, and progression-free survival after chemotherapy was disappointingly limited (1, 6). In 2016, immunotherapy marked a significant advancement in MCC treatment with the introduction of PD1/PDL1 immune-checkpoint inhibitors (ICI), showing efficacy in some MCC patients. Nevertheless, the responsiveness to ICI therapy remains limited to ∼50% of patients and many responding patients become resistant to continued therapy (7, 8). Hence, the clinical landscape for MCC still lacks effective and broadly applicable therapeutic agents. Addressing this knowledge gap and identifying alternative treatment options with improved efficacy is of utmost importance in the pursuit of better outcomes for MCC patients.

The search for effective drugs targeting vulnerabilities in MCC requires a comprehensive understanding of its development. MCC can arise from either Merkel cell polyomavirus (MCPyV) infection or ultraviolet exposure (UV) or both, with MCPyV being present in approximately 80% of MCC (9, 10). MCPyV-positive (MCCP) and MCPyV-negative tumors (MCCN) exhibit distinct genetic profiles. MCCN is characterized by a high tumor mutational burden (TMB) with recurrent mutations in *TP53* and *RB1*, whereas MCCP shows a low TMB and lacks hallmark mutations (11–16). Both subtypes share common overexpressed surface markers associated with normal Merkel cells and neuroendocrine tumors (NET) (17–19). Merkel cells are not thought to be the cell of origin of MCC; instead, B cells, dermal fibroblasts, keratinocytes, and neural progenitors have all been proposed as candidates, with dermal fibroblasts being the only cell type demonstrated to support MCPyV viral replication (17). Based on previous studies focusing on the function of the two MCPyV antigens, investigators have identified multiple targeted therapies, including but not limited to: activating wild-type p53 (9, 10), specifically in the context of MCCP (18); inhibiting LSD1, as the complex formed by MCPyV small T antigen with MYCL and EP400 activates the expression of LSD1 (19, 20); inhibiting survivin (21, 22), which is derepressed due to the sequestration of RB1 by large T antigen; and inhibiting EZH2, a histone-lysine N-methyltransferase linked to tumorigenesis via epigenetic silencing of tumor suppressor genes (23–25). However, these drugs have not yet been approved for MCC patients and there is still a need for more effective targeted therapies in this aggressive cancer type.

The Wnt signaling pathway is an intricate network of protein interactions, primarily associated with embryonic development, cell morphogenesis and proliferation (26, 27). Canonical Wnt signaling and nuclear β-catenin activation are well-known to be associated with tumorigenesis, especially in the context of colon cancer and melanomas (28, 29). Non-canonical Wnt signaling is known to induce terminal neuron differentiation (30, 31). One of its ligands, WNT5A, is notably associated with increased cell motility and invasion, and its expression correlates with higher tumor grade in melanoma (32). In MCC, on the other hand, although previous genomic studies have revealed alterations in Wnt pathway members(33, 34), β-catenin remains localized to the membrane rather than the nucleus, and WNT5A is not expressed (35–37). The functional role of Wnt signaling in MCC therefore remains unclear.

In this study, we perform a wide range of bioinformatic and experimental analyses to identify members of the canonical and non-canonical Wnt signaling pathways involved in MCC development. We then characterize the effect of pyrvinium pamoate, an FDA-approved Wnt inhibitor that has shown anti-tumor potential in other cancer types, including pancreatic cancer (38, 39), colorectal cancer (40–44), breast cancer (45, 46), acute myeloid leukemia (46–48), and glioblastoma (48, 49). The multifaceted mechanism of action (MOA) of pyrvinium includes, but is not limited to, inhibiting the canonical Wnt pathway (39, 40, 42–44), disrupting mitochondrial function (50, 51), activating the unfolded protein response (UPR) (50, 52), inhibiting tumor stemness (46–48) and impeding PD1/PDL-1 interaction (53). Here, we demonstrate that MCC is sensitive to pyrvinium and explore the impact of pyrvinium’s diverse range of MOAs on the tumorigenic features of MCC.

## Results

### Wnt signaling genes are perturbed by MCPyV and in MCC tumors

In an effort to gain deeper insight into the dynamic regulatory events driving MCC tumorigenesis, we established a time-series cell line model using IMR90 normal human embryonic lung fibroblasts. These cells were transduced with a lentivirus containing the MCPyV L21 early region (ER), which codes for both the small (ST) and truncated large T (LT) antigens, or GFP (as a control), under the control of a doxycycline-inducible promoter. To characterize the host transcriptional response to MCPyV-ER, we performed RNA sequencing in triplicate at ten time points from 0 to 48 hours (Supplemental Figure 1A, Supplemental Spreadsheet 1). The raw reads were aligned to a concatenated human and viral genome to confirm that ST and LT were expressed in IMR90-ER samples in a time-dependent manner (Supplemental Table 1). Principal component analysis (PCA) revealed a trajectory representing a dynamic host transcriptome change induced by MCPyV-ER (Figure 1A). We then examined the genes that were differentially expressed between the ER and GFP samples at 48 hours (Supplemental Spreadsheet 2). As expected, the most significantly enriched GO terms were related to DNA replication and cell cycle, consistent with LT binding and inactivating Rb. In addition, we found enrichment for a wide range of other pathways (Supplemental Spreadsheet 3), including “regulation of neurogenesis” (GO:0050767; *P*_adj_ = 2.01 x 10^-2^; *OR* = 1.22) and “axonogenesis” (GO:0007409; *P*_adj_ = 8.81 x 10^-4^; *OR* = 1.28). Examining the genes annotated to these neuronal GO terms, we found that some MCC marker genes, including *ENO2, NEFM, NEFH* and *HES6,* were increased by ER (Supplemental Figure 1, B-E; Supplemental Spreadsheet 2). However, we did not observe changes in the MCC markers *CHGA, ATOH1, SOX2* and *INSM1*, likely due to cell-type-specific gene regulation in IMR90 cells.

**Figure 1.**
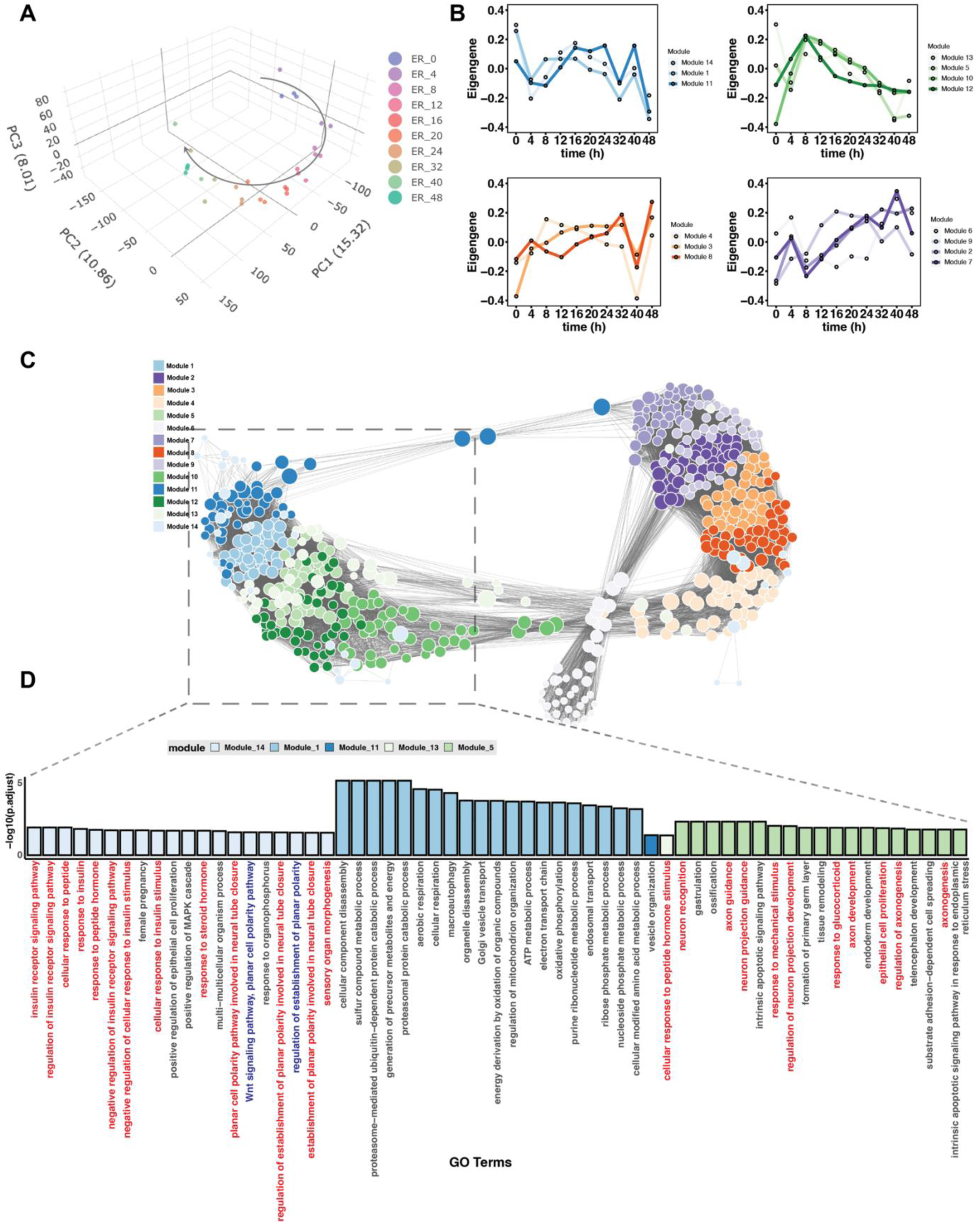
MCPyV-perturbed cell model reveals signaling pathways altered during MCC development. IMR90 normal human fibroblasts expressing inducible MCPyV early region (ER) were subjected to bulk RNA-sequencing. (**A**) Principal component analysis (PCA) performed on all 13870 expressed genes in the time series RNA-seq data. (**B**) The eigengenes of the 14 WGCNA modules were projected onto each time point and the modules were grouped by their dynamic patterns using hierarchical clustering. (**C**) Force-directed network of hub genes in the 14 WGCNA modules. The attraction forces between nodes were defined by the topological overlap matrix and were inversely proportional to the length of edges in the graph. (**D**) GO term enrichment analysis of each WGCNA gene module. The terms are ranked by adjusted p-value, and the top-ranked terms are shown. Neuroendocrine related terms are highlighted in red, Wnt signaling related terms are highlighted in blue.

To identify functional modules of host genes perturbed by ER, we conducted Weighted Gene Co-expression Network Analysis (WGCNA) on the IMR90-ER data across all time points. This analysis led to the identification of 14 modules which we further characterized based on their dynamic patterns (Figure 1B) and module eigengenes (MEs) (Supplemental Figure 2A). We visualized the top hub genes within each module and the interrelationships between the modules using a force-directed layout that places two genes closer together if they have a stronger correlation (Figure 1C). To gain insight into the biological functions associated with each module, we performed Gene Ontology (GO) enrichment analysis. The network visually separated into two components; the blue (Module 1, 11, 14) and green (Module 5, 10, 12, 13) modules clustered together on the left side, whereas the purple and orange modules clustered together on the right side and were enriched in cell cycle and metabolism pathways (Supplemental Figure 3). Among the blue modules, we observed that Module 14 genes were strongly enriched for the Wnt signaling pathway and steroid hormone response, Module 1 was enriched for cellular respiration, and Module 11 was enriched for vesicle organization, whereas the green modules were enriched for neuronal pathways (e.g. “axon development” and “neuron projection guidance” in Module 5). The eigengene analysis (Figure 1B) and module-trait (with time being considered the trait) relationship analysis (Supplemental Figure 2B) indicated that the gene expression signatures of the blue and green modules were initially distinct from each other but gradually converged over time.

To determine how host genes were being transcriptionally regulated in the presence of ER, we inferred the transcriptional regulatory networks active in both IMR90-GFP and IMR90-ER cells by integrating RNA-seq data, protein-protein interactions (PPI) and transcription factor (TF)-motif information using PANDA (54) and LIONESS (55) in five distinct time periods between zero to 48 hours. We then employed ALPACA (56) to identify differentially regulated gene sets (or “communities”) that best distinguish the IMR90-ER and IMR90-GFP networks during each of these five time periods (Figure 2A). Alongside capturing general biological processes like cell cycle, nucleic acid synthesis, protein synthesis, metabolic pathways, and histone modification, which are typically regulated during the transformation process, we also found that ALPACA community 1, the largest differential gene community between IMR90-ER and IMR90-GFP at later time periods, was enriched in Wnt signaling (*P*_adj_ = 8.79×10^-5^, *OR* = 1.70) and embryonic development (*P*_adj_ = 7.10×10^-5^, *OR* = 1.74) (Figure 2B) (Supplemental Spreadsheet 4, sheet: "er_vs_gfp_t5_module_1”) The IMR90 fibroblast cell model is incomplete and does not fully reflect the true cell of origin of MCC, which remains unknown. We therefore compared our analysis of the IMR90 cell lines to transcriptomic data from MCC patient samples. We focused on genes that are significantly differentially expressed (*P*_adj j_ ≤ 0.05, |log2 fold change| ≥ 1) between MCC tumors versus normal skin samples in a previously published dataset (GSE39612). Among these tumor DEGs, the strongest enrichment was associated with the Wnt signaling pathway (*P*_adj_ = 8.24×10^-11^, *OR* = 1.67) and neural development pathways including axon development (*P*_adj_ = 9.81×10^-17^, *OR* = 2.49), regulation of neuron projection development (*P*_adj_ = 5.12×10^-15^, *OR* = 2.42), and axonogenesis (*P*_adj_ = 3.51×10^-14^, *OR* = 2.41) (Figure 3A). In a WGCNA analysis of MCC patient samples (Supplemental Figure 4, A and B), we observed that Wnt pathway genes exhibited close co-expression with genes enriched for keratinocyte and skin development pathways (Supplemental Figure 4C).

**Figure 2.**
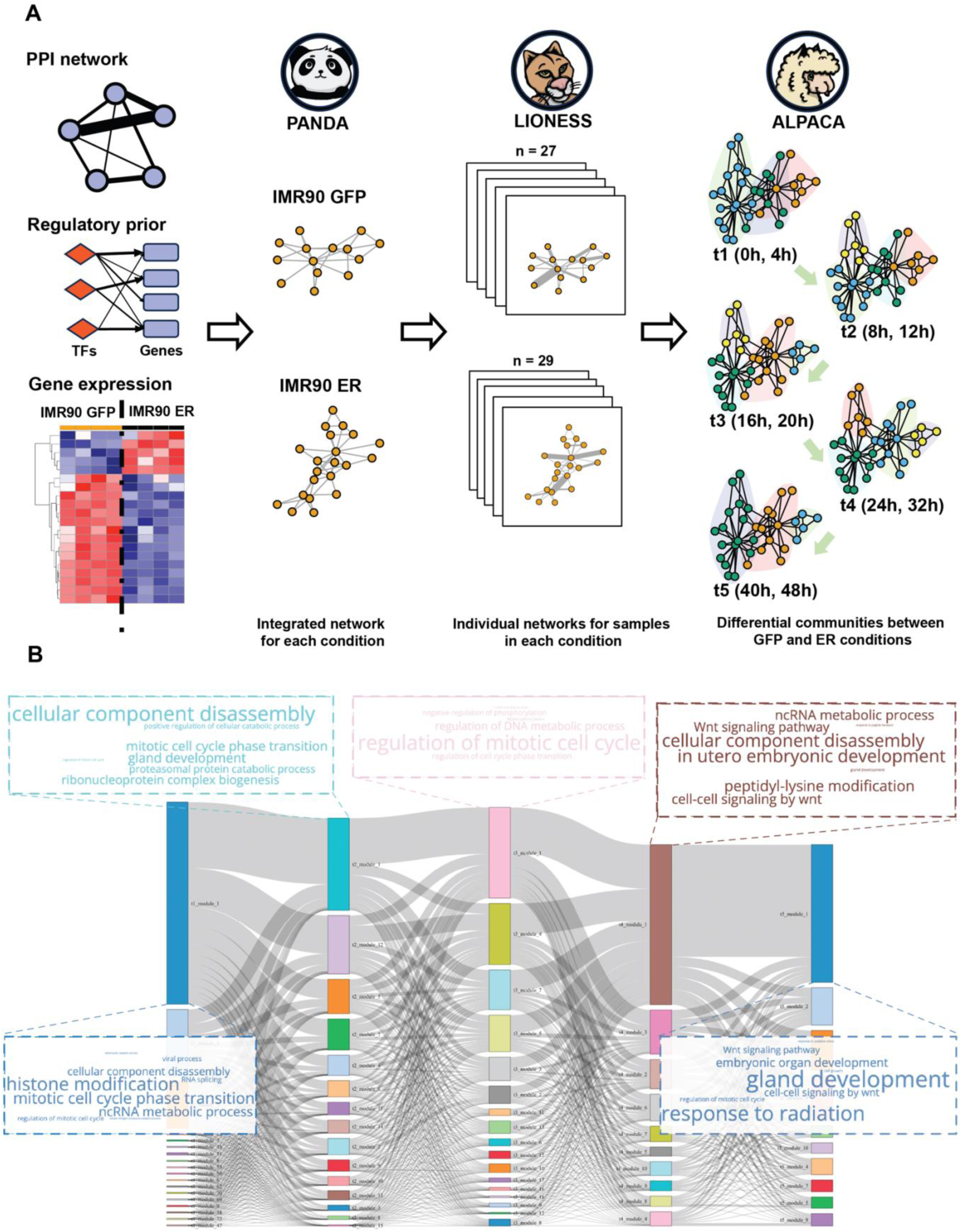
Identifying areas of active gene regulation in IMR90 cells expressing MCPyV T antigens. (**A**) Graphic workflow of regulatory network analysis. RNA-seq data was integrated with TF motif binding prior and TF protein-protein interactions to infer sample-specific regulatory networks using PANDA and LIONESS. IMR90-ER networks were grouped into five time periods and compared with IMR90-GFP networks from the same time period using ALPACA, to identify differential modules. (**B**) Sankey plot shows the dynamics of differential network communities detected by the workflow shown in panel A. Each vertical bar represents a differential community, with the size of the bar proportional to the number of genes in the community. Ribbons between adjacent bars represent the number of overlapping genes. Word cloud in the same color as the gene module annotates the enriched biological functions of genes inside the module (font size reflects the *P*_adj_ value).

**Figure 3.**
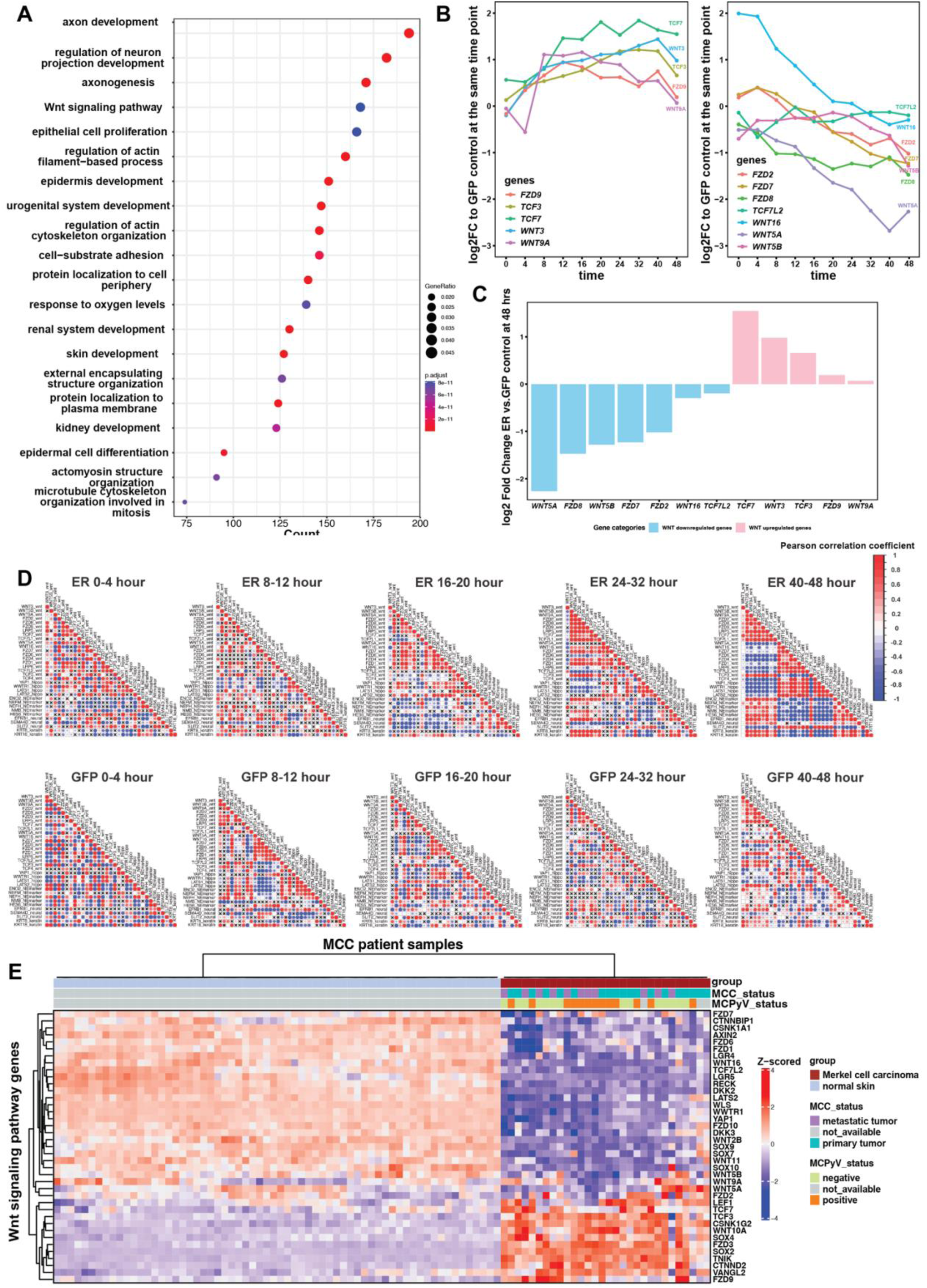
MCPyV-ER induces characteristic changes in the expression and correlation of Wnt and neuroendocrine marker genes. (**A**) Bubble plot showing GO term enrichment results for DEGs between MCC tumor samples and normal skin samples (*P*_adj_ ≤0.05 and |log2 fold change| ≥ 1). The Wnt signaling pathway ranked as one of the most significantly enriched pathways. (**B**) Log2 fold change of selected Wnt gene expression levels in IMR90-ER samples, relative to the IMR90-GFP samples at the corresponding time points. The genes were categorized into two sets based on their expression dynamics. (**C**) Bar plot showing selected Wnt gene expression levels in IMR90-ER samples relative to the IMR90-GFP samples at the 48-hour time point. The log2 fold change values were estimated from a linear model using the R package LIMMA. (**D**) The Pearson correlation coefficient was calculated to evaluate the relationship between Wnt genes and NE marker genes (*ENO2, NEFM, NEFH, NMB* and *HES6*), as well as MCC-related genes from previous studies, including hippo pathway genes (*YAP1* and *WWTR1*), neural markers (*EFNB1*, *SEMA4DM*), and keratin markers (*KRT8* and *KRT18*). The resulting correlation coefficients are color-coded according to the legend, with circle size reflecting the absolute value. Statistical significance was determined using a two-tailed t-test with a threshold of *P* < 0.05. Significant correlations are highlighted in color, while non-significant ones are crossed out. (**E**) Heatmap of Wnt signaling pathway genes in MCC tumor samples and normal skin samples. Two distinct trends in Wnt genes expression were observed, with several genes, including *FZD7, WNT16, TCF3, and TCF7,* showing trends consistent with those observed in our IMR90 model.

Next, we examined the temporal pattern of Wnt gene expression in more detail. In the IMR90 cell line model, we observed that canonical Wnt genes, such as *WNT3, TCF7,* and *TCF3,* were upregulated over time in ER samples relative to GFP control, whereas non-canonical Wnt genes, such as *WNT5A, WNT5B,* and *WNT16,* were downregulated over time (Figure 3, B and C). To validate this pattern of expression, we transduced normal human dermal fibroblasts (nHDFs) with the lentivirus containing the MCPyV L21 early region (ER) or vector control under a dox-inducible promoter and performed RNA-seq at 48 and 96 hours post-induction. Normal human dermal fibroblasts can support productive MCPyV infection and thus represent a more likely cell of origin for MCC than IMR90s (17). In the nHDF-ER cells, we again found significant enrichment for neuronal GO terms at 48 hours (e.g. GO:0010975, regulation of neuron projection development; *P*_adj_ = 1.88x10^-5^, *OR* = 1.63) and upregulation of selected MCC marker genes including *NOTCH3, EZH2, HES6, HES1,* and *NEFH* (Supplemental Spreadsheet 5 and 6). All the Wnt signaling genes that were perturbed in IMR90-ER cells, except for *TCF7*, exhibited similar trends in nHDF-ER cells, particularly the non-canonical Wnt signaling genes *WNT5A, WNT5B,* and *WNT16,* and the canonical Wnt signaling genes *WNT3* and *TCF3* (Supplemental Figure 5B). We confirmed these trends for *TCF3* and *WNT5A/B* at the protein level (Supplemental Figure 5C).

Finally, we looked more closely at the correlation between the expression of Wnt genes and a curated set of neuroendocrine (NE) related genes that were upregulated in IMR90 cells during the previously defined five time periods. IMR90-ER cells demonstrated an increased absolute Pearson correlation coefficient between the two sets of genes, whereas IMR90-GFP controls did not show any change in correlation, indicating that the Wnt signaling pathway is co-regulated with MCC and NE markers specifically in the presence of ER. In particular, canonical Wnt genes such as *WNT3, TCF7* and *TCF3* were upregulated over time and positively correlated with NE or MCC markers, whereas non-canonical Wnt genes like *WNT5A* and *WNT16* showed strong negative correlation with MCC markers (Figure 3D). The same pattern of upregulation of canonical Wnt genes and downregulation of non-canonical Wnts was found in the MCC tumor dataset (Figure 3E). Therefore, we hypothesized that MCC development and neuroendocrine differentiation requires suppression of non-canonical Wnt signaling and activation of canonical Wnt signaling. Targeting the Wnt pathway could potentially reverse the neuroendocrine features of MCC.

### Pyrvinium pamoate effectively reverses MCC signature compared to other Wnt perturbagens

We next set out to identify small-molecule perturbagens that could target the Wnt signaling pathway and reverse the gene expression changes observed in MCC. To do this, we leveraged the LINCS L1000 data to examine the impact of various Wnt signaling perturbagens on reversing the gene expression signature of MCC (Figure 4A). We focused on the drugs pyrvinium pamoate, XAV-939, IWR-1-ENDO, mesalazine, PRI-724, and indirubin, as these were the only compounds annotated by LINCS to perturb the Wnt pathway that were also commercially available. Utilizing the MCC patient samples in the GSE39612 dataset, we generated a set of MCC signature genes by computing the average Pearson correlation coefficient between the expression of each gene with a set of known MCC marker genes: *ENO2, NEFM, NEFH, NMB, HES6, SOX2, ATOH1* and *CHGA.* We selected the top 500 genes with the highest and lowest correlation scores which we call the MCC1000 signature (Supplemental Spreadsheet 7). By comparing the MCC1000 with the top 1000 differentially expressed genes from each drug perturbation, we discovered that pyrvinium (*P*_adj_ = 8.9 x 10^-9^, *OR* = 1.97), a CK1α activator which promotes the phosphorylation and degradation of β-catenin by proteosomes (44), exhibited the highest efficiency in reversing MCC1000 expression when compared to other Wnt perturbagens (Figure 4B, Supplemental Figure 6, A-F). To validate our in-silico findings, we treated MCC cell lines with pyrvinium pamoate. Pyrvinium effectively inhibited the growth of MCC cell lines (Figure 4C), even outperforming the previously reported effective drug CHIR99021, a GSK3β inhibitor sharing the same MOA as indirubin in the L1000 dataset (Figure 4D). The MTT cell growth assay demonstrated that pyrvinium significantly hindered MCC growth at concentrations as low as 100 nM. Immunofluorescence revealed that pyrvinium significantly reduced the expression of Ki67, a nuclear proliferation marker (Figure 4, E and F). Furthermore, flow cytometry analysis showed that pyrvinium can induce cell apoptosis in a dose-and time-dependent manner in four different MCC cell lines (Figure 4, G and H, Supplemental Figure 7).

**Figure 4.**
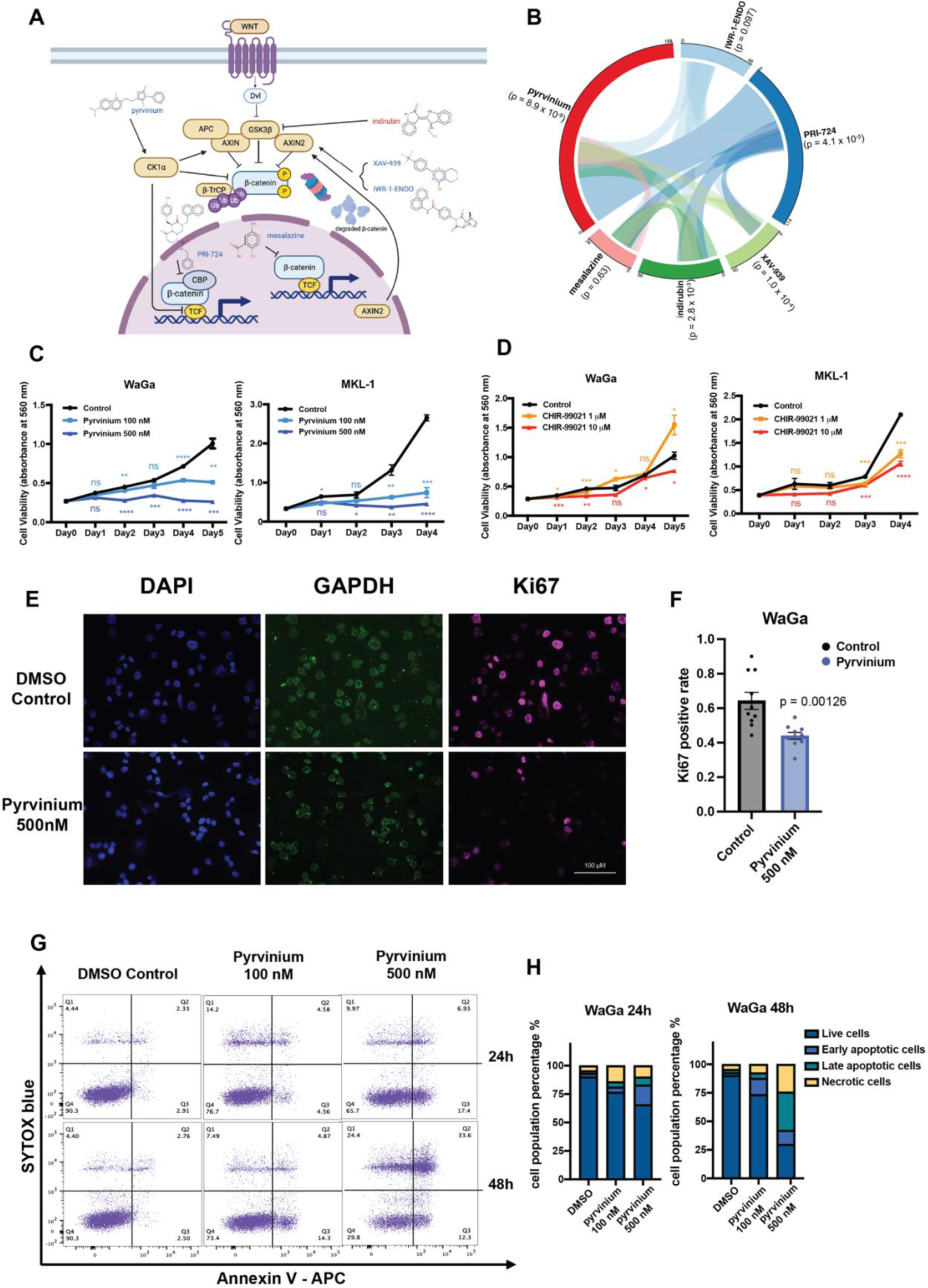
Characterization of pyrvinium pamoate as an effective perturbagen against MCC. (**A**) Simplified diagram of commercially available Wnt signaling perturbagens in LINCS L1000 dataset and their reported MOAs. (**B**) Circos plot showing pairwise comparison between MCC signature genes (MCC1000) and top drug-perturbed genes (ranked by z-scored log2[fold change]) under different drug treatments. Statistical significance was determined by Fisher’s exact test. (**C-D**) Cell proliferation assay in WaGa and MKL-1 cells, under treatment with pyrvinium, CHIR-99021 and DMSO control. Statistical significance was determined by unpaired two sample t-test (****, *P* < 0.0001; ***, *P* < 0.001; **, *P* < 0.01; *, *P* < 0.05). (**E**) Representative immunofluorescence (IF) images of WaGa cells treated with 500 nM pyrvinium and DMSO, with Ki67 staining as a proliferation marker, GAPDH as internal control, and DAPI for nucleus. (**F**) Bar graph showing the quantification of the Ki67 positive cell count relative to the total nucleus count in IF images of WaGa cells treated with 500 nM pyrvinium or DMSO. Data are presented as mean ± SEM (n = 10, *P* = 0.00126). Statistical significance was determined by unpaired two sample t-test. (**G**) Flow cytometry analysis showing the levels of Annexin V-APC and SYTOX blue staining to assess apoptotic populations in WaGa cells treated with varying doses and durations. (**H**) Bar graphs showing the quantification of different populations in pyrvinium-treated WaGa cells at 24h and 48h separately.

### Pyrvinium pamoate reveals role of Wnt signaling in maintaining neuroendocrine state

According to multiple studies on pyrvinium’s potential as an anti-tumor agent, its main mechanisms of action (MOAs) include the inhibition of canonical Wnt signaling, mitochondrial inhibition, and the activation of unfolded protein response (38, 44, 50, 57). To gain a comprehensive understanding of pyrvinium’s effect on MCC, we conducted RNA-seq on WaGa and MKL-1 cells treated with 1μM of pyrvinium, using DMSO as vehicle control, for 6 hours and 24 hours (n = 3). In pyrvinium-treated MCC cells, we observed downregulation of MCC marker genes, such as *ATOH1, SOX2, CHGA, HES6* and *NEUROD1* (Figure 5A). To identify potential mechanisms by which pyrvinium could reverse MCC marker gene expression, we set out to predict master regulator activity in pyrvinium-treated MCC cells using the state-of-the-art TF activity prediction algorithm, VIPER, in conjunction with the human TF regulon database DoRothEA. Upon pyrvinium treatment, the predicted protein activity of several MCC master regulators, including p53, MYCN, SMADs, and SOX11, exhibited an opposite trend to that previously reported during MCC development (Figure 5B) (20, 58–62). DoRothEA is a generic TF regulon database that is agnostic to cell type. We therefore repeated the VIPER analysis using MCCP-specific regulons built using ARACNe. This analysis revealed additional MCC-specific regulators such as POU4F3, HES6 and ATOH1 whose activities were reversed by pyrvinium treatment (Supplemental Figure 8, A and B).

**Figure 5.**
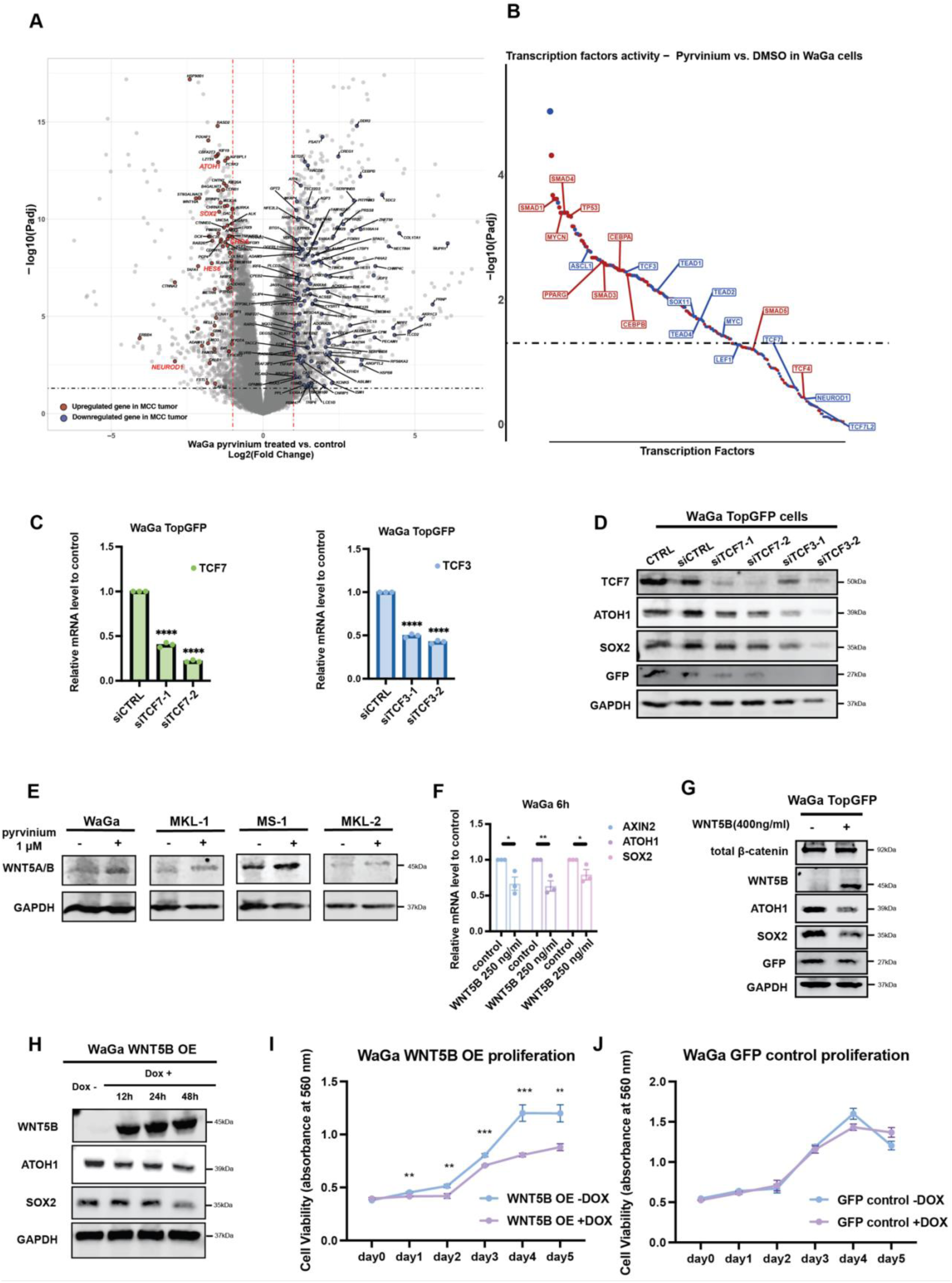
Pyrvinium pamoate reverses neuroendocrine and Wnt signaling signature in MCC cells. (**A**) Volcano plot showing differentially expressed genes (DEGs) in WaGa cells treated with pyrvinium compared to DMSO for 24 hours. DEGs with *P*_adj_ ≤ 0.05 and |log2 fold change|≥ 1 that show a reversed expression trend relative to MCC vs. normal skin are highlighted in red (upregulated in MCC) or blue (downregulated in MCC). Known MCC marker genes are labeled in red text. (**B**) Scatter plot showing the predicted activity levels of master regulators in pyrvinium-treated MCC versus DMSO-treated MCC cells. Blue (or red) indicates regulators with decreased (or increased) activity. (**C**) Relative mRNA levels of *TCF7* and *TCF3* in WaGa TopGFP cells treated with siRNA negative control, siTCF7, and siTCF3, as measured by RT-qPCR. Statistical significance determined by unpaired two sample t-test. (**D**) Protein levels of ATOH1 and SOX2, along with GFP, in untreated WaGa TopGFP cells and WaGa TopGFP cells treated with siRNA negative control, siTCF7, and siTCF3. (**E**) Protein levels of WNT5A/B following pyrvinium treatment for 24 hours in WaGa, MKL-1, MS-1 and MKL-2 cells. (**F**) Relative mRNA levels of *AXIN2, ATOH1* and *SOX2* in WaGa cells treated with recombinant human WNT5B protein for 6 hours, as measured by RT-qPCR. Statistical significance determined by unpaired two sample t-test. (**G**) Protein levels of total β-catenin, WNT5B, ATOH1, SOX2 and GFP following 6 hours of treatment with recombinant human WNT5B in WaGa TopGFP cells. (**H**) Protein levels of WNT5B, ATOH1, and SOX2 at 0, 12, 24 and 48 hours after 1 ug/ml doxycycline induction in WaGa WNT5B OE cells. (**I-J**) Cell viability in WaGa WNT5B OE cells and WaGa GFP control cells with or without 1 μg/ml doxycycline treatment. Statistical significance was determined between dox+ and dox-conditions at the same day by unpaired two sample t-test. (****, *P* < 0.0001; ***, *P* < 0.001; **, *P* < 0.01; *, *P* < 0.05).

Consistent with pyrvinium’s reported activity as a canonical Wnt inhibitor, we observed that TCF3, the main effector of canonical Wnt signaling, had higher expression in IMR90-ER and nHDF-ER cells, higher predicted activity in IMR90-ER cells, and lower predicted activity after pyrvinium treatment in MCC cell lines. Although previous studies showed low nuclear β-catenin expression in MCC (37), TCFs can be activated independently of nuclear β-catenin (63). We therefore hypothesized that MCC development involves direct activation of TCF family members. We introduced siRNAs targeting *TCF3* and *TCF7* into WaGa cells transduced with a TopGFP reporter. The TopGFP plasmid expresses GFP under the control of a 7x TCF/LEF promoter cassette, reflecting canonical Wnt signaling activity levels. After introducing the siRNAs into WaGa TopGFP cells, we confirmed that TCF/LEF activity was significantly reduced (Figure 5C). We then found that knocking down *TCF3*, and to a lesser extent *TCF7*, reduced the protein levels of neuroendocrine markers ATOH1 and SOX2 (Figure 5D; please see Supplemental Spreadsheet 8 for statistics and quantification of all Western blots). This confirms that MCC cells require TCFs to maintain neuroendocrine marker expression.

In pyrvinium-treated MCC cells, we observed not only the expected inhibition of canonical Wnt signaling (Figure 5B, Supplemental Figure 8C) but also a significant mRNA upregulation of *WNT5A* and *WNT5B*, which are Wnt ligands known to activate the non-canonical Wnt signaling pathway, which we confirmed with RT-qPCR analysis (Supplemental Figure 8, D and E). Consistent with this, Western blot results showed a modest increase in WNT5A/B levels in WaGa, MKL-1 and MS-1 cells treated with 1μM of pyrvinium (Figure 5E). WNT5A has been previously reported to promote neuron differentiation and morphological development (30, 31) and to be highly repressed in MCC tumors (35). We therefore asked whether perturbation of non-canonical Wnt alone could affect the neuroendocrine and Wnt signatures seen in MCC. Treating WaGa cells with WNT5B human recombinant protein resulted in decreased transcription of master neural development regulators, *ATOH1* and *SOX2*, as well as reduced expression of the canonical Wnt target gene *AXIN2* (Figure 5F), as evident in our RT-qPCR results. Furthermore, our Western Blot results showed a modest decrease in ATOH1, SOX2 and GFP levels upon WNT5B recombinant protein treatment (Figure 5G, Supplemental Figure 8F). However, β-catenin levels remained unchanged following WNT5B recombinant protein treatment. This suggests that WNT5B inhibits MCC regulators and canonical Wnt activity through a β-catenin-independent mechanism. To confirm the effect of WNT5B on MCC, we established a doxycycline-inducible WNT5B overexpression (WNT5B OE) WaGa cell line. Overexpression of WNT5B resulted in decreased protein levels of ATOH1 and SOX2, consistent with the recombinant protein treatment results (Figure 5H). A cell growth assay using WaGa WNT5B OE and WaGa GFP control cells suggested that the decrease in ATOH1 and SOX2 protein levels following WNT5B overexpression may be accompanied by inhibitory effects on MCC cell growth (Figure 5I and J). In summary, the neuroendocrine features of MCC tumors can be inhibited by downregulating canonical Wnt or upregulating non-canonical Wnt, both of which can be enabled by pyrvinium. This is one mechanism by which pyrvinium may reverse the process of MCC carcinogenesis.

### Transcriptome analyses reveal other mechanisms of action of pyrvinium

To characterize the genome-wide impact of pyrvinium on MCC cells, we performed GO term enrichment on all the differentially expressed genes (DEGs) following pyrvinium treatment. The most enriched GO terms were strongly associated with the “intrinsic apoptotic signaling pathway” (GO: 0097193, MKL1-24h, *P*_adj_ = 4.24 × 10^-5^), “oxidative phosphorylation” (GO: 000619, MKL1-24h: *P*_adj_ = 1.23 × 10^-2^), and “axon guidance” (GO: 0007411, MKL1-24h: *P*_adj_ = 7.98 × 10^-4^), among others (Figure 6A). To further integrate the direction of fold change and their linkages to enriched pathways, we used all the DEGs from 24 hours after pyrvinium treatment, divided them into up-(log2 fold change > 1) and down-(log2 fold change < -1) regulated clusters, performed KEGG pathway overrepresentation analysis, and constructed a gene-biological concepts network for each regulation direction cluster (Figure 6B). This network highlighted the “p53 signaling pathway” (hsa04115, MKL1-up: *P*_adj_ = 1.59 × 10^−6^, WaGa-up: *P*_adj_ = 1.42 × 10^−5^) as the top activated pathway and “oxidative phosphorylation” (hsa00190, MKL1-down: *P*_adj_ = 1.47 × 10^−5^, WaGa-down: *P*_adj_ = 1.19 × 10^−8^) as the top inhibited pathway in both cell lines. Interestingly, the network also highlighted “Small cell lung cancer” (hsa05222, MKL-1-up: *P*_adj_ = 9.7 × 10^−3^) and “human papillomavirus infection” (hsa05165, MKL-1-up: *P*_adj_ = 9.7 × 10^−3^), the biological characteristics of which are similar to MCC. These results suggested that pyrvinium’s effects are specific to neuroendocrine cancer and tumor viruses. Our analysis also revealed strong activation of endoplasmic reticulum (ER) stress by pyrvinium treatment in MCC cell lines (Supplemental Figure 9).

**Figure 6.**
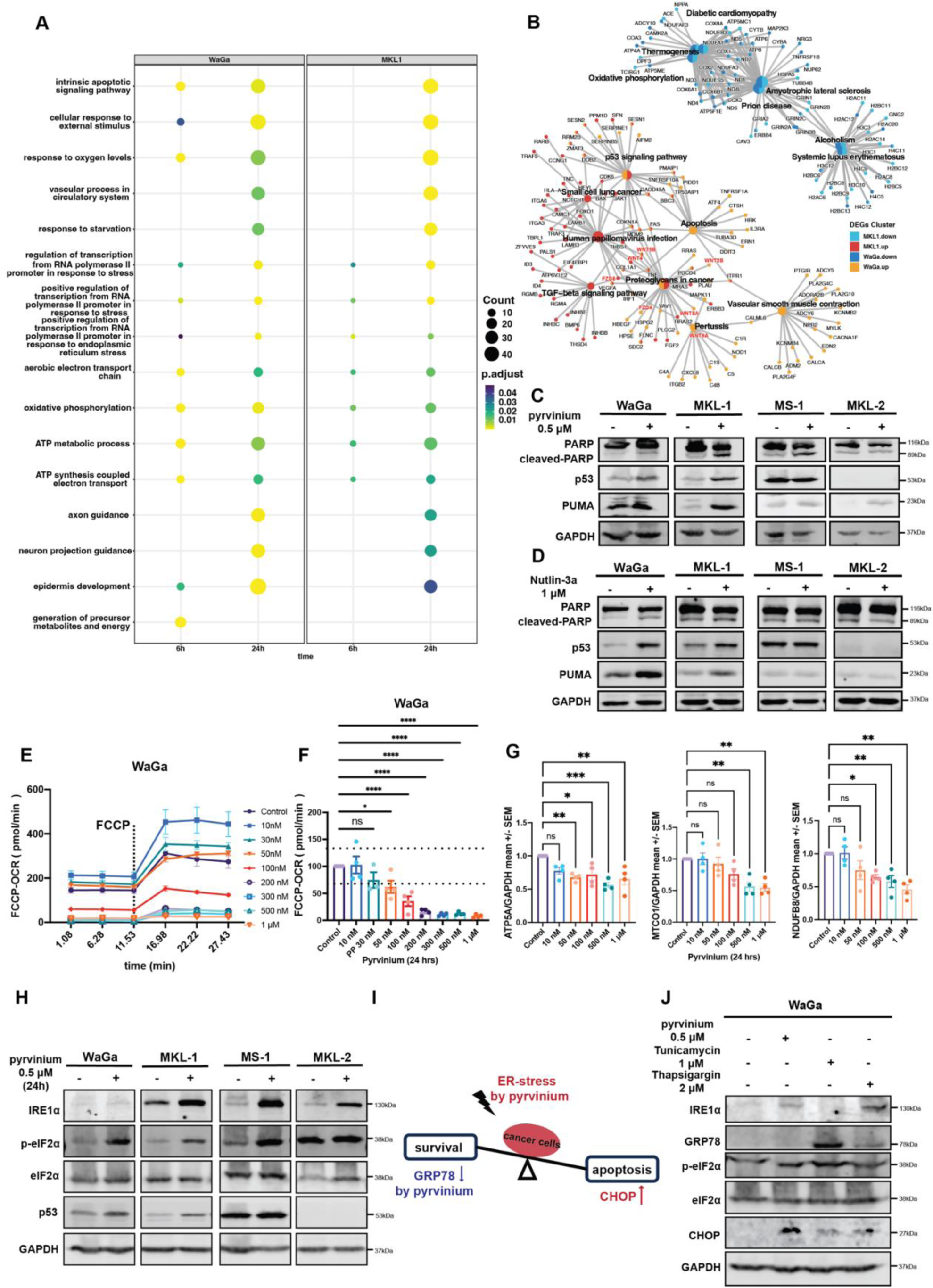
Pyrvinium targets multiple vulnerabilities of MCC. (**A**) GO term enrichment for 1 μM pyrvinium-treated WaGa and MKL1 cells at 6 and 24 hours. Size of dot indicates number of genes annotated to the GO term, and color reflects the adjusted p-value from the hypergeometric test. GO terms are primarily ranked by significance in the 24-hour MKL1 analysis. (**B**) Gene-biological concepts network showing KEGG pathway enrichment for significantly upregulated (*P*_adj_ ≤ 0.05, log2 fold change ≥ 1) and downregulated (*P*_adj_ ≤ 0.05 and log2 fold change ≤ -1) DEGs in WaGa and MKL-1 cell lines. (**C-D**) Protein levels of p53, cleaved-PARP, and PUMA in *TP53* wild type cell lines (WaGa, MKL-1) and *TP53*^Mut^/*TP53*^-/-^ cell lines (MS-1, MKL-2) 24 hours after treatment with 0.5 μM pyrvinium and 1 μM Nutlin-3a. (**E**) Representative data displayed as a line chart showing basal respiration level and maximal respiration capacity (after FCCP injection) at each time point (means ± SEM, n=4). (**F**) Seahorse OCR analysis measuring uncoupled OCR in WaGa cells treated with different doses of pyrvinium for 24 hours, compared with DMSO treated cells. Statistical significance determined by Kruskal-Wallis H Test followed by Dunnett’s post hoc test. (**G**) Quantification of OXPHOS protein levels from Western blot (Supplementary Figure 10B). All values are presented relative to mean GAPDH expression in vehicle control samples (means ± SEM, n = 4 replicate blots). Statistical significance determined by ANOVA followed by Dunnett’s multiple comparison test. (****, P < 0.0001; ***, P < 0.001; **, P < 0.01; *, P < 0.05). (**H**) Protein levels of ER stress markers in MCC cells treated with pyrvinium for 6 and 24 hours. (**I**) A graphic model illustrating pyrvinium’s effect on the balance between ER stress and UPR signaling. (**J**) Proteins levels of Unfolded Protein Response (UPR) markers in WaGa cells after 24 hours of treatment with pyrvinium or other ER stress inducers.

### Pyrvinium pamoate induces MCC cell apoptosis through p53-dependent and -independent mechanisms

p53 functions as an essential tumor suppressor, and loss of function mutations in the *TP53* gene are found in approximately 50% of all cancers (64). However, p53 inactivation mutations are less frequently (13%-28%) reported in MCC (11, 12, 65). To determine if the pro-apoptotic effect of pyrvinium is dependent on wild-type p53, we used a panel of four MCC cell lines: WaGa, MKL-1, MS-1 and MKL-2. WaGa and MKL-1 cell lines are p53 wild-type cells; MS-1 harbors a *TP53* deletion mutation, resulting in an inactive p53 protein lacking amino acids 251-253; in MKL-2, p53 protein is undetectable due to post-transcriptional repression (66). Our Western blot results showed significant increases in p53, cleaved PARP, and PUMA protein levels in pyrvinium treated WaGa and MKL-1 cells. In contrast, there was no change in MS-1 and MKL-2 cells, except for MKL-2 which exhibited an increase in PUMA (Figure 6C). We further compared the effect of pyrvinium in all four cell lines with the MDM2 inhibitor Nutlin-3a. Nutlin-3a increased p53, cleaved-PARP and PUMA protein levels in WaGa and MKL-1 cells, but with lower efficacy than pyrvinium. Nutlin-3a did not increase the levels of cleaved-PARP and PUMA in MS-1 and MKL-2 (Figure 6D). MTT assay results revealed that at the 48-hour time point, the IC_50_ values for pyrvinium in WaGa (0.4217 μM), MKL-1 (0.1104 μM), MS-1 (0.3359 μM) and MKL-2 (0.4362 μM) cells were all in the 100 nanomolar range. In contrast, Nutlin-3a was only effective in WaGa (1.290 μM) and MKL-1 (0.1441 μM) cells and showed no inhibitory effect in MS-1 and MKL-2 even at 10 μM (Supplemental Figure 10A). Our results suggest that, although p53 activation plays a significant role in cell apoptosis during pyrvinium treatment, there are also p53-independent pro-apoptotic mechanisms that provide pyrvinium with an advantage as a novel therapy for treating *TP53^mut^*or *TP53^-/-^* MCC.

Pyrvinium has been previously identified as a mitochondrial inhibitor and has demonstrated even greater potency in nutrient-deficient conditions (38, 39, 67). To assess the extent of oxidative phosphorylation inhibition by pyrvinium in MCC cells, we conducted an FCCP-OCR analysis using the Seahorse XF96 Analyzer. Employing dose-dependent treatments of pyrvinium over 24 hours, we evaluated the FCCP-OCR response. Remarkably, we observed a significant decrease in FCCP-uncoupled OCR (a measurement of maximal ETC activity), indicative of electron transport chain impairment, starting from around 50-100 nM of pyrvinium (Figure 6, E and F). Additionally, Western blots performed using the same samples as the Seahorse experiment revealed that pyrvinium treatment reduced the protein levels of multiple components in mitochondrial complexes (NDUFB8 subunit of Complex I, MTCO1 subunit of Complex IV, ATP5A subunit of Complex V, n = 4) (Figure 6G, Supplemental Figure 10, B-D). The combined results from OCR and Western blotting indicates that pyrvinium impairs mitochondrial function by inhibiting the expression of mitochondrial complex subunits. Moreover, pyrvinium treatment downregulated numerous mitochondrial protein-coding genes (Supplemental Figure 10E), indicating that the decrease in mitochondrial complex proteins could also arise from direct suppression of mitochondrial gene transcription.

A potential downstream effect of mitochondrial dysfunction is endoplasmic reticulum (ER) stress. The ER is tightly associated with mitochondria by multiple contact sites and forms special domains called mitochondria-ER associated membranes (MAMs). ER stress can be triggered by various intra-or extracellular factors, such as glucose starvation, hypoxia, Ca^2+^ depletion and protein misfolding. In our RNA-seq data, we observed significant enrichment and upregulation of ER stress signaling, which could lead to cell apoptosis. Western blot analysis demonstrated that pyrvinium treatment elevated the activity of ER stress sensors residing in the ER membrane, including increased IRE1α protein levels and increased phosphorylation of eIF2α by PERK in all MCC cell lines, regardless of their p53 status (Figure 6H). The master regulator of unfolded protein response (UPR) signaling, GRP78 (BiP), can mitigate ER stress by arresting transient transcription, degrading ER associated proteins and inducing ER chaperones to support cell survival. However, in cases of severe ER stress, apoptosis is triggered by the effector protein CHOP (52). To investigate changes in ER stress and UPR levels with varying pyrvinium dosages, we performed Western blot analysis on WaGa cells treated with pyrvinium for 24 hours at different concentrations. The results indicated that pyrvinium increased ER stress and reduced GRP78 levels in a dose-dependent manner, while CHOP levels increased promptly when treated with 500 nM of pyrvinium (Supplemental Figure 10F). Overall, these results suggest that pyrvinium treatment elevates ER stress levels and impairs the unfolded protein response, leading to an amplification of ER stress and induction of cell apoptosis (Figure 6I). Furthermore, we compared the ER-stress inducing capacity of pyrvinium with known ER inducers, namely tunicamycin (TM), an N-glycans blocker that induces unfolded protein generation, and thapsigargin (TG), an inhibitor of sarco-endoplasmic reticulum Ca^2+^ ATPase (SERCA) that prevents Ca^2+^ flow from the cytoplasm into the ER. We found that 500 nM pyrvinium induced CHOP in MCC cells to a stronger extent than thapasigargin and tunicamycin applied at higher concentrations (Figure 6J). In conclusion, pyrvinium efficiently enhances ER stress in all MCC cells, potentially contributing to cell death.

### Pyrvinium pamoate effectively inhibits tumor growth in xenograft mouse model

To assess the effect of pyrvinium on MCC cells in vivo, we performed a xenograft study with MKL-1 cells in NSG mice (Figure 7A). The administration of gradually increasing doses of pyrvinium, from 0.1 mg/kg to 1.0 mg/kg daily via intraperitoneal injection, was enough to cause tumor growth inhibition; the treated mice exhibited significantly slower tumor growth across time than the control mice (n = 7, *P* < 0.001) (Figure 7B, Supplemental Figure 11, A and B), with control mice exhibiting a slope of 1.34 (SE = 0.0823), while treated mice exhibited a slope of 0.937 (SE = 0.0861). Two other independent studies, in which we administered higher doses of pyrvinium from 0.6 mg/kg to 1.0 mg/kg of pyrvinium daily (n = 4, *P* < 0.001, control slope: 2.61, treated slope: 1.43), or 1.0 mg/kg three times a week (n = 10, *P* < 0.001, control slope: 2.24, treated slope: 1.72) (Supplemental Figure 12, A-E, Supplemental Table 2 -4), both demonstrated even larger reductions in tumor burden. Among these three independent studies, the gradual dose escalation strategy was optimally tolerated by NSG mice and resulted in minimal side effects. To test whether pyrvinium acts through the pathways identified in vitro, we performed H&E and IHC staining for ATOH1 and Ki67 on the xenograft tumors from the control and treatment groups (Figure 7C). We observed that pyrvinium-treated MCC tumors exhibited a modest decrease in ATOH1 expression, coupled with decreased levels of Ki67, thus demonstrating reduced proliferation in drug-treated tumors (Figure 7D). These in vivo results align with our in silico and in vitro findings.

**Figure 7.**
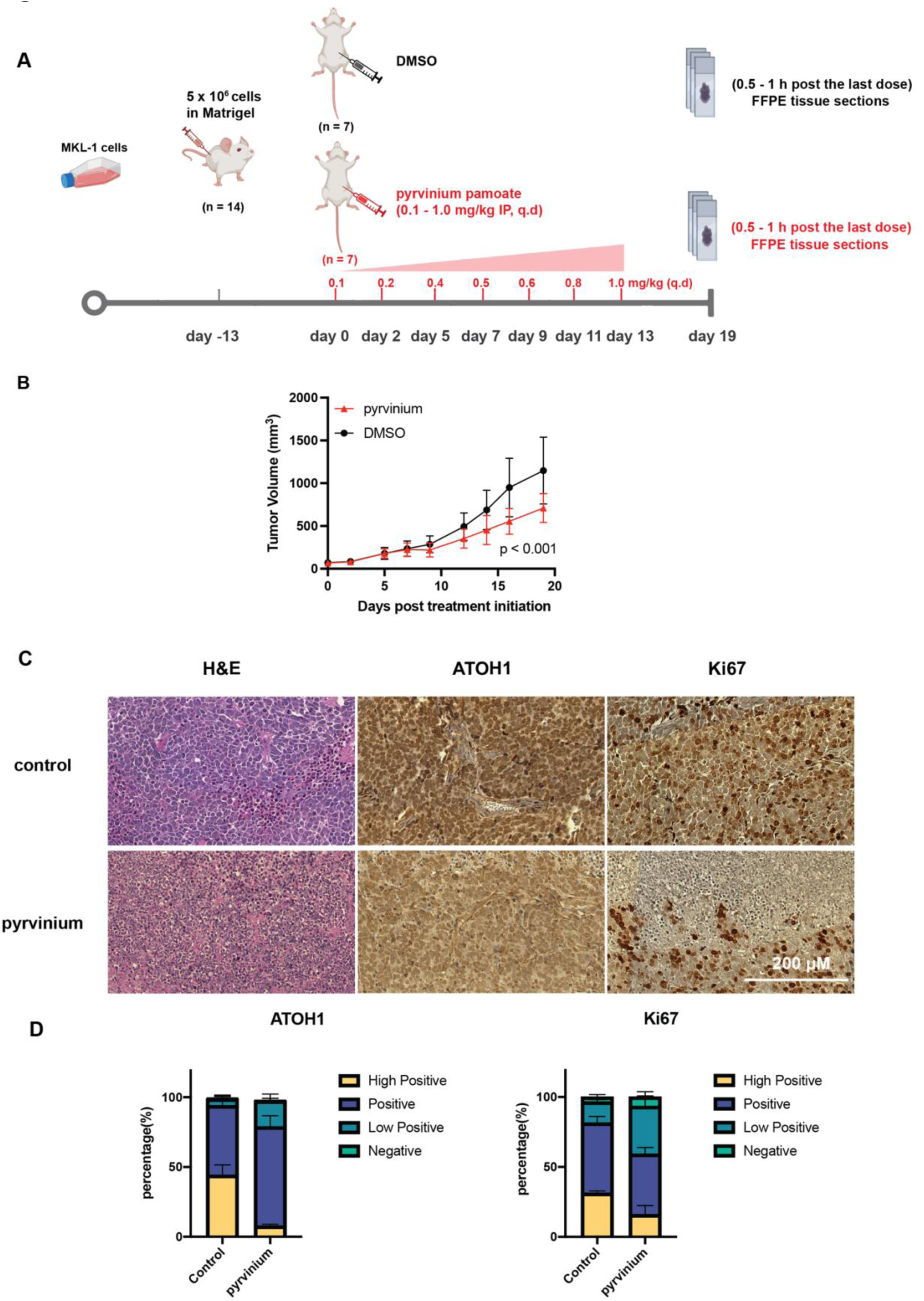
Antitumor activity of pyrvinium in an MKL-1 xenograft tumor model. (**A**) Experimental design of the in vivo study. (**B**) Tumor growth curve showing the mean tumor volume of vehicle control and pyrvinium treated mice from day 0 to day 20 of treatment (means ± SEM, n = 7). (**C**) H&E and IHC staining results on serial sectioning slides for each marker in the same lesion. Tumor tissues were collected 0.5 to 1 hour after the final 1 mg/kg dose of pyrvinium in in vivo Study #2 (experimental design shown in Supplemental Figure 12C). (**D**) Percentage of MKL-1 xenograft tumor tissue expressing ATOH1 or Ki67 in vehicle control group and pyrvinium-treated group.

## Discussion

In this study, we used an inducible cell line model to identify cellular pathways driving Merkel cell carcinoma. Leveraging genomic analyses and multiple databases, we shed light on the role of the Wnt signaling pathway and discovered that MCC is sensitive to the Wnt inhibitor pyrvinium pamoate. By studying the impact of pyrvinium on MCC cells, we discovered specific elements of canonical and non-canonical Wnt signaling that help maintain the neuroendocrine features of MCC. Pyrvinium also has anti-tumor effects through multiple mechanisms, including the activation of p53, downregulation of mitochondrial complex genes, and induction of ER stress. These insights contribute to our understanding of MCC development and offer a new avenue for targeted therapeutic strategies for this aggressive neuroendocrine malignancy.

In human skin, Wnt signaling enables intercellular communication between keratinocytes and fibroblasts to induce proliferation of dermal cells, regeneration of hair follicles (68), and Merkel cells (69). Previous research has demonstrated that, although β-catenin activity level is low in MCC tumors, canonical Wnt signaling can stimulate MCPyV infection in human dermal fibroblasts (17). It is known that many members of canonical and non-canonical Wnt signaling are expressed in the developing and mature nervous systems (70, 71). We found that aspects of both canonical and non-canonical Wnt signaling are altered in Merkel cell carcinoma. Using our IMR90 cell line model expressing MCPyV ER, we discovered that canonical Wnt signaling genes, including *WNT3, TCF7*, and *TCF3*, are associated with neuroendocrine marker expression, whereas non-canonical Wnt genes, like *WNT5A*, *WNT5B*, and *WNT16* are negatively correlated with MCC markers, consistent with previous findings of low WNT5A expression in MCC cells (35). In particular, YAP and WWTR1 are two hippo pathway regulators that have growth-suppressive properties and are silenced in many NE cancers and MCCP (72, 73). It has been reported that YAP and WWTR1 are also mediators of non-canonical Wnt signaling (73). Consistent with this hypothesis, in IMR90 ER-expressing cells, we found that *WNT5B* expression is positively correlated with the expression of *YAP* and *WWTR1* and is negatively correlated with NE markers (Figure 3B). We further observed that reducing TCF activity using siRNAs and increasing WNT5B through recombinant protein or an overexpressing vector in MCC cells exerts inhibitory effects on MCC marker genes and canonical Wnt activity. Overexpressing WNT5B over time in MCC cells inhibits cell growth. These results together suggest that the observed alterations in Wnt signaling play an important function in maintaining the neuroendocrine features of MCC. Since non-canonical Wnt signaling is known to induce terminal neuron differentiation, it is possible that Merkel cell carcinoma tumors must suppress non-canonical Wnts to remain in a proliferative progenitor-like cell state, while maintaining canonical Wnt (or TCF) activity at a level that supports proliferation. However, further experiments are needed to determine how non-canonical Wnt ligands are suppressed during MCC development.

Our IMR90 cell line model has several limitations, including the fact that IMR90s are derived from embryonic lung tissue and that we only profiled their transcriptome for 48 hours after ER expression, whereas MCPyV is a lifelong infection and MCC tumor development occurs over a timescale of years. To mitigate these issues, we confirmed the observed changes to the Wnt pathway in neonatal human dermal fibroblasts (nHDFs) and MCC tumors. Moreover, in both IMR90s and nHDFs, we observed that the response of the host transcriptome stabilizes by 48 hours after induction of MCPyV ER, with little change observed between 48 to 96 hours (60). This suggests that our experimental design sufficiently captures the immediate response of the host cell to MCPyV ER. However, expression of MCPyV ER over months or years could cause epigenetic alterations and other long-term changes that promote MCC development. We leave this question to future work.

Next, we used bioinformatic analysis of the LINCS L1000 dataset to identify pyrvinium pamoate, an FDA-approved anthelminthic drug and known inhibitor of canonical Wnt signaling, as a therapeutic candidate for MCC. Pyrvinium reduces the activity of canonical Wnt master regulators TCF3 and TCF7, and increases expression of non-canonical Wnt ligands WNT5A/B. Moreover, analysis of LINCS L1000 genetic perturbation data, MCC patient samples, and RNA-seq data from pyrvinium-treated MCC cell lines suggests that pyrvinium’s effect in MCC might involve a combined perturbation of multiple Wnt components (Supplemental Figure 8G). To further elucidate the relationship between pyrvinium, Wnt signaling, and MCC, future experiments could determine target genes of TCF3/7 using ChIP-seq or ATAC-seq, and systematically measure the activity of key components of the canonical and non-canonical Wnt pathways – such as β-catenin, CK1α, WNT5A/B and their isoforms, and NFATs – upon T antigen expression or pyrvinium treatment.

We found that pyrvinium acts through both p53-dependent and -independent pathways to inhibit MCC growth. Because of the relatively low *TP53* mutation rate in MCC, particularly in MCCP, compared to other cancer types, the ability of pyrvinium to activate p53 response could prove beneficial (2, 74). Prior studies applying the ubiquitin ligase MDM2 inhibitors alone or with MDM4 inhibitors have led to p53 activation and cell apoptosis (9, 10, 66). Through transcriptomic profiling of pyrvinium-treated MCC cells, we observed significant activation of p53 signaling and validated it at protein levels. Pyrvinium exhibited similar or even higher efficacy for p53 activation and p53-mediated cell apoptosis than the MDM2 inhibitor Nutlin-3a in *TP53*^WT^ MCC cells. Pyrvinium is known as an activator of CK1α, a serine/threonine protein kinase, the activation of which promotes the phosphorylation and degradation of β-catenin by proteasomes (44). Prior work has shown that MCPyV ST can induce the overexpression of CK1α in MCC (9). Notably, other studies determined that CK1α could phosphorylate the N-terminal phosphorylation sites of p53, especially at the serine 20 site, which is believed to attenuate interaction of p53 with MDM2 and stabilize the binding of the co-activator p300, thereby activating p53 function (75, 76). CK1α activation could be one of the mechanisms by which pyrvinium activates p53, but further experiments are required to elucidate this.

Our analyses revealed that pyrvinium efficiently inhibited oxidative phosphorylation in MCC cells, at an effective dose as low as 50 nM. Both transcriptomic and protein level assessments indicated that pyrvinium suppressed the transcription of mitochondrial DNA. This observation aligns with a previous study that demonstrated a correlation between pyrvinium efficacy and the expression of mitochondrial-related genes in other cell lines (38). A potential mechanism for mitochondrial inhibition could be pyrvinium binding to and stabilizing mitochondrial G-quadruplexes, thus disrupting mitochondrial transcription (38). We also revealed that pyrvinium induced apoptosis by enhancing ER stress and abrogating the UPR signaling by targeting GRP78. In addition to these MOAs, we also observed that pyrvinium downregulates expression of EZH2 and survivin (BIRC5), two previously reported drug targets in MCC (21, 77). Potential co-treatment with pyrvinium and EZH2 inhibitors should be tested in MCC models.

In a xenograft model, we demonstrated that pyrvinium effectively suppressed MCC tumor growth in NSG mice and concurrently reduced MCC marker genes within the xenograft tumor tissue. There remains a need for further optimizing the administration method – through either IP injection or oral delivery systems – since some of the mice in our higher-dose studies did not tolerate IP injection at 0.6-1 mg/kg dosage. Encouragingly, pyrvinium pamoate was reported safe with oral dosing with the best effects noted at 35 mg/kg daily in mice (38). Moreover, pyrvinium pamoate has received approval for a Phase 1 clinical trial aimed at treating pancreatic ductal adenocarcinoma (PDAC) (NCT05055323), showing that a safe treatment protocol is possible for human patients (78). Our in vivo data, combined with other published reports, highlights pyrvinium as a candidate for anticancer therapy in Merkel cell carcinoma.

Through a combination of genomic studies, bioinformatics, and in vitro and in vivo work, our research has shown that the Wnt signaling pathway plays a functional role in maintaining the neuroendocrine features of Merkel cell carcinoma. Furthermore, we have demonstrated the potential of pyrvinium pamoate as an anti-tumor agent that targets multiple vulnerabilities of MCC. Further studies are needed to comprehensively characterize the role of Wnt signaling on cancer hallmarks, and to optimize treatment protocols for the development of pyrvinium pamoate as a clinically useful drug for Merkel cell carcinoma.

## Methods

### Sex as a Biological Variable

Sex was not considered as a biological variable. Experiments were conducted on female and male mice in separate studies, and similar results were observed in both sexes.

### Cell Culture and Chemicals

Characteristics of IMR90 cells, nHDF cells, and Merkel cell carcinoma cell lines WaGa, MKL-1, MS-1 and MKL-2 have been previously reported (17, 20, 60). MCC cell lines WaGa, MKL-1, MS-1 and MKL-2, as well as fibroblast cell line IMR90 have been previously described (79). Fibroblast cell line nHDF was obtained from ATCC (#PCS-201-010). All the cell lines were cultured at 37°C under 5% CO2. IMR90 cells were cultured in DMEM complete medium (Corning, cat: #10-013-CV), containing 15% fetal bovine serum (FBS), 10 U/ml penicillin, and 10 mg/ml streptomycin (GIBCO, cat: #35050061). nHDF cells were cultured in DMEM complete medium, containing 10% FBS, 1% GlutaMAX, 10 U/ml penicillin, and 10 mg/ml streptomycin. WaGa, MKL-1, MS-1 and MKL-2 cells were cultured in RPMI-1640 medium (Corning, cat: #10-040-CV), containing 10% FBS, 1% GlutaMAX, 10 U/ml penicillin, and 10 mg/ml streptomycin; See Supplemental Table 5 for chemicals used in this study.

### Cell Transfection

Doxcycline-inducible IMR90 and nHDF lines expressing MCPyV ER and GFP (or vector control) were generated as previously described (60). The inducible plasmids containing MCPyV ER and GFP were generated by cloning PCR products of the MCPyV early region DNA sequence, isolated from MCC tumor sample MCCL21 (NCBI accession number: KC426955), and EGFP sequence into the pLIX_402 donor vector using the Gateway system. WaGa cell lines expressing 7xTcf-eGFP (WaGa TopGFP) and hWNT5B (WaGa WNT5B OE) were generated with Mirus Bio TransITLenti Transfection Reagent (Thermo Fisher Scientific, cat: #MIR6604), according to the manufacturer’s instruction. 7TGP reporter plasmid was a gift from Roel Nusse (Addgene plasmid #24305), and the inducible WNT5B overexpression plasmid was generated by cloning the hWNT5B[NM_030775.2] protein coding sequence into the Gateway pLIX_403 donor vector. pLIX_403 donor vector plasmid was a gift from David Root (Addgene plasmid #41395). Lentiviral packaging plasmid psPAX2 and envelope plasmid pMD2.G were gifts from Didier Trono (Addgene plasmids #12260, #12259). The siRNAs for TCF3, TCF7 and negative control (IDT, cat: #51-01-14-03) were resuspended in Nuclease-Free Duplex Buffer (IDT, cat: #11-01-03-01) to achieve a stock concentration of 100 μM. Transfection was performed using the Neon Transfection system (Thermo Fisher Scientific, cat: #MPK5000, #MPK100). For each transfection, 2 μl of siRNA was added to 100 μl of one million WaGa cells in DPBS. A single electrical pulse of 1800 mV for 20 ms was applied to introduce the siRNA into the cells. The electroporated cells were then transferred into 2 ml of fresh complete culture medium, resulting in a final siRNA concentration of 10 nM. After 48 hours of culture, the cells were collected for assays. The full sequences of all plasmid constructs, siRNAs, and primers are provided in the Supplemental Materials.

### Western Blot

Cells were seeded in 25 cm^2^ cell culture flasks and were treated with pyrvinium for 24 hours. Then, cells were washed with cold PBS and lysed with RIPA buffer (Thermo Fisher Scientific, cat: #89901) supplemented with EDTA-free protease inhibitor cocktail (Roche, cat: #04693132001). After centrifugation for 20 min at 17000 rpm at 4℃, supernatant was collected. Protein concentrations were determined using the Pierce BCA assay kit (Thermo Fisher Scientific, cat: #23227). Protein samples were denatured with 4x Laemmli Sample buffer with 10% Beta-Mercaptoethanol (Sigma-Aldrich, cat: #M6250) followed by 10 min boiling at 95℃. Then, samples were loaded and run on SDS-PAGE for 80 min at 120V. Proteins were transferred to nitrocellulose membranes at 200mA for 90 min. Blocking and incubation of primary and secondary antibody were performed under manufacturer recommended conditions. The membranes were visualized by Li-Cor Odyssey FC imager. See Supplemental Table 6 for antibodies used in this study. Every blot was repeated at least two times, and quantification and statistics are provided in Supplemental Spreadsheet 8.

### Mitochondrial Oxygen Consumption Rate (OCR) Analysis

The assay plates and cartridges were treated with 1ml of Seahorse XF96 calibrant overnight in a 37℃ non-CO2 incubator, ensuring that each well was fully immersed in liquid (Seahorse Bioscience, cat: #103680 -100). WaGa cells were washed by DPBS once and resuspended in fresh culture media to 3×10^6^ cells/ml. The cell suspension was aliquoted into different 15 ml tubes and the pyrvinium was added to final concentrations of 10, 30, 50, 100, 200, 300, 500 nM, and 1 μM. A multichannel pipette was used to transfer 50 µL of the cell suspension (1.5×10^5^ cells) evenly to each well of the PDL plates (Seahorse Bioscience, cat: #103798-100). The cells were centrifuged at 200 rcf (zero braking) for 1 minute and plates were transferred to a 37°C incubator. After 24 hours of treatment, the PDL plates were gently tapped on stacks of tissue paper to get rid of the drug containing culture media. Then, 180 µL of warm assay media (Seahorse Bioscience, cat: # 103681-100) was slowly and gently added along the side of each well after washing the cell with assay media once. The plates were placed in the incubator for 30 minutes and cells were observed under the microscope to check that cells were not detached and 95% of the well bottom area was covered by WaGa cells. Maximal respiratory capacity (FCCP-OCR) was measured using the Seahorse Bioscience XF-96 Extracellular Flux Analyzer (Seahorse Bioscience, Billerica, MA). The protocol is optimized for the given condition; basal OCR was measured three times followed by injection of 20 μl of 5 μM FCCP (Sigma-Aldrich, cat: #C2920) in port A to reach a final concentration of 0.5 μM. The stepwise settings used to measure the FCCP-OCR in the WaGa cells are shown in Supplemental Table 7.

### Quantitative real-time PCR (qPCR)

Total cellular RNA was extracted using TRIzol reagent (Invitrogen cat: #15596026) according to the manufacturer’s instructions. Then, the RNA samples were reverse transcribed into cDNA using the SuperScript IV RT-PCR kit (Thermo Fisher Scientific, cat: #12594100). Real-time PCR was then performed using the Applied Biosystems StepOne™ system with SYBR Green RT-PCR Master Mixes (Thermo Fisher Scientific, cat: #A25742). The PrimePCR SYBR green assay primers (see Supplemental Table 8) were used to amplify gene of interests and housekeeping gene. The data were acquired as a threshold cycle (Ct) value. The ΔCt values were determined by subtracting the average internal housekeeping gene Ct value from the average target gene Ct value. Since the amplification efficiency of the target genes and internal control gene was equal, the relative gene expression was calculated using the 2^-ΔΔCt^ method. Each measurement was performed in triplicate and repeated three times.

### Xenograft efficacy study

5×10^6^ MKL-1 cells with 50% Matrigel were implanted in the right flank of 8-week-old NSG mice (The Jackson Laboratory, Strain#: 005557) subcutaneously. Tumors were allowed to grow to an average size range 50 – 100 mm^3^ until randomized into 2 groups per group. Mice were treated with vehicle control (10% DMSO + 90% of 20% HP-β-CD) or pyrvinium pamoate administered intraperitoneally once daily. We used both female and male mice in three independent studies. In Study #1, male mice were treated with 1 mg/kg pyrvinium three times (MWF) a week (n = 10). In Study #2, female mice were treated at 0.6 mg/kg for 14 days, followed by 7 days at 1mg/kg dose (n = 4). In Study #3, male mice were treated with increasing doses, starting at 0.1 mg/kg on day 1, then escalating to 0.2 mg/kg on day 3, 0.4 mg/kg on day 6, 0.6 mg/kg on day 8, 0.8 mg/kg on day 11, and finally 1.0 mg/kg from day 13 until the study ended on day 19 (n = 7). All control and treated mice received subcutaneous injections of saline concurrently with each dose. For all three studies, tumor volumes and body weights were measured three times a week. Tumor samples were collected from mice reaching endpoint (tumor volume exceeding 2000 mm^3^) or on study termination on day 20. One third of each tumor sample was fixed using 10% neutral formalin and kept at room temperature for 24 hours. After 24 hours, samples were then transferred and preserved in 70% ethanol and proceeded to paraffin embedding for IHC. The rest of the tissue was snap frozen with liquid nitrogen and stored at -80℃ for further analysis. Statistical analysis was performed using a linear mixed effects model to account for the correlation in the tumor volume measurements across time within a mouse. Tumor volume was transformed and normalized by using the square root. The model incorporates the effects of time, treatment, and their interaction; the interaction tested whether the tumor volume growth rate across time (slope) differed in the pyrvinium-treated versus control mice. A p-value < 0.05 was considered statistically significant.

### RNA-seq

IMR90 cells were transduced with doxycycline-inducible lentiviral vectors containing MCPyV-ER (with truncated large T antigen) or GFP sequence (as a control). The cells were treated with doxycycline for 48 hours and harvested at 0, 4, 8, 12, 16, 20, 24, 32, 40, and 48 hours post dox treatment, triplicated for each time point. RNA was purified using the RNeasy Plus Mini Kit (QIAGEN, cat#: 74034) and mRNA was isolated with NEBNext Poly(A) mRNA Magnetic Isolation Module (New England BioLabs, cat#: E7490). Sequencing libraries were prepared with the NEBNext mRNA library Prep Master Mix Set for Illumina (New England BioLabs, cat#: E6110), passed Qubit bioanalyzer and qPCR QC analyses, and sequenced on the Illumina HiSeq 2000 platform.

WaGa and MKL1 cells were treated with dimethyl sulfoxide (DMSO) or 1 μM pyrvinium for 6h and 24h in triplicate for each condition. nHDF-ER and nHDF-Ctrl cells were treated with doxycycline for 0, 48, and 96 hours, also in triplicate for each time point. Total RNA was isolated from the cells using the RNeasy mini kit (QIAGEN, cat: #74104). The isolated RNA was subjected to quality control using an Agilent 2100 bioanalyzer, and all samples passed the quality check. The RNA samples were then subjected to library preparation and sequencing on the Illumina NovaSeq 6000 platform.

### WGCNA

Weighted gene co-expression network analysis (WGCNA) was performed using the WGCNA R package (80). A signed network was constructed on the 10513 genes differentially expressed between ER 48h and GFP 48h samples (with *P*_adj_ ≤ 0.05 and no threshold on fold change) across all time points. The soft thresholding power was set to 16 after comparing with scale-free topology. To identify modules of highly correlated genes, we applied a minimum module size of 30 and a height cutoff of 0.25 to the hierarchical clustering dendrogram. The algorithm assigned the 10513 genes to 14 signed modules. The eigengene of each module was then projected onto individual samples, and the mean of the triplicates for each time point was used for plotting. GO enrichment analysis was performed on each module to identify biological pathways enriched among the genes within the module.

We then calculated the sum of the Topological Overlap Matrix (TOM) for all edges that connect to each gene and sorted genes based on this score. The forty genes that had the highest summed edge weight in each module were selected as the hub genes for that module. Next, we further filtered the network by selecting the leading edges (the top quintile) ranked by TOM edge weight. The R package networkD3 was used to generate the force directed network on the filtered top hub genes.

### Regulatory network analysis

We used PANDA (54), a method that integrates information from gene expression, protein-protein interaction, and transcription factor binding motif data, to generate aggregate gene regulatory networks for IMR90-ER (n = 29) and IMR90-GFP (n = 27) samples. Then, LIONESS (55) was applied to extract single-sample networks. To avoid issues caused by negative edge weights, we transformed the network edges using the following equation:

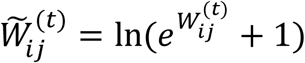

Where 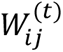 is the edge weight calculated by PANDA and LIONESS between TF(*i*) and gene(*j*) in a single-sample network (*t*).

Next, samples for each cell line were divided into 5 time periods (t1 = [0h, 4h], t2 = [8h, 12h], t3 = [16h, 20h], t4 = [24h, 32h], t5 = [40h, 48h]) and an averaged network was calculated for all individual LIONESS networks from each period. ALPACA, a method for detecting significant changes in the community structure of weighted bipartite transcriptional networks, was then applied to compare the community structure of the averaged ER and GFP networks at different time periods (56). We kept and reported differential communities that included more than 30 nodes. For each detected differential community, we performed GO enrichment analysis and reported the GO terms with *P*_adj_ < 0.05.

### Statistics

R (version 4.2.2) and GraphPad Prism (version 9.0.2) were used in all statistical tests for computational and in vitro analyses. Specific statistical methods and details are described in main and supplemental figure legends. Stata/SE (version 15.1) was utilized for the mouse xenograft study, with a comprehensive description of the model outlined in the xenograft efficacy study methods section. A p-value less than 0.05 was considered significant.

## Conflicts-of-interests statement

The authors have declared that no conflict of interest exists.

## Study approval

This study was approved by the IACUC for the care and use of laboratory animals at the University of Arizona (protocol #2021-0772).

## Data availability

The data generated in this study are publicly available in Gene Expression Omnibus (GEO) under accession numbers GSE130639, GSE229701 and GSE278335. Previously published data analyzed in this study were obtained from Gene Expression Omnibus (GEO) under accession numbers GSE39612 and GSE70138. The protein-protein interaction data used for regulatory network analysis was obtained from the STRING database (file: "9606.protein.links.v11.5.txt.gz"). The TF binding motif data for regulatory network analysis was obtained from the website https://sites.google.com/a/channing.harvard.edu/kimberlyglass/tools/resources using the link for the Human Motif Scan (Homo sapiens; hg38). All code and processed data are available on GitHub (https://github.com/JiawenYang16/pyrvinium_in_MCC). Values for all data points in graphs are reported in the Supporting Data Values file.

## Author contributions

JY conceptualized and designed the study, developed methods, conducted experiments, analyzed data, drafted, and revised the manuscript. JTL designed the study, developed methods, and analyzed data. PV developed methods, conducted experiments, and analyzed data. MGC and GDP conducted experiments. QP and HK assisted with methods development. CC assisted on statistics and data analysis. RGS provided resources and facilities for the research and assisted in study design. DJR assisted on statistics and revised the manuscript. PCG provided guidance on xenograft model development and revised the manuscript. JAD provided resources and facilities for the research and revised the manuscript for intellectual content. MP conceptualized and supervised the study and revised the manuscript for intellectual content.

## Acknowledgments

This work was supported by the NIH grant R01 CA251729 to MP. We thank the Experimental Mouse Shared Resource (EMSR) at the University of Arizona Cancer Center (UACC), supported by the National Cancer Institute of the National Institutes of Health under award number P30 CA023074, for assistance with the xenograft study. We thank John Fitch from the flow cytometry shared resource at UACC for technical guidance. We thank Yanghuan Yu from Ingmar Riedel-Kruse’s lab at the University of Arizona for assistance with imaging. We thank Julia Schnabel from James DeCaprio’s lab at Harvard Medical school for assistance with experimental materials. We thank Dr. Guang Yao at the University of Arizona, Department of Molecular and Cellular Biology, and Dr. Curtis Thorne and members of the Thorne lab at the University of Arizona, Department of Cellular and Molecular Medicine for useful discussion and suggestions.

## Notes

### Competing Interest Statement

The authors have declared no competing interest.

### Summary of Updates

In this revised version, we added a third mouse xenograft study to show pyrvinium remains effective at lower doses, added siRNA and overexpression perturbations, coupled with use of the TopGFP TCF/LEF reporter system, to show precisely how Wnt signaling regulates neuroendocrine cell fate in MCC, and Validated our results in neonatal human dermal fibroblasts (nHDF), a type of skin cell in which Merkel cell polyomavirus has been shown to replicate; We have Figure 3 - 7 revised; Supplemental files updated; authors updated.

## References

1. Becker JC, et al. Merkel cell carcinoma. Nature Reviews Disease Primers. 2017;3(1):17077.

2. DeCaprio JA. Molecular Pathogenesis of Merkel Cell Carcinoma. Annu Rev Pathology Mech Dis. 2020;16(1):1–23.

3. Lewis CW, et al. Patterns of distant metastases in 215 Merkel cell carcinoma patients: Implications for prognosis and surveillance. Cancer Med-us. 2020;9(4):1374–1382.

4. Harms KL, et al. Analysis of Prognostic Factors from 9387 Merkel Cell Carcinoma Cases Forms the Basis for the New 8th Edition AJCC Staging System. Ann Surg Oncol. 2016;23(11):3564–3571.

5. Trinidad CM, et al. Update on eighth edition American Joint Committee on Cancer classification for Merkel cell carcinoma and histopathological parameters that determine prognosis. J Clin Pathol. 2019;72(5):337.

6. Iyer JG, et al. Response rates and durability of chemotherapy among 62 patients with metastatic Merkel cell carcinoma. Cancer Med-us. 2016;5(9):2294–2301.

7. Harms PW, et al. The biology and treatment of Merkel cell carcinoma: current understanding and research priorities. Nat Rev Clin Oncol. 2018;15(12):763–776.

8. Nghiem P, et al. Durable Tumor Regression and Overall Survival in Patients With Advanced Merkel Cell Carcinoma Receiving Pembrolizumab as First-Line Therapy. J Clin Oncol. 2019;37(9):JCO.18.01896.

9. Park DE, et al. Dual inhibition of MDM2 and MDM4 in virus-positive Merkel cell carcinoma enhances the p53 response. Proc National Acad Sci. 2019;116(3):1027–1032.

10. Ananthapadmanabhan V, et al. Milademetan is a highly potent MDM2 inhibitor in Merkel cell carcinoma. Jci Insight. 2022;7(13).

11. Harms PW, et al. The Distinctive Mutational Spectra of Polyomavirus-Negative Merkel Cell Carcinoma. Cancer Res. 2015;75(18):3720–3727.

12. Knepper TC, et al. The Genomic Landscape of Merkel Cell Carcinoma and Clinicogenomic Biomarkers of Response to Immune Checkpoint Inhibitor Therapy. Clin Cancer Res. 2019;25(19):5961–5971.

13. Wong SQ, et al. UV-Associated Mutations Underlie the Etiology of MCV-Negative Merkel Cell Carcinomas. Cancer Res. 2015;75(24):5228–5234.

14. Goh G, et al. Mutational landscape of MCPyV-positive and MCPyV-negative Merkel cell carcinomas with implications for immunotherapy. Oncotarget. 2015;7(3):3403–3415.

15. González-Vela M del C, et al. Shared Oncogenic Pathways Implicated in Both Virus-Positive and UV-Induced Merkel Cell Carcinomas. J Invest Dermatol. 2017;137(1):197–206.

16. Carter MD, et al. Genetic profiles of different subsets of Merkel cell carcinoma show links between combined and pure MCPyV-negative tumors. Hum Pathol. 2018;71:117–125.

17. Liu W, et al. Identifying the Target Cells and Mechanisms of Merkel Cell Polyomavirus Infection. Cell Host Microbe. 2016;19(6):775–787.

18. Cheng J, et al. Merkel Cell Polyomavirus Large T Antigen Has Growth-Promoting and Inhibitory Activities. J Virol. 2013;87(11):6118–6126.

19. Leiendecker L, et al. LSD1 inhibition induces differentiation and cell death in Merkel cell carcinoma. EMBO Mol Med. 2020;12(11):e12525.

20. Park DE, et al. Merkel cell polyomavirus activates LSD1-mediated blockade of non-canonical BAF to regulate transformation and tumorigenesis. Nat cell Biol. 2020;22(5):603–615.

21. Arora R, et al. Survivin Is a Therapeutic Target in Merkel Cell Carcinoma. Sci Transl Med. 2012;4(133):133ra56.

22. Kim J, McNiff JM. Nuclear expression of survivin portends a poor prognosis in Merkel cell carcinoma. Modern Pathol. 2008;21(6):764–769.

23. Harms PW, et al. Next generation sequencing of Cytokeratin 20-negative Merkel cell carcinoma reveals ultraviolet-signature mutations and recurrent TP53 and RB1 inactivation. Modern Pathol. 2016;29(3):240–248.

24. Harms KL, et al. Increased expression of EZH2 in Merkel cell carcinoma is associated with disease progression and poorer prognosis. Hum Pathol. 2017;67:78–84.

25. Gartin AK, et al. Merkel Cell Carcinoma Sensitivity to EZH2 Inhibition Is Mediated by SIX1 Derepression. J Invest Dermatol. 2022;142(10):2783–2792.e15.

26. Nusse R, Varmus HE. Wnt genes. Cell. 1992;69(7):1073–1087.

27. Wodarz A, Nusse R. MECHANISMS OF WNT SIGNALING IN DEVELOPMENT. Cell Dev Biol. 1998;14(1):59–88.

28. Morin PJ, et al. Activation of β-Catenin-Tcf Signaling in Colon Cancer by Mutations in β-Catenin or APC. Science. 1997;275(5307):1787–1790.

29. Rubinfeld B, et al. Stabilization of β-Catenin by Genetic Defects in Melanoma Cell Lines. Science. 1997;275(5307):1790–1792.

30. Andersson ER, et al. Wnt5a cooperates with canonical Wnts to generate midbrain dopaminergic neurons in vivo and in stem cells. Proc Natl Acad Sci United States Am. 2013;110(7):E602–10.

31. Arredondo SB, et al. Wnt5a promotes differentiation and development of adult-born neurons in the hippocampus by noncanonical Wnt signaling. STEM CELLS. 2020;38(3):422–436.

32. Weeraratna AT, et al. Wnt5a signaling directly affects cell motility and invasion of metastatic melanoma. Cancer Cell. 2002;1(3):279–288.

33. Starrett GJ, et al. Merkel Cell Polyomavirus Exhibits Dominant Control of the Tumor Genome and Transcriptome in Virus-Associated Merkel Cell Carcinoma. mBio. 2017;8(1):e02079–16.

34. Harms PW, et al. Distinct Gene Expression Profiles of Viral-and Nonviral-Associated Merkel Cell Carcinoma Revealed by Transcriptome Analysis. J Invest Dermatol. 2013;133(4):936–945.

35. Weeraratna AT, et al. Lack of Wnt5A Expression in Merkel Cell Carcinoma. Arch Dermatol. 2010;146(1):88–89.

36. Houben R, et al. Inhibition of T-antigen expression promoting glycogen synthase kinase 3 impairs merkel cell carcinoma cell growth. Cancer Lett. 2022;524:259–267.

37. Liu S, et al. The Wnt-signaling pathway is not implicated in tumorigenesis of Merkel cell carcinoma. J Cutan Pathol. 2007;34(1):22–26.

38. Schultz CW, et al. The FDA approved anthelmintic Pyrvinium Pamoate inhibits pancreatic cancer cells in nutrient depleted conditions by targeting the mitochondria. Mol Cancer Ther. 2021;20(11):molcanther.MCT-20-0652-A.2020.

39. Esumi H, et al. Antitumor activity of pyrvinium pamoate, 6-(dimethylamino)-2-[2-(2,5-dimethyl-1-phenyl-1H-pyrrol-3-yl)ethenyl]-1-methyl-quinolinium pamoate salt, showing preferential cytotoxicity during glucose starvation. Cancer Sci. 2004;95(8):685–690.

40. Faux MC, et al. APC regulation of ESRP1 and p120-catenin isoforms in colorectal cancer cells. Mol Biol Cell. 2021;32(2):120–130.

41. Mologni L, et al. Synergistic Effects of Combined Wnt/KRAS Inhibition in Colorectal Cancer Cells. Plos One. 2012;7(12):e51449.

42. Wiegering A, et al. The impact of pyrvinium pamoate on colon cancer cell viability. Int J Colorectal Dis. 2014;29(10):1189–1198.

43. Song P, et al. β-catenin represses miR455-3p to stimulate m6A modification of HSF1 mRNA and promote its translation in colorectal cancer. Mol Cancer. 2020;19(1):129.

44. Thorne CA, et al. Small-molecule inhibition of Wnt signaling through activation of casein kinase 1α. Nat Chem Biol. 2010;6(11):829–836.

45. Xu W, et al. The Antihelmintic Drug Pyrvinium Pamoate Targets Aggressive Breast Cancer. Plos One. 2013;8(8):e71508.

46. Xiang W, et al. Pyrvinium selectively targets blast phase-chronic myeloid leukemia through inhibition of mitochondrial respiration. Oncotarget. 2015;6(32):33769–33780.

47. Lamb R, et al. Antibiotics that target mitochondria effectively eradicate cancer stem cells, across multiple tumor types: Treating cancer like an infectious disease. Oncotarget. 2015;6(7):4569–4584.

48. Venugopal C, et al. Pyrvinium Targets CD133 in Human Glioblastoma Brain Tumor–Initiating Cells. Clin Cancer Res. 2015;21(23):5324–5337.

49. Li H, et al. Pyrvinium pamoate regulates MGMT expression through suppressing the Wnt/β-catenin signaling pathway to enhance the glioblastoma sensitivity to temozolomide. Cell Death Discov. 2021;7(1):288.

50. Fu Y-H, et al. Deciphering the Role of Pyrvinium Pamoate in the Generation of Integrated Stress Response and Modulation of Mitochondrial Function in Myeloid Leukemia Cells through Transcriptome Analysis. Biomed. 2021;9(12):1869.

51. Wander P, et al. High-throughput drug screening reveals Pyrvinium pamoate as effective candidate against pediatric MLL-rearranged acute myeloid leukemia. Transl Oncol. 2021;14(5):101048.

52. Wang M, et al. Role of the Unfolded Protein Response Regulator GRP78/BiP in Development, Cancer, and Neurological Disorders. Antioxid Redox Sign. 2009;11(9):2307–2316.

53. Fattakhova E, et al. Identification of the FDA-Approved Drug Pyrvinium as a Small-Molecule Inhibitor of the PD-1/PD-L1 Interaction. Chemmedchem. 2021;16(18):2769–2774.

54. Glass K, et al. Passing Messages between Biological Networks to Refine Predicted Interactions. PLoS ONE. 2013;8(5):e64832.

55. Kuijjer ML, et al. Estimating Sample-Specific Regulatory Networks. iScience. 2019;14:226–240.

56. Padi M, Quackenbush J. Detecting phenotype-driven transitions in regulatory network structure. npj Syst Biol Appl. 2018;4(1):16.

57. Yu D-H, et al. Pyrvinium Targets the Unfolded Protein Response to Hypoglycemia and Its Anti-Tumor Activity Is Enhanced by Combination Therapy. Plos One. 2008;3(12):e3951.

58. Park DE, et al. Dual inhibition of MDM2 and MDM4 in virus-positive Merkel cell carcinoma enhances the p53 response. Proc National Acad Sci. 2019;116(3):1027–1032.

59. Berger A, et al. N-Myc-mediated epigenetic reprogramming drives lineage plasticity in advanced prostate cancer. J Clin Invest. 2019;129(9):3924–3940.

60. Berrios C, et al. Merkel Cell Polyomavirus Small T Antigen Promotes Pro-Glycolytic Metabolic Perturbations Required for Transformation. PLoS Pathog. 2016;12(11):e1006020.

61. Gupta P, et al. Merkel Cell Polyomavirus Downregulates N-myc Downstream-Regulated Gene 1, Leading to Cellular Proliferation and Migration. J Virol. 2020;94(3).

62. Cho WC, et al. SOX11 Is an Effective Discriminatory Marker, When Used in Conjunction With CK20 and TTF1, for Merkel Cell Carcinoma: Comparative Analysis of SOX11, CK20, PAX5, and TTF1 Expression in Merkel Cell Carcinoma and Pulmonary Small Cell Carcinoma. Arch Pathol Lab Med. 2023;147(7):758–766.

63. Grumolato L, et al. β-Catenin-Independent Activation of TCF1/LEF1 in Human Hematopoietic Tumor Cells through Interaction with ATF2 Transcription Factors. PLoS Genet. 2013;9(8):e1003603.

64. Roemer K. Mutant p53: Gain-of-Function Oncoproteins and Wild-Type p53 Inactivators. Biol Chem. 1999;380(7–8):879–887.

65. Veija T, et al. Hotspot mutations in polyomavirus positive and negative Merkel cell carcinomas. Cancer Genet-ny. 2016;209(1–2):30–35.

66. Houben R, et al. Mechanisms of p53 Restriction in Merkel Cell Carcinoma Cells Are Independent of the Merkel Cell Polyoma Virus T Antigens. J Invest Dermatol. 2013;133(10):2453–2460.

67. Schultz CW, Nevler A. Pyrvinium Pamoate: Past, Present, and Future as an Anti-Cancer Drug. Biomed. 2022;10(12):3249.

68. Chen D, et al. Dermal β-catenin activity in response to epidermal Wnt ligands is required for fibroblast proliferation and hair follicle initiation. Development. 2012;139(8):1522–1533.

69. Perdigoto CN, et al. Polycomb-Mediated Repression and Sonic Hedgehog Signaling Interact to Regulate Merkel Cell Specification during Skin Development. PLoS Genet. 2016;12(7):e1006151.

70. Rosso SB, Inestrosa NC. WNT signaling in neuronal maturation and synaptogenesis. Front Cell Neurosci. 2013;7:103.

71. He C-W, Liao C-P, Pan C-L. Wnt signalling in the development of axon, dendrites and synapses. R Soc Open Biol. 2018;8(10):180116.

72. Frost TC, et al. YAP1 and WWTR1 expression inversely correlates with neuroendocrine markers in Merkel cell carcinoma. J Clin Investig. 2023;133(5):e157171.

73. Park HW, et al. Alternative Wnt Signaling Activates YAP/TAZ. Cell. 2015;162(4):780–794.

74. Starrett GJ, et al. Clinical and molecular characterization of virus-positive and virus-negative Merkel cell carcinoma. Genome Med. 2020;12(1):30.

75. MacLaine NJ, et al. A Central Role for CK1 in Catalyzing Phosphorylation of the p53 Transactivation Domain at Serine 20 after HHV-6B Viral Infection*. J Biol Chem. 2008;283(42):28563–28573.

76. Huart A-S, et al. CK1α Plays a Central Role in Mediating MDM2 Control of p53 and E2F-1 Protein Stability. J Biol Chem. 2009;284(47):32384–32394.

77. Gartin AK, et al. Merkel Cell Carcinoma Sensitivity to EZH2 Inhibition Is Mediated by SIX1 Derepression. J Invest Dermatol. 2022;142(10):2783–2792.e15.

78. Ponzini FM, et al. Repurposing the FDA-approved anthelmintic pyrvinium pamoate for pancreatic cancer treatment: study protocol for a phase I clinical trial in early-stage pancreatic ductal adenocarcinoma. BMJ Open. 2023;13(10):e073839.

79. Cheng J, et al. Merkel cell polyomavirus recruits MYCL to the EP400 complex to promote oncogenesis. Plos Pathog. 2017;13(10):e1006668.

80. Langfelder P, Horvath S. WGCNA: an R package for weighted correlation network analysis. BMC Bioinform. 2008;9(1):559.

