## Supplemental Figure for "Integrative analysis reveals therapeutic potential of pyrvinium pamoate in Merkel cell carcinoma"


**Supplemental Figures**

**Supplemental Figures 1-12.** **p1-15**

**Supplemental Tables**

**Supplemental Table 1-8.** **p16-23**

**Supplemental Methods** **p24-26**

**Supplemental Materials**

**Plasmid constructs.** **p27-38**

**siRNA sequences.** **p39**

**References** **p40**

### Supplemental Figures

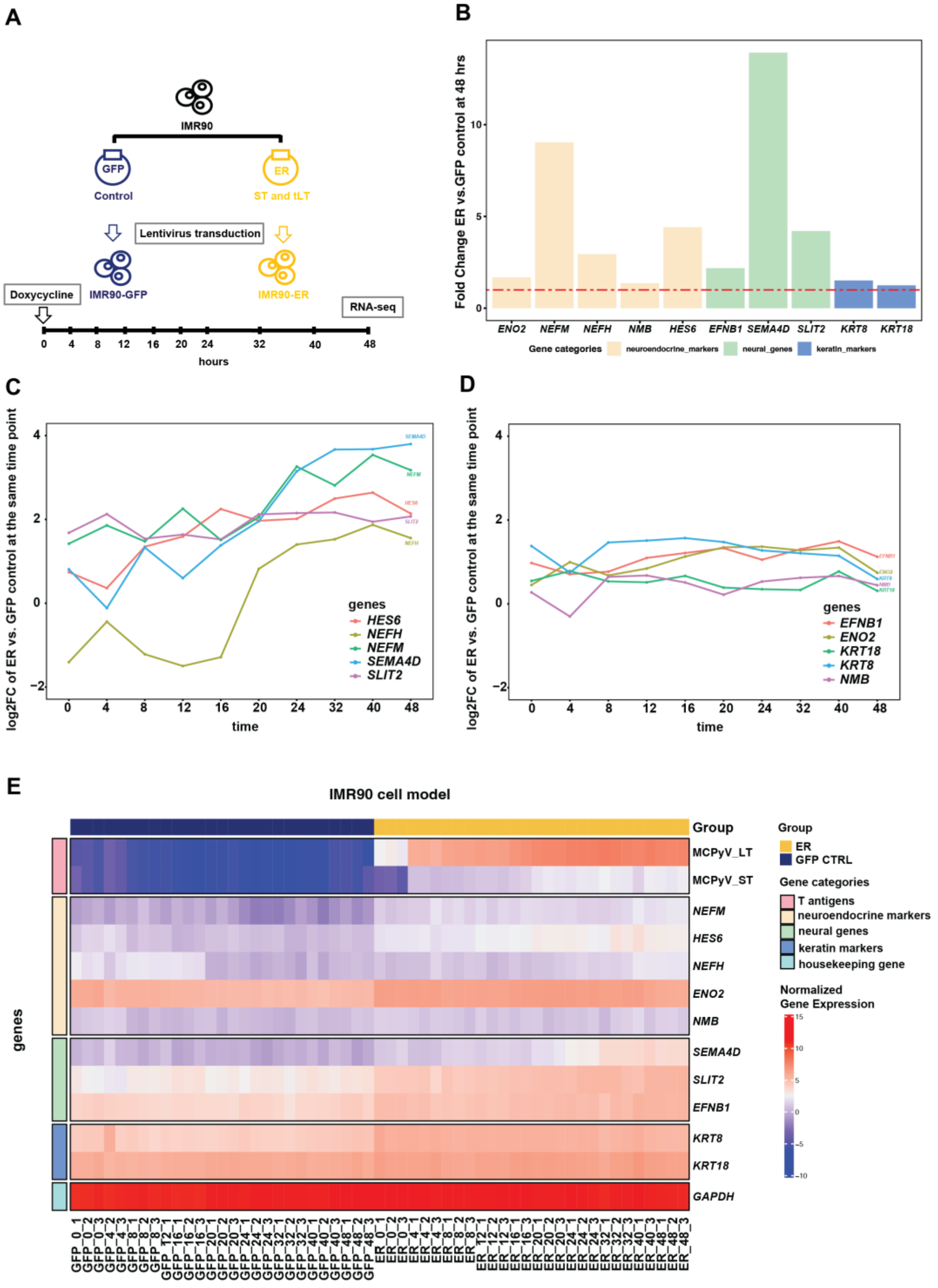


**Supplemental Figure 1. MCPyV-ER upregulates a subset of Merkel cell carcinoma markers.** (**A**) Schematic diagram of the IMR90-ER and IMR90-GFP induction model. (**B**) Fold change in the expression of some MCC marker genes in IMR90-ER compared to IMR90-GFP at 48 hours. (**C-D**) Temporal fold change in a subset of MCC marker genes in IMR90-ER versus IMR90-GFP inducible samples at each time point. (**E**) Heatmap showing normalized gene expression levels for T antigen genes, subset of MCC marker genes, and the housekeeping gene *GAPDH*.

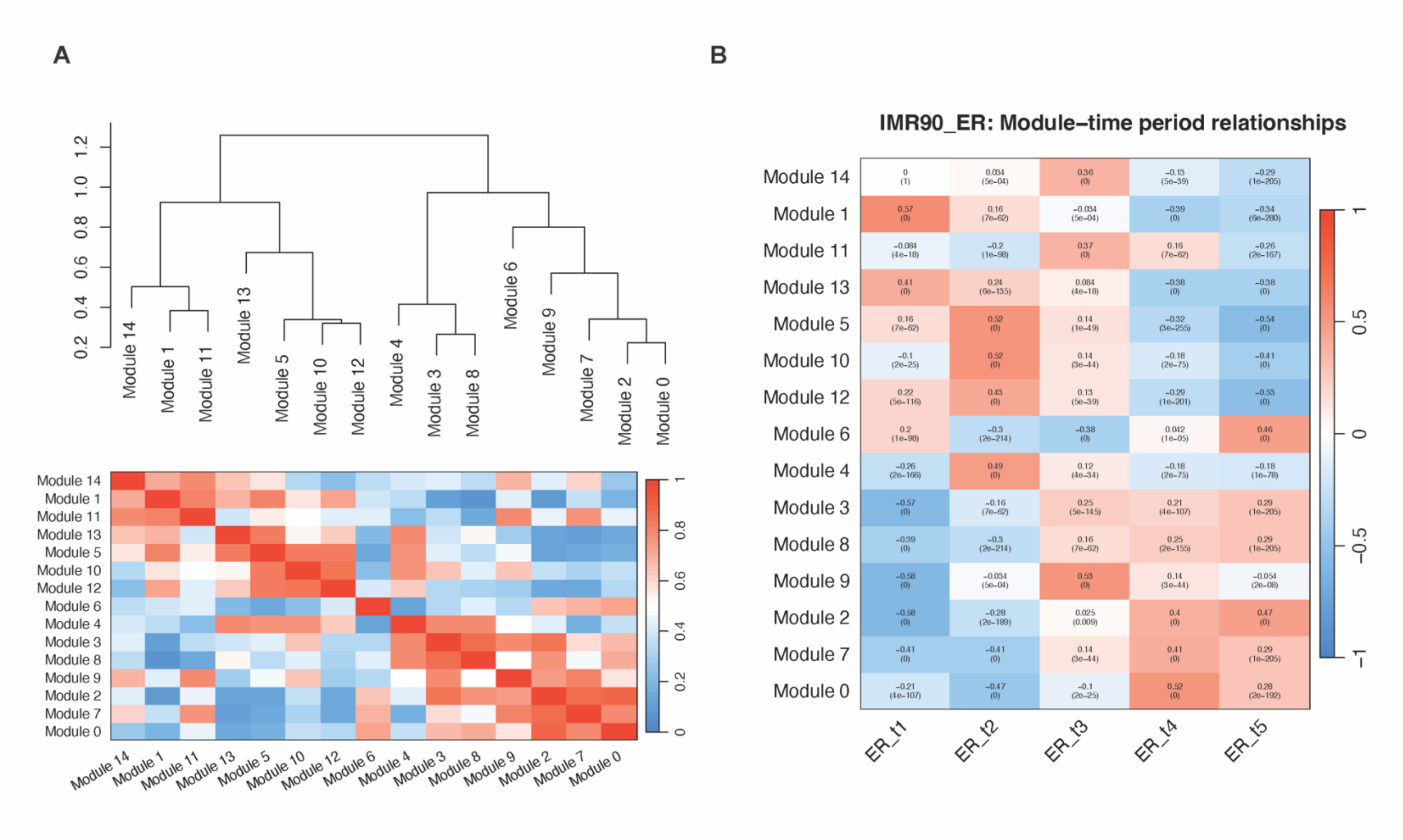


**Supplemental Figure 2. WGCNA eigengenes analysis on IMR90-ER data.** (**A**) Heatmap and dendrogram illustrating eigengenes correlations and the hierarchical clustering of eigengenes. (**B**) Correlations between module eigengenes and time period, calculated using the Kendall rank correlation coefficient.

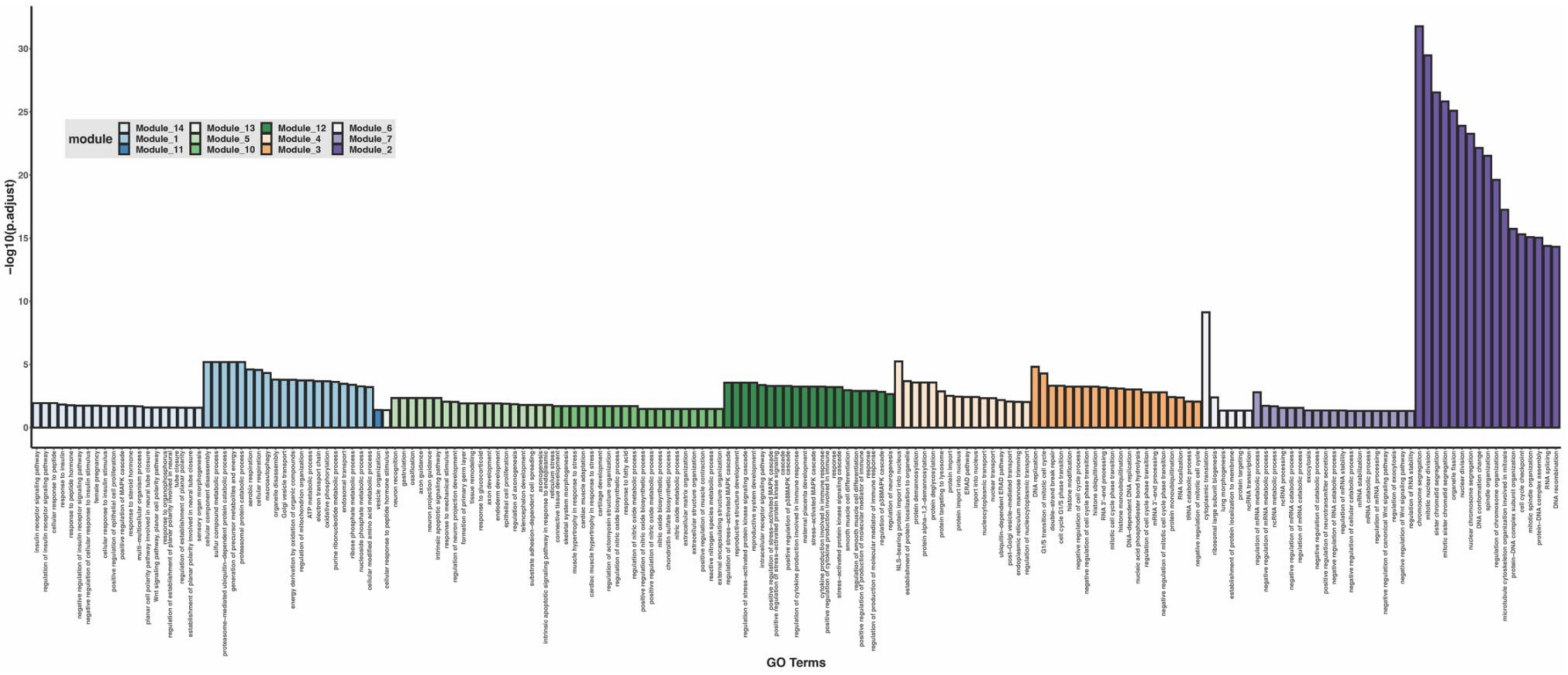


**Supplemental Figure 3. Gene co-expression modules in MCPyV-ER and their enriched biological functions.** Over-representation GO term enrichment results for 14 gene co-expression modules in MCPyV-ER samples. Only significant ($P_{adj}$ < 0.05) terms from each module are presented.

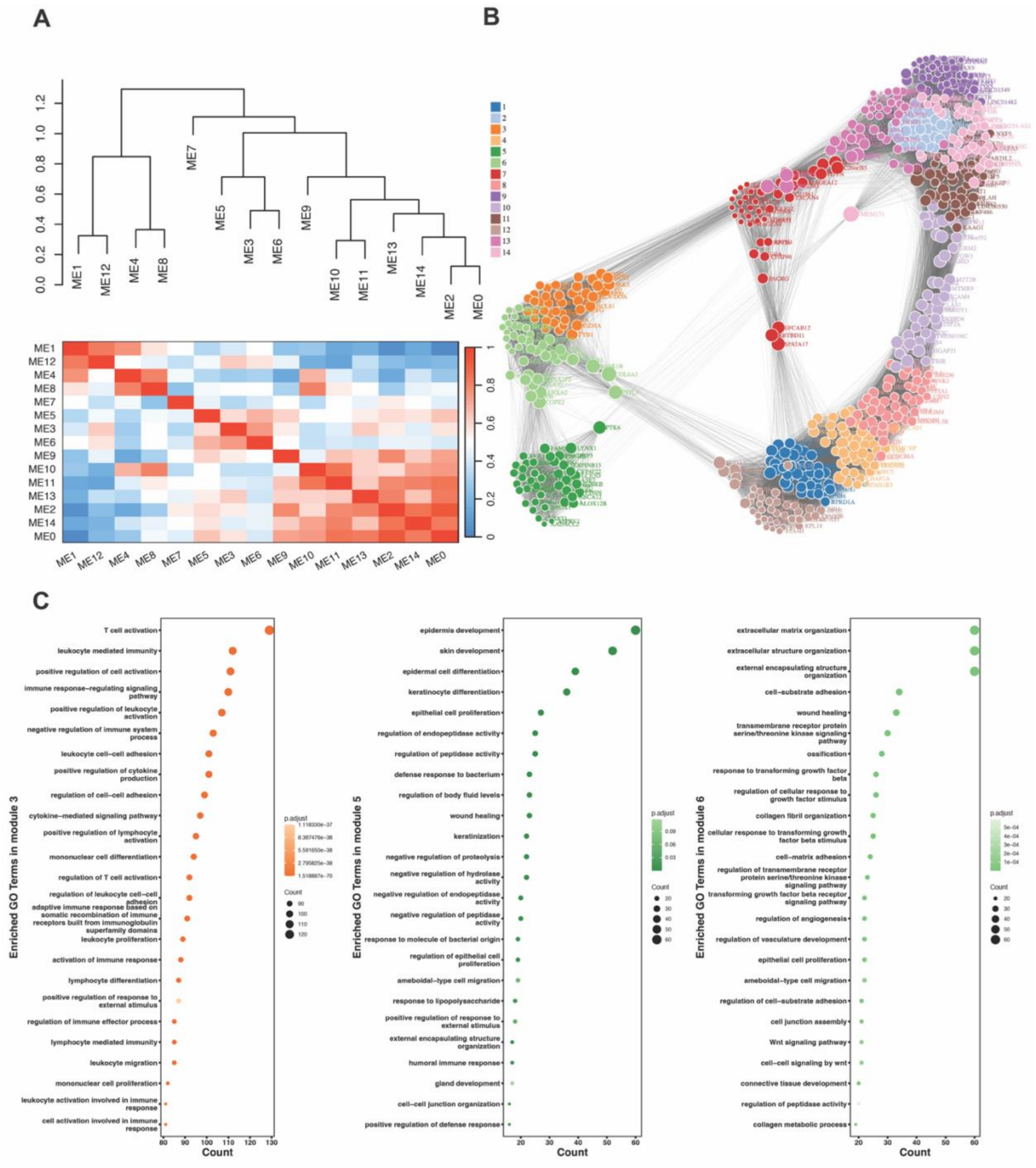


**Supplemental Figure 4. WGCNA analysis on MCC patient samples reveals that Wnt signaling pathway is highly co-expressed with MCC-related pathways.** (**A**) Heatmap and dendrogram illustrating eigengenes correlations and the hierarchical clustering of eigengenes, based on differentially expressed genes between MCC tumor samples and normal skin samples in a cohort of 30 MCC tumor samples. (**B**) Force-directed network of hub genes in 14 modules derived from MCC tumor samples. The attraction forces between genes were defined by their topological overlaps and were inversely proportional to the length of the edges in the graph. (**C**) GO term enrichment results for genes in module 3, 5 and 6 respectively.

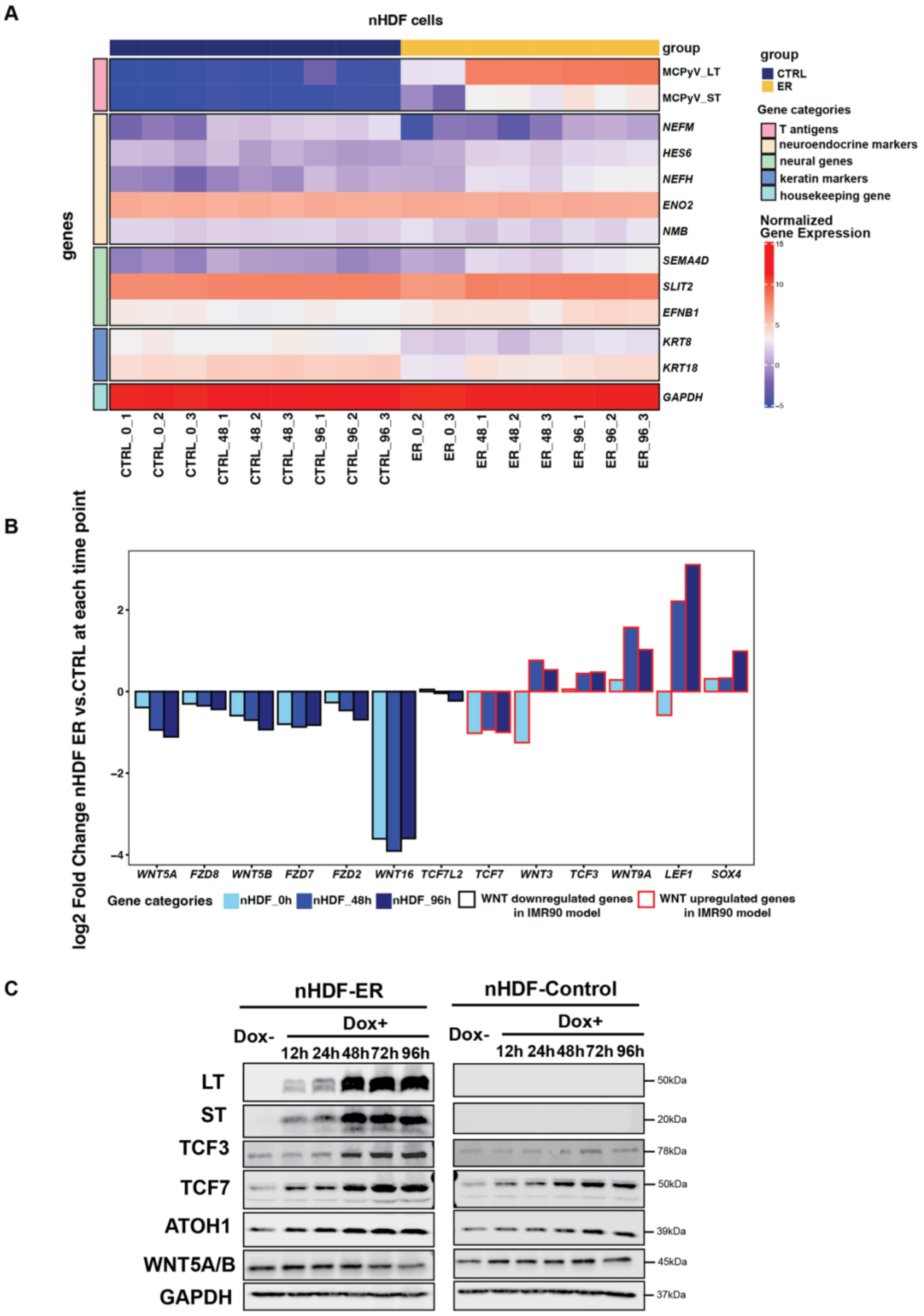


**Supplemental Figure 5. The MCPyV-ER inducible model in nHDF cells supports findings from the IMR90 model.** (**A**) Heatmap showing normalized gene expression levels for T antigen genes, a subset of MCC marker genes, and housekeeping gene *GAPDH* in nHDF-ER and nHDF-Control samples. (**B**) Bar plot illustrating the log2 fold changes of WNT gene expression levels in nHDF-ER samples across all time points, relative to the nHDF-Control samples at the corresponding time points. (**C**) Protein levels of MCPyV large T antigen, small T antigen, Wnt genes and NE marker ATOH1 in nHDF-ER and nHDF-Control cells at 0, 12, 24, 48, 72 and 96 hours following ER expression induction.

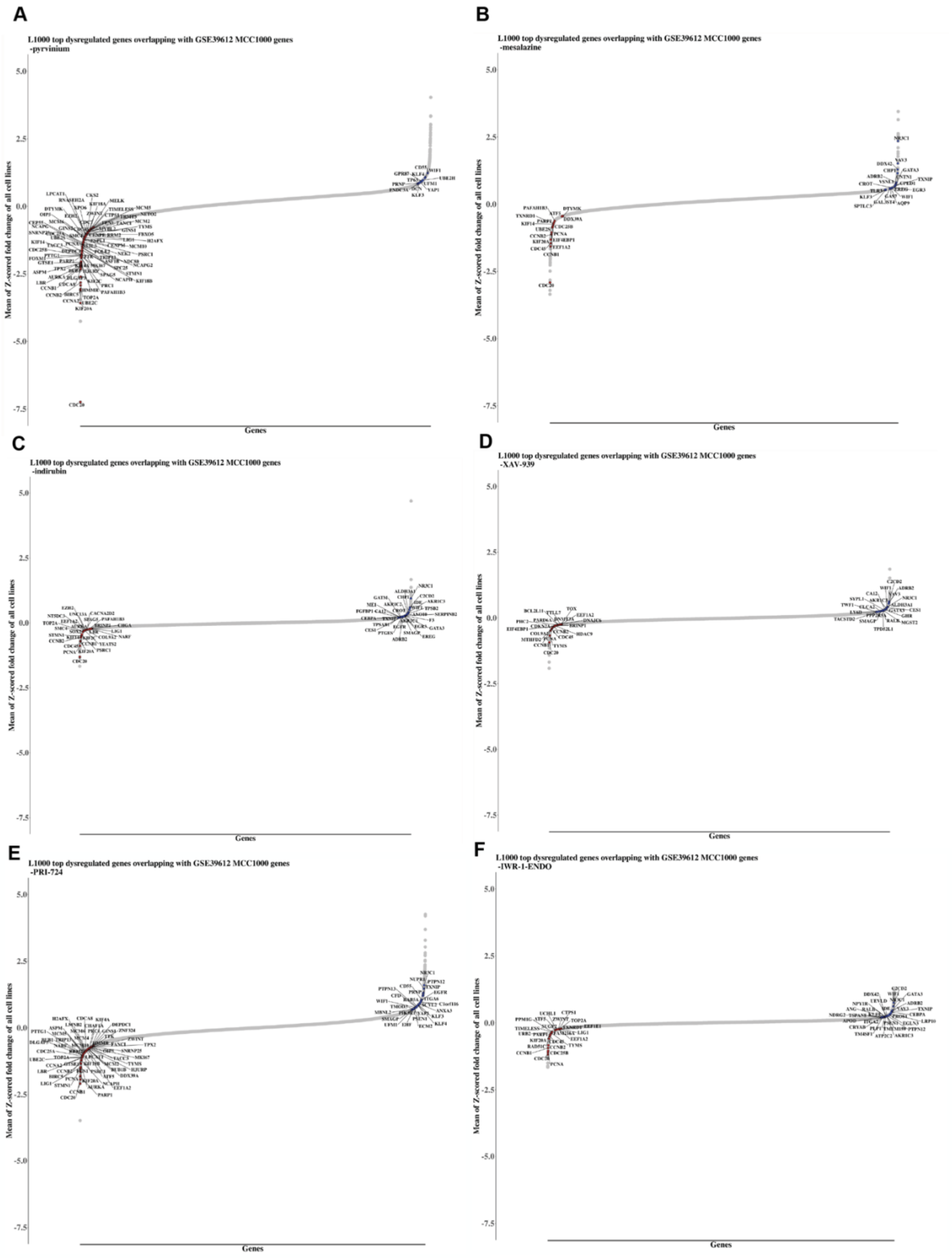


**Supplemental Figure 6. Transcriptomic analysis of Wnt signaling perturbagens in LINCS L1000 and MCC tumor databases.** (**A - F**) Six commercially available WNT signaling perturbagens reverse MCC1000 genes in the L1000 dataset. All cell lines from LINCS L1000 dataset were analyzed, and the top 500 reversed MCC1000 and L1000 genes from both sides were annotated.

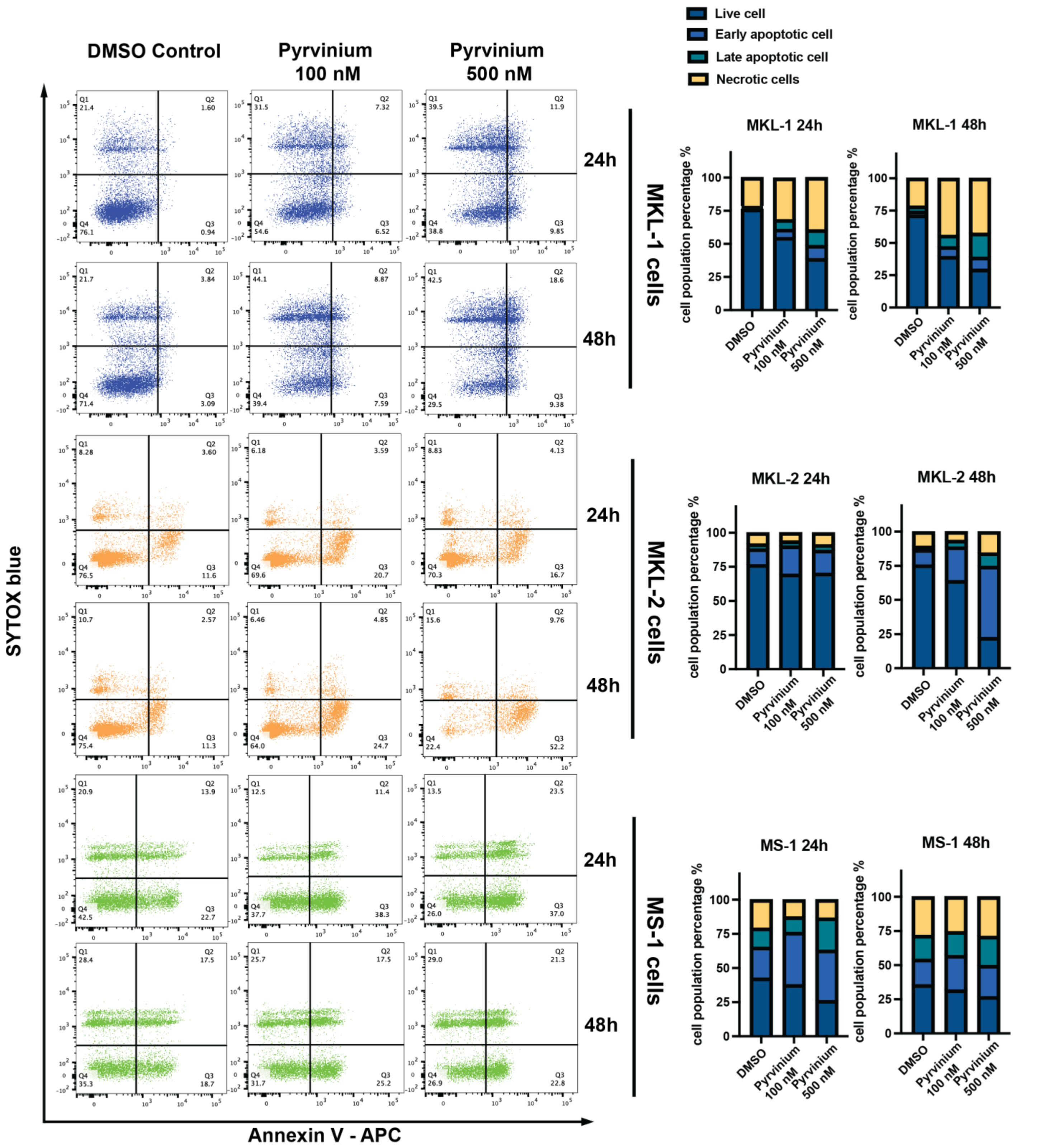


**Supplemental Figure 7. Pyrvinium pamoate induces apoptosis in MCC cells.** Flow cytometry analyses showing the levels of Annexin V-APC and SYTOX blue staining to assess the apoptotic population in MKL-1, MKL-2, and MS-1 cells treated with varying doses of pyrvinium pamoate over different time periods. Bar graphs on the right display the quantification of different populations in pyrvinium-treated MCC cells at 24h and 48h, respectively.

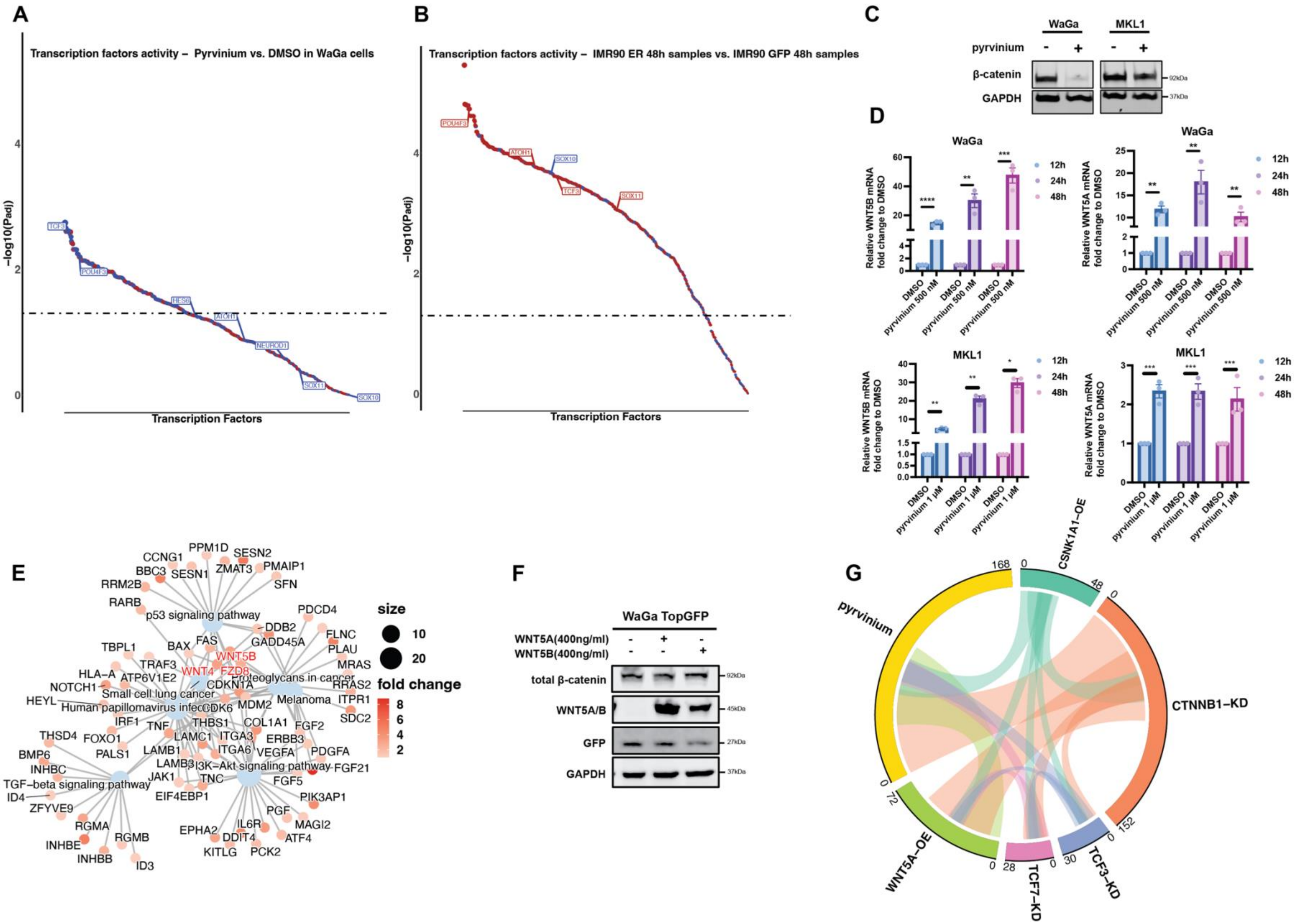
A
Transcription factors activity – Pyrvinium vs. DMSO in WaGa cells
-log10(Padj)
Transcription Factors
B
Transcription factors activity – IMR90 ER 48h samples vs. IMR90 GFP 48h samples
-log10(Padj)
Transcription Factors
C
WaGa
MKL1
pyrvinium
β-catenin
GAPDH
D
WaGa
MKL1
Relative WNT5B mRNA fold change to DMSO
Relative WNT5A mRNA fold change to DMSO
12h
24h
48h
DMSO
pyrvinium 500 nM
pyrvinium 1 μM
E
p53 signaling pathway
Small cell lung cancer
Human papillomavirus infection
Proteoglycans in cancer
Melanoma
PI3K-Akt signaling pathway
TGF-beta signaling pathway
size
fold change
F
WaGa TopGFP
WNT5A(400ng/ml)
WNT5B(400ng/ml)
total β-catenin
WNT5A/B
GFP
GAPDH
G
pyrvinium
CSNK1A1-OE
CTNNB1-KD
TCF3-KD
TCF7-KD
WNT5A-OE

**Supplemental Figure 8. Pyrvinium reverses Wnt signaling in MCC by targeting multiple proteins.** (**A-B**) Scatter plots showing the predicted activity levels of master transcription factors in pyrvinium-treated vs. DMSO-treated MCC cells and in IMR90-ER vs. IMR90-GFP cells at 48 hours, respectively. The MCCP specific regulon used in VIPER was constructed using ARACNe on 13 virus-positive MCC patient samples. (**C**) Total β-catenin protein levels in WaGa and MKL-1 cells under DMSO and 1μM pyrvinium treatment, as measured by WB. (**D**) RT-qPCR validation of relative *WNT5B* and *WNT5A* mRNA levels in WaGa and MKL-1 cells treated with pyrvinium at 500 nM and 1 μM respectively for 12, 24, and 48 hours, using *GAPDH* as the internal control. The mean relative mRNA levels in vehicle control samples were set to 1. Statistical significance was determined by an unpaired two sample t-test (n = 3). (****, $P < 0.0001$; ***, $P < 0.001$; **, $P < 0.01$; *, $P < 0.05$). (**E**) Gene-biological concepts network showing KEGG pathway enrichment analysis results of significantly upregulated DEGs ($P_{adj} \leqslant 0.05$, log2 fold change $\geqslant 1$) in WaGa cells. (**F**) Total β-catenin and GFP protein levels in WaGa TopGFP cells treated with 400 ng/ml human WNT5A and WNT5B recombinant proteins, as measured by WB. (**G**) Circos plot depicting the pairwise comparison of overlapping MCC1000 signature genes and the top dysregulated genes across different perturbations of the Wnt signaling pathway by pyrvinium.

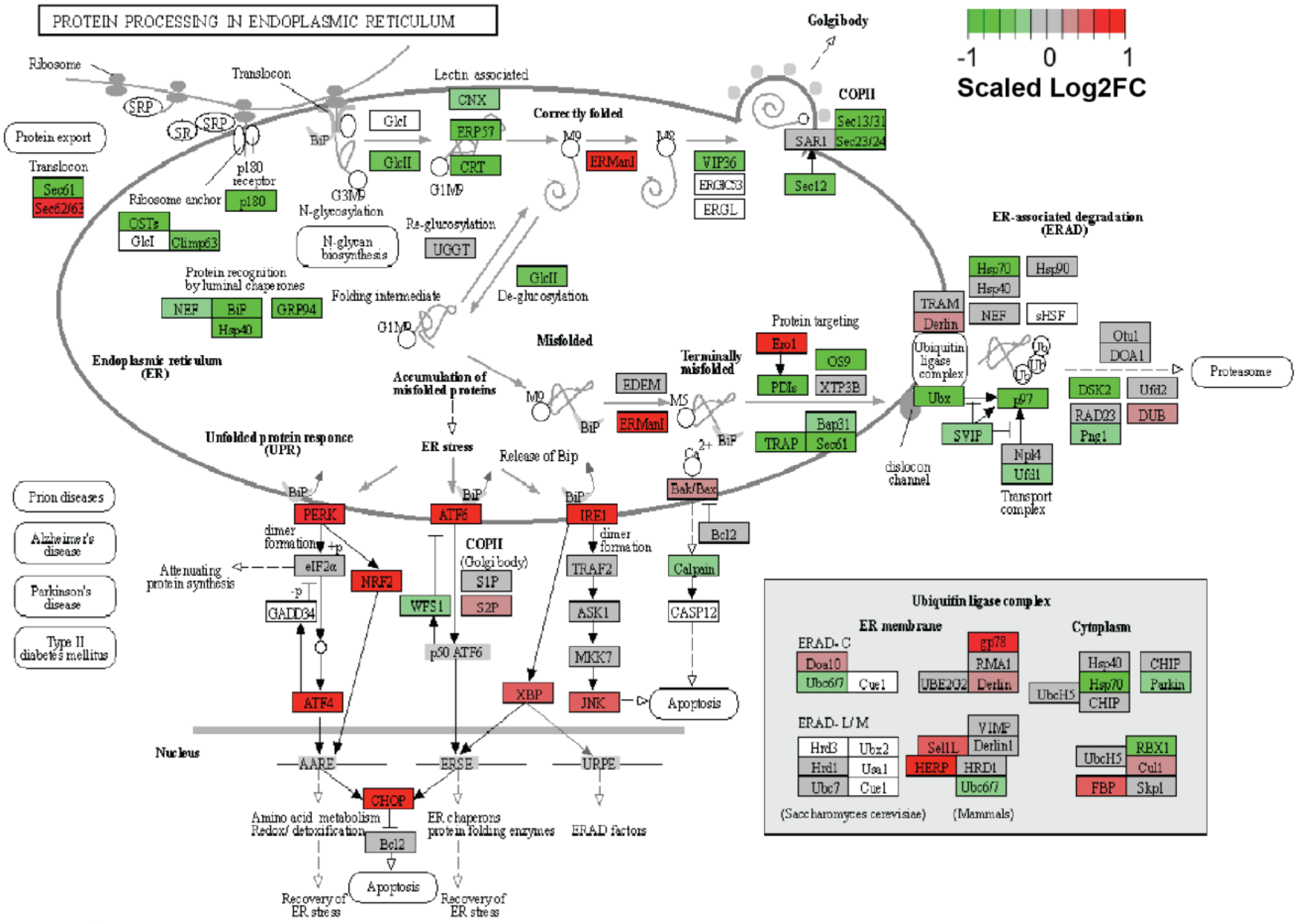


**Supplemental Figure 9. Pyrvinium pamoate activates endoplasmic reticulum (ER) stress.** KEGG pathway visualization of UPR and ER stress related genes. Gene coloring reflects the log2 fold change from the comparison between pyrvinium- and DMSO-treated WaGa cells after 24 hours. The log2 fold change values are scaled between -1 and 1.

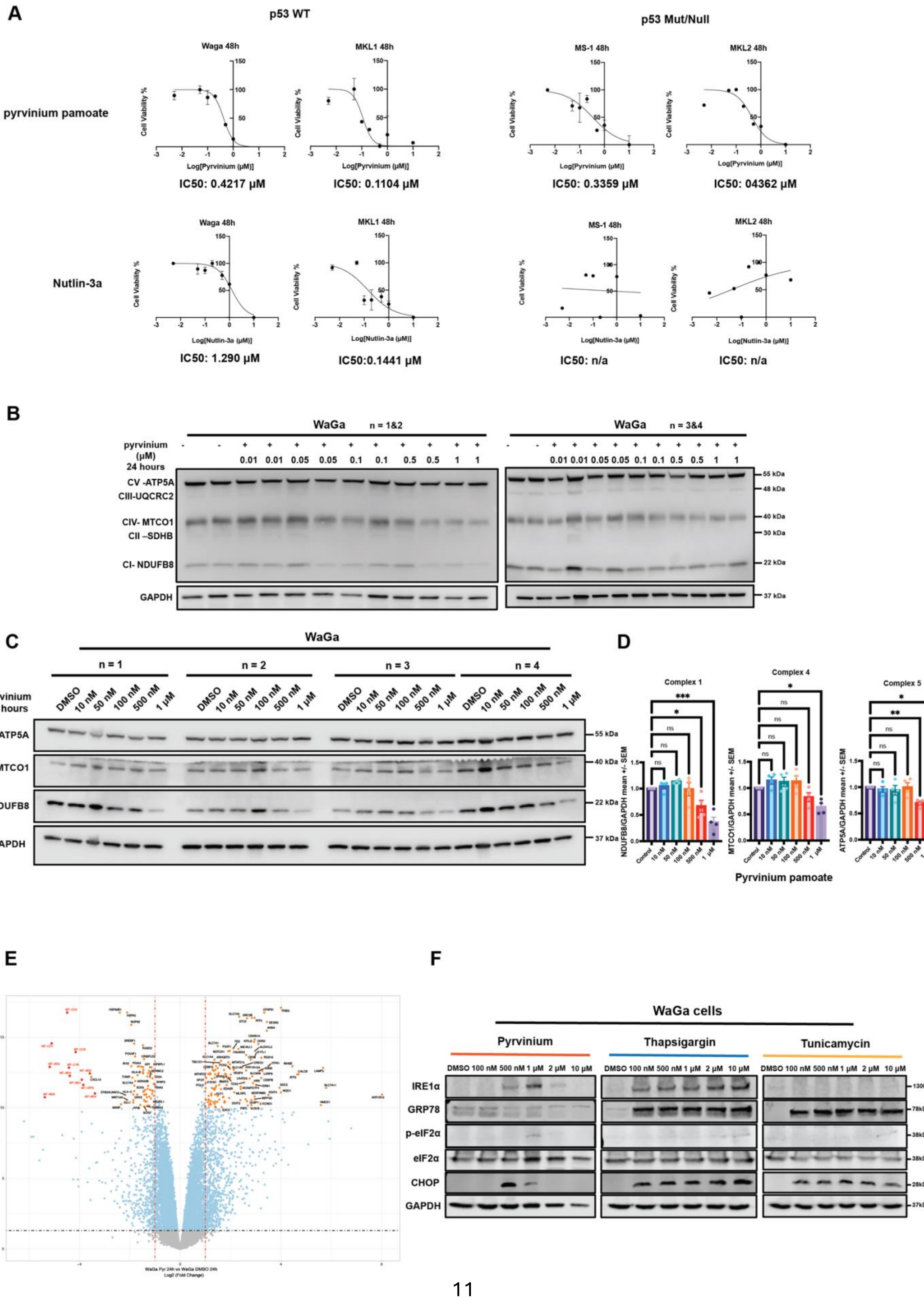
A
p53 WT
p53 Mut/Null
pyrvinium pamoate
Nutlin-3a
Waga 48h
MKL1 48h
MS-1 48h
MKL2 48h
Cell Viability %
Log[Pyrvinium (µM)]
Log[Nutlin-3a (µM)]
IC50: 0.4217 µM
IC50: 0.1104 µM
IC50: 0.3359 µM
IC50: 04362 µM
IC50: 1.290 µM
IC50:0.1441 µM
IC50: n/a
IC50: n/a
B
WaGa n = 1&2
WaGa n = 3&4
pyrvinium (µM) 24 hours
0.01 0.01 0.05 0.05 0.1 0.1 0.5 0.5 1 1
CV -ATP5A
CIII-UQCRC2
CIV- MTCO1
CII –SDHB
CI- NDUFB8
GAPDH
55 kDa
48 kDa
40 kDa
30 kDa
22 kDa
37 kDa
C
WaGa
n = 1
n = 2
n = 3
n = 4
pyrvinium 24 hours
DMSO 10 nM 50 nM 100 nM 500 nM 1 µM
CV -ATP5A
CIV- MTCO1
CI- NDUFB8
GAPDH
55 kDa
40 kDa
22 kDa
37 kDa
D
Complex 1
Complex 4
Complex 5
NDUFB8/GAPDH mean +/- SEM
MTCO1/GAPDH mean +/- SEM
ATP5A/GAPDH mean +/- SEM
Control 10 nM 50 nM 100 nM 500 nM 1 µM
Pyrvinium pamoate
E
WaGa Pyr 24h vs WaGa DMSO 24h
Log2 (Fold Change)
-log10(Padj)
F
WaGa cells
Pyrvinium
Thapsigargin
Tunicamycin
DMSO 100 nM 500 nM 1 µM 2 µM 10 µM
IRE1α
GRP78
p-eIF2α
eIF2α
CHOP
GAPDH
130kDa
78kDa
38kDa
28kDa
37kDa

**Supplemental Figure 10. Mechanisms of action of pyrvinium pamoate in MCC**. (**A**) Half-maximal inhibitory concentration (IC50) values for pyrvinium and Nutlin-3a after 48 hours of treatment in p53 wild-type cell lines (WaGa, MKL-1) and $TP53^{Mut}/TP53^{-/-}$ cell lines (MS-1, MKL-2), measured using the MTT assay. (**B-C**) Two independent sets of OXPHOS Western blot analyses showing levels of mitochondria complex proteins in WaGa cells treated with pyrvinium at various concentration for 24 hours (n = 4). (**D**) Quantification of protein levels for NDUFB8 (Complex I subunit), MTCO1 (Complex IV subunit), ATP5A (Complex V subunit), and internal control GAPDH in panel (**C**). Data are presented relative to mean GAPDH expression in vehicle control samples (means ± SEM, n = 4 biological replicate blots). Statistical analysis was performed using ordinary ANOVA followed by Dunnett's multiple comparison test. (***, $P < 0.001$; **, $P < 0.01$; *, $P < 0.05$). Quantitative data for western blot result in (**B**) are included in Figure 6G. (**E**) Volcano plot illustrating results from DEG analysis. The x-axis represents log2 fold change, and the y-axis indicates -log10($P_{adj}$). (**F**) Protein levels of CHOP and ER-stress markers were detected by WB in WaGa cells treated with pyrvinium, thapsigargin and tunicamycin at various concentration for 24 hours.

**A**

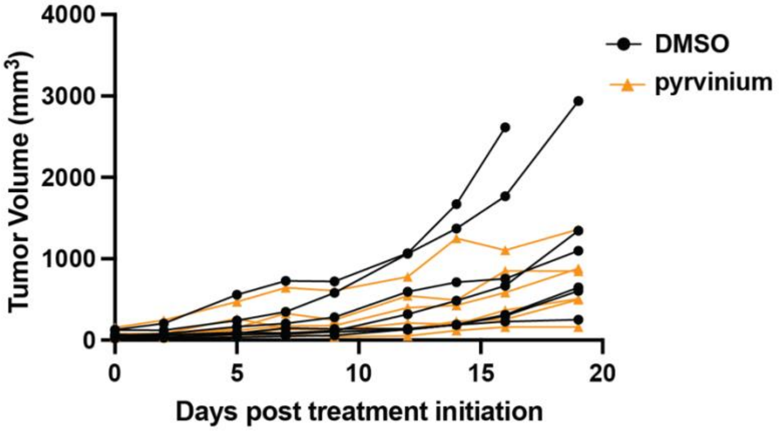


**B**

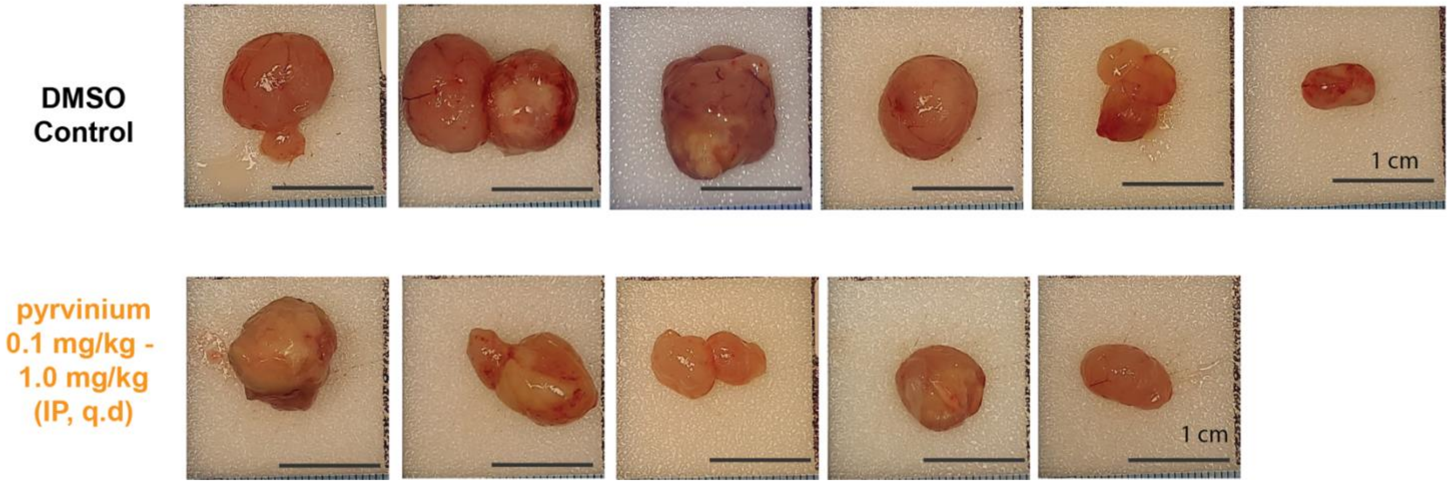


**Supplemental Figure 11. Pyrvinium pamoate inhibits tumor growth in an MCC xenograft model.** (**A**) MKL-1 xenograft tumor growth curve from in vivo Study #3 depicting individual tumor volumes in the vehicle control and pyrvinium-treated groups from day 0 to day 20. Treatment followed a gradually increasing dosage schedule of 0.1-1.0 mg/kg per day (q.d.). (**B**) Images of xenograft tumor collected from in vivo Study #3 with the 0.1 -1.0 mg/kg q.d. dosing schedule. Tumors were harvested 0.5 – 1 hour after the final treatment on day 20. (One mouse in the vehicle control group was sacrificed on day 16 because its tumor volume exceeded 2000 mm$^3$. In the treatment group, one mouse was euthanized on day 7 due to severe dermatitis on its neck, which was determined to be unrelated to pyrvinium toxicity. Another mouse in the treatment group was sacrificed on day 20, prior to the final injection, due to treatment-related side effects.) All images are scaled uniformly.

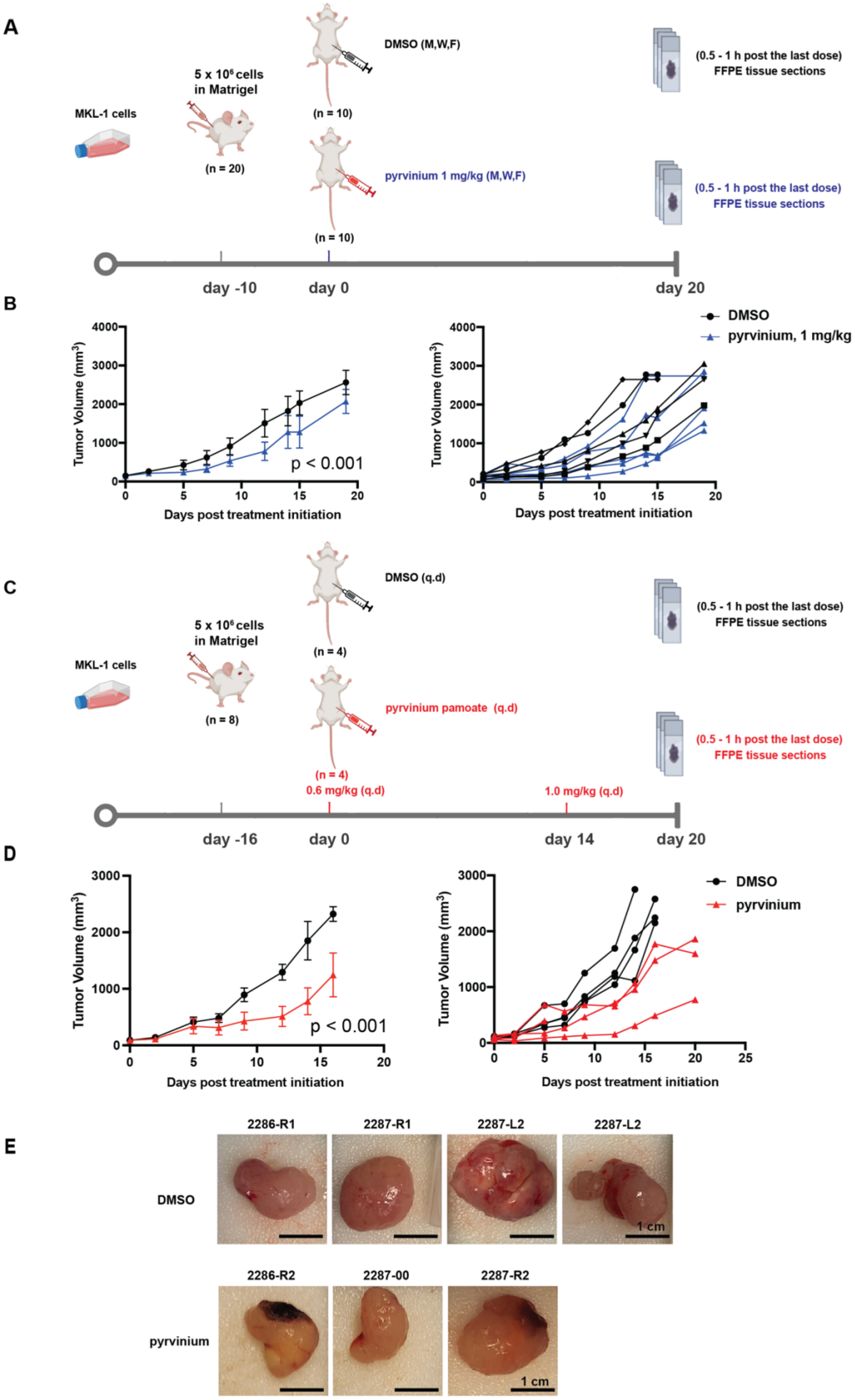

A
MKL-1 cells
5 x 10^6 cells in Matrigel
(n = 20)
DMSO (M,W,F)
(n = 10)
pyrvinium 1 mg/kg (M,W,F)
(n = 10)
(0.5 - 1 h post the last dose) FFPE tissue sections
(0.5 - 1 h post the last dose) FFPE tissue sections
day -10
day 0
day 20
B
Tumor Volume (mm³)
Days post treatment initiation
p < 0.001
DMSO
pyrvinium, 1 mg/kg
C
MKL-1 cells
5 x 10^6 cells in Matrigel
(n = 8)
DMSO (q.d)
(n = 4)
pyrvinium pamoate (q.d)
(n = 4)
0.6 mg/kg (q.d)
1.0 mg/kg (q.d)
(0.5 - 1 h post the last dose) FFPE tissue sections
(0.5 - 1 h post the last dose) FFPE tissue sections
day -16
day 0
day 14
day 20
D
Tumor Volume (mm³)
Days post treatment initiation
p < 0.001
DMSO
pyrvinium
E
2286-R1
2287-R1
2287-L2
2287-L2
DMSO
1 cm
2286-R2
2287-00
2287-R2
pyrvinium
1 cm

**Supplemental Figure 12. Pyrvinium pamoate inhibits tumor growth in MCC xenograft models – pilot studies.**

(**A**) Schematic representation of the experimental design for in vivo Study #1, featuring a treatment schedule of 1.0 mg/kg administered three days per week. (**B**) Tumor growth curve illustrating both the mean and individual tumor sizes in vehicle control and pyrvinium-treated mice from day 0 to day 20 in in vivo Study #1. (**C**) Schematic representation of the experimental design for in vivo Study #2, which employed a daily treatment schedule with doses ranging from 0.6 -1.0 mg/kg per day. (**D**) Tumor growth curve showing the mean and individual tumor volumes in vehicle control and pyrvinium-treated groups from day 0 to day 19 in in vivo Study #2. (**E**) Images of xenograft tumor tissues collected from in vivo Study #2. Tumors were harvested following the final 1.0 mg/kg dose of pyrvinium or DMSO control. (One mouse in the treatment group was sacrificed prior to the dose escalation to 1.0 mg/kg due to treatment-related side effects.) All images are scaled uniformly.

### Supplemental Tables

**Supplemental Table 1. MCPyV T antigens transcripts read count matrix from IMR90 ER samples**

| | MCPyV_gp1 (capsid protein) | MCPyV_gp2 (VP1, major capsid protein) | MCPyV_gp3 (large T-antigen) | MCPyV_gp4 (small T-antigen) |
|---|---|---|---|---|
| **ER_0_1** | 0 | 0 | 261 | 3 |
| **ER_0_2** | 0 | 0 | 268 | 3 |
| **ER_0_3** | 0 | 0 | 271 | 2 |
| **ER_4_1** | 0 | 0 | 2297 | 68 |
| **ER_4_2** | 0 | 0 | 1769 | 50 |
| **ER_4_3** | 0 | 0 | 1285 | 35 |
| **ER_8_1** | 0 | 0 | 2640 | 51 |
| **ER_8_2** | 0 | 0 | 2851 | 52 |
| **ER_8_3** | 0 | 0 | 2507 | 55 |
| **ER_12_1** | 0 | 0 | 4720 | 96 |
| **ER_12_2** | 0 | 0 | 3659 | 82 |
| **ER_12_3** | 0 | 0 | 4336 | 86 |
| **ER_16_1** | 0 | 0 | 5370 | 96 |
| **ER_16_3** | 0 | 0 | 5299 | 90 |
| **ER_20_1** | 0 | 0 | 7276 | 167 |
| **ER_20_2** | 0 | 0 | 6043 | 151 |
| **ER_20_3** | 0 | 0 | 7510 | 180 |
| **ER_24_1** | 0 | 0 | 9560 | 169 |
| **ER_24_2** | 0 | 0 | 7018 | 146 |
| **ER_24_3** | 0 | 0 | 9645 | 193 |
| **ER_32_1** | 0 | 0 | 7492 | 175 |
| **ER_32_2** | 0 | 0 | 12703 | 252 |
| **ER_32_3** | 0 | 0 | 12368 | 273 |
| **ER_40_1** | 0 | 0 | 6080 | 80 |
| **ER_40_3** | 0 | 0 | 11485 | 255 |
| **ER_48_1** | 0 | 0 | 19499 | 475 |
| **ER_48_2** | 0 | 0 | 2820 | 65 |
| **ER_48_3** | 0 | 0 | 7869 | 203 |

**Supplemental Table 2. In vivo Study #1 – 1 mg/kg IP injection tumor volume statistics with mixed-effects model**

| sqrvolume | Coef. | Std. Err. | z | P>\|z\| | [95% Conf. | Interval] |
|---|---|---|---|---|---|---|
| 1.Treatment | 3.795161 | 4.610891 | 0.82 | 0.410 | -5.242019 | 12.83234 |
| day | 2.241025 | .1057545 | 21.19 | 0.000 | 2.03375 | 2.4483 |
| Treatment#c.day | | | | | | |
| 1 | -.5139983 | .1395729 | -3.68 | 0.000 | -.7875561 | -.2404404 |
| _cons | -16.13178 | 3.324688 | -4.85 | 0.000 | -22.64805 | -9.615512 |

**[sqrvolume]day + [sqrvolume]1.Treatment#c.day = 0**

| sqrvolume | Coef. | Std. Err. | z | P>\|z\| | [95% Conf. | Interval] |
|---|---|---|---|---|---|---|
| (1) | 1.727027 | .0910855 | 18.96 | 0.000 | 1.548503 | 1.905551 |

**Supplemental Table 3. In vivo Study #2 – 0.6 – 1.0 mg/kg IP injection tumor volume statistics with mixed-effects model**

| sqrvolume | Coef. | Std. Err. | z | P>\|z\| | [95% Conf. | Interval] |
|---|---|---|---|---|---|---|
| 1.Treatment | 20.4434 | 4.940943 | 4.14 | 0.000 | 10.75933 | 30.12747 |
| day | 2.613092 | .1218058 | 21.45 | 0.000 | 2.374357 | 2.851827 |
| Treatment#c.day | | | | | | |
| 1 | -1.181657 | .1646966 | -7.17 | 0.000 | -1.504456 | -.8588573 |
| _cons | -35.12977 | 3.600441 | -9.76 | 0.000 | -42.18651 | -28.07304 |

**[sqrvolume]day + [sqrvolume]1.Treatment#c.day = 0**

| sqrvolume | Coef. | Std. Err. | z | P>\|z\| | [95% Conf. | Interval] |
|---|---|---|---|---|---|---|
| (1) | 1.431436 | .1108527 | 12.91 | 0.000 | 1.214168 | 1.648703 |

**Supplemental Table 4. In vivo Study #3 – 0.1 – 1.0 mg/kg IP injection tumor volume statistics with mixed-effects model**

| sqrvolume | Coef. | Std. Err. | z | P>\|z\| | [95% Conf. | Interval] |
|---|---|---|---|---|---|---|
| 1.Treatment | 6.616166 | 4.256251 | 1.55 | 0.120 | -1.725932 | 14.95826 |
| day | 1.339024 | .0823468 | 16.26 | 0.000 | 1.177627 | 1.500421 |
| Treatment#c.day | | | | | | |
| 1 | -.4020586 | .1191159 | -3.38 | 0.001 | -.6355214 | -.1685958 |
| _cons | -11.69644 | 2.997815 | -3.90 | 0.000 | -17.57205 | -5.820831 |

**[sqrvolume]day + [sqrvolume]1.Treatment#c.day = 0**

| sqrvolume | Coef. | Std. Err. | z | P>\|z\| | [95% Conf. | Interval] |
|---|---|---|---|---|---|---|
| (1) | .9369654 | .0860674 | 10.89 | 0.000 | .7682763 | 1.105654 |

**Supplemental Table 5. Chemicals used in the study**

| Reagent | Supplier | Catalog number |
|---|---|---|
| Pyrvinium pamoate | MedChemExpress | MFCD00010090 |
| Nutlin-3a | Selleck Chemicals | S8059 |
| WNT5A | R&D systems Inc | 645WN010 |
| WNT5B | R&D systems Inc | 7347WN025 |
| Tunicamycin | Sigma-Aldrich | T7765 |
| Thapsigargin | Sigma-Aldrich | T9033 |

**Supplemental Table 6. Antibodies used in the study**

| Antibodies | Supplier | Catalog number | RRID number |
|---|---|---|---|
| Anti-GAPDH | Thermo Fisher Scientific | MA5-15738 | AB_10977387 |
| Anti-β-catenin | BD Bioscience | 610154 | AB_397555 |
| Anti-MCPyV T antigens (1) | The DeCaprio Lab | Custom generation | |
| Anti-eIF2α | Cell Signaling Technology | 3192 | AB_2095847 |
| Anti-phosphorylated eIF2α | | 3179 | AB_2095853 |
| Anti-PARP | | 9542 | AB_2160739 |
| Anti-PUMA | | 4976 | AB_2064551 |
| Anti-WNT5A/B | | 2530 | AB_2215595 |
| Anti-TCF3 | | 2883 | AB_2199136 |
| Anti-TCF7 | | 2203 | AB_2199302 |
| Anti-IRE1α | | 3294 | AB_823545 |
| Anti-GRP78 | | 3177 | AB_2119845 |
| Anti-p53 | Santa Cruz Biotechnology | sc-126 | AB_628082 |
| Anti-GFP | | sc-9996 | AB_627695 |
| Anti-OXPHOS Cocktail | Abcam | ab110411 | AB_2756818 |
| Anti-Ki67 | | ab16667 | AB_302459 |
| Anti-ATOH1 | Proteintech | 21215-1-AP | AB_10733126 |

**Supplemental Table 7. Optimized settings for FCCP-OCR test for WaGa cells**

| Settings | Cycles | Command | Time (min) |
|---|---|---|---|
| Calibrate | | | |
| Basal | 3 | Loop Start | 3 |
| | | Mix | 2 |
| | | Measure | 2.50 |
| Loop End | | | |
| Inject – Port A | | | |
| FCCP - 0.5 µM | 3 | Loop Start | 3 |
| | | Mix | 2 |
| | | Measure | 2.50 |
| Loop End | | | |

**Supplemental Table 8. Primers used for RT-qPCR assay**

| Primer | Supplier | Catalog number |
|---|---|---|
| AXIN2 | Bio-Rad | qHsaCIP0031547 |
| SOX2 | | qHsaCED0036871 |
| WNT5A | | qHsaCIP0028356 |
| WNT5B | | qHsaCID0038673 |
| ATOH1 | | qHsaCED0019647 |
| GAPDH | | qHsaCED0038674 |
| TCF3 (NM_003200) | IDT (Customized) | Forward: CGAGAAGCCCCAGACCAAAC Reverse: ACCTTTTCCTCTTCTCGCCG |
| TCF7 (NM_201632) | IDT (Customized) | Forward: ACCAGCGGCATGTACAAAGA Reverse: GTGGGATGTGGGCTGTTGAA |

### Supplemental Methods

#### Cell Viability and Cell Proliferation

MCC cells were seeded in 96 well plates at a density of $5 \times 10^4$ cells in 100 µl. Water soluble MTT (Thiazolyl blue tetrazolium bromide) (Sigma-Aldrich, cat: #M5655) compound was dissolved in DPBS to 5 mg/ml. Upon each measurement time point,10 µl of 5 mg/ml MTT was added to each 100 µl of cell suspension in each well of a 96 well plate and the plate was incubated at 37℃ for 3 hours. After incubation, 100 µl of acidified isopropanol was added into each well. Then, the plate was wrapped in foil and shaken on an orbital shaker for 10 mins at 37℃. The BioTek Synergy LX plate reader was used to read the 96 well plate at 562 nm.

#### Flow Cytometry

For apoptosis assay, MCC cells were seeded in 25 $cm^2$ cell culture flasks and were treated with pyrvinium for 24 hours before collection. Cells were centrifuged at 400 rcf for 5 min and washed in cold PBS once. Then, cells were washed with binding buffer. Next, cells were resuspended in 100 µl binding buffer at 5 x $10^6$ cells/mL. 5 µl of Annexin V-APC (Thermo Fisher Scientific, cat: #A35110) was added to 100 µl of the cell suspension. After 15 minutes incubation at room temperature in the dark, the cells were spun down and resuspended in 500 µl binding buffer. 0.5 µl of SYTOX blue (Thermo Fisher Scientific, cat: #S34857) was added to each reaction and the cell suspension was transferred into 1ml Falcon tubes. The cells were incubated on ice for 15 min before measuring by a flow cytometer (BD FACS Canto II) in the allophycocyanin (APC) and Pacific Blue channels.

#### Immunofluorescence

WaGa cells were seeded in 25 $cm^2$ cell culture flasks and were treated with pyrvinium for 24 hours. Cells under different conditions were collected and washed with DPBS once. Then, DPBS was used to resuspend the cells in $2 \times 10^5$ cells/ml. After funnels were assembled for the Cytospin, 200 ul of cell suspension of each condition was added to the funnel, and the cells were centrifuged at 800 rpm for 5 min with high acceleration. Slides were air-dried for 5 min. Next, the cells were fixed on the slides with 4% paraformaldehyde in DPBS for 15 min at room temperature. After rinsing the cells two times in DPBS, the DPBS was aspirated and the cells were permeabilized with 0.1% Triton X-100 in DPBS for 10 min. The cells were again washed with DPBS twice, and samples were blocked in 3% BSA (in DPBS) for 30 min at room temperature. Then, incubation of primary and secondary antibodies was performed under manufacturer recommended conditions. The Antifade (Thermo Fisher Scientific, cat: #P36935) mounting media with DAPI and 1.5 mm slides were added to cover the cells. The samples were imaged under a fluorescent microscope (Nikon Ti2 inverted Microscope) 24 hours later.

**Immunohistochemistry**

Immunohistochemistry was performed on paraffin embedded xenograft tumor tissue sections (5 μm). The sections were first deparaffinized in xylene for 5 mins twice. Then, samples were hydrated using 100%, 95% and 75% ethanol for 3 mins each. After ethanol hydration, the samples were rinsed with distilled water for 5 minutes twice. Antigen retrieval was done with citrate buffer (pH 6.2), under near-boiling temperature for 20 min. Antibodies were diluted 1:200 in goat serum and applied to sections overnight at 4℃ after 30 mins of blocking. UltraVision LP Detection System (Fisher Scientific, cat: # TL015HD) was used for signal detection. Staining procedures were performed followed the recommended conditions by the manufacturer. IHC profiler (2), an automated method that uses color deconvolution and computerized pixel profiling to assign scores to images, was applied to quantify the IHC images based on the DAB color spectrum.

**RNA-seq processing pipeline**

The raw reads were aligned with the reference genome “GDC.h38.d1.vd1 STAR2 Index Files (v36)” downloaded from https://gdc.cancer.gov/about-data/gdc-data-processing/gdc-reference-files using STAR 2.7.10a and counts were quantified using the Rsubread R package. The raw data and counts matrix can be found in the Gene Expression Omnibus (GEO) database (accession numbers: GSE130639, GSE229701, and GSE278335). The edgeR pipeline using TMM normalization and voom with default parameters was used to normalize the counts matrix. To filter out low-expressing genes, we performed calculations of counts per million (cpm) using the raw read counts matrix. Genes with less than half of the samples exceeding 1 cpm were subsequently removed.

**Differential expression analysis**

Differentially expressed genes were identified for RNA-seq data and for the publicly available microarray dataset GSE39612 dataset using limma R package.

**Gene Ontology (GO) term enrichment analysis**

GO term over-representation analysis was performed using R packages ClusterProfiler and msigdbr. Only the Biological Pathway GO terms were included in the analysis. Hypergeometric test followed by Benjamini-Hochberg (BH) adjustment was used to calculate the adjusted p-value ($P_{adj}$).

**TF activities analysis**

VIPER (3), a method for inferring the activity of transcription factors (TFs) from gene expression data, was utilized to predict TF activities in our samples. The human DoRothEA (4) TF-target interaction database was used to generate the regulon, and only interactions with high confidence score (“A”, “B” and “C”) were included. ARACNe-AP (5), an algorithm for gene regulatory network reconstruction, was used to create tissue-specific regulons. The normalized gene expression matrices containing GFP control and ER samples were then transformed to regulatory protein activity matrices by using the viper function in the R viper package. For each TF within the regulon dataset, a Student’s t-test was performed and BH adjustment was used to calculate an adjusted p-value for altered TF activity.

**L1000 data analysis**

The LINCS L1000 Level 4 data was obtained from https://clue.io/releases/data-dashboard. The file “level4_beta_all_n3026460x12328.gctx” was downloaded to extract the z-scored fold change of both drug-treated and genetic modified samples relative to the plate vehicle control. Samples treated with pyrvinium pamoate, XAV-939, indirubin, IWR-1-ENDO, mesalazine, and PRI-724 with dosage ≤ 10 μM were selected to perform the Wnt signaling small molecule compound perturbagen analysis. Samples with CSNK1A1 and WNT5A overexpression, as well as TCF3/7 and CTNNB1 knockdown, were selected for Wnt signaling genetic modification perturbation analysis. The files “GSE92742_Broad_LINCS_cell_info.txt” and “GSE92742_Broad_LINCS_gene_info.txt” were downloaded to annotate the cell lines and gene symbols. The CMapR package was used to parse the .gctx file. MCC signature gene reversal analysis was performed using Fisher’s exact test in R with its function fisher.test().

**Supplemental Materials**

**Plasmids sequences:**

**>pLIX_402_ER_MCCL21 (9217 bp)**

aaaaaaaattagtcagccatggggcggagaatgggcggaactgggcggagttaggggcgggatgggcggagttaggggcg
ggatagctagagccagacatgataagatacattgatgagtttggacaaaccacaactagaatgcagtgaaaaaaatgctt
tatttgtgaaatttgtgatgctattgctttatttgtaaccattataagctgcaataaacaagttcctctcactctctgat
attcatttctttgcaagttataaatactgaataataagatgacatgaactactactgctagagattttccacactgacta
aaagggtctgagggatctctagttaccagagtcacacaacagacgggcacacactacttgaagcactcaaggcaagcttt
attgaggcttaagcagtgggttccctagttagccagagagctcccaggctcagatctggtctaaccagagagacccagta
caagcaaaaagcagatcttgtcttcgttgggagtgaattagcccttccagtccccccttttcttttaaaaagtggctaag
atctacagctgccttgtaagtcattggtcttaaaggtacctctagtccggacgccgaggcgaaacaggcggggaggcggc
ccaaagggagatccgactcgtctgagggcgaaggcgaagacgcggaagaggccgcagagccggcagcaggccgcgggaag
gaaggtccgctggattgagggccgaagggacgtagcagaaggacgtcccgcgcagaatccaggtggcaacacaggcgagc
agccatggaaaggacgtcagcttccccgacaacaccacggaattgtcagtgcccaacagccgagcccctgtccagcagcg
ggcaaggcaggcggcgatgagttccgccgtggcaatagggagggggaaagcgaaagtcccggaaaggagctgacaggtgg
tggcaatgccccaaccagtggggggttgcgtcagcaaacacagtgcacaccacgccacgttgcctgacaacgggccacaac
tcctcataaagagacagcaaccaggatttatacaaggaggagaaaatgaaagccatacgggaagcaatagcatgatacaa
aggcattaaagcagcgtatccacatagcgtaaaaggagcaacatagttaagaatacgatatcttggatcgatccttactt
agttacccggggagcatgtcaaggtcaaaatcgtcaagagcgtcagcaggcagcatatcaaggtcaaagtcgtcaagggc
atcggctgggagcatgtctaagtcaaaatcgtcaagggcgtcggtcggcccgccgctttcgcactttagctgtttctcca
ggccacatatgattagttccaggccgaaaaggaaggcaggttcggctccctgccggtcgaacagctcaattgcttgtctc
agaagtgggggcatagaatcggtggtaggtgtctctctttcctcttttgctacttgatgctcctgttcctccaatacgca
gcccagtgtaaagtggcccacggcggacagagcgtacagtgcgttctccagggagaagccttgctgacacaggaacgcga
gctgattttccagggtttcgtactgtttctctgttgggcgggtgccgagatgcactttagccccgtcgcgatgtgagagg
agagcacagcggtatgacttggcgttgttccgcagaaagtcttgccatgactcgccttccaggggcagaagtgggtatg
atgcctgtccagcatctcgattggcagggcatcgagcagggcccgcttgttcttcacgtgccagtacagggtaggctgct
caactcccagcttttgagcgagtttccttgtcgtcaggccttcgataccgacaccattgagtaattccagagctccgttt
atgactttgctcttgtccagtctagacattggaccagggttttcttcaacatcaccacaagtgaggagagaacctctacc
ttcggcaccgggcttgcgggtcatgcaccaggtgcgcggtccttcgggcacctcgacgtcggcggtgacggtgaagccga
gccgctcgtagaaggggaggttgcggggcgcggaggtctccaggaaggcgggcaccccggcgcgctcggccgcctccact
ccggggagcacgacggcgctgcccagacccttgccctggtggtcgggcgagacgccgacggtggccaggaaccacgcggg
ctccttgggccggtgcggcgccaggaggccttccatctgttgctgcgcggccagccgggaaccgctcaactcggccatgc
gcgggccgatctcggcgaacaccgcccccgcttcgacgctctccggcgtggtccagaccgccaccgcggcgccgtcgtcc
gcgacccacaccttgccgatgtcgagcccgacgcgcgtgaggaagagttcttgcagctcggtgacccgctcgatgtggcg
gtccggatcgacggtgtggcgcgtggcggggtagtcggcgaacgcggcggcgagggtgcgtacggccctggggacgtcgt
cgcgggtggcgaggcgcaccgtgggcttgtactcggtcatggtgaattgctggggagagaggtcggtgattcggtcaacg
agggagccgactgccgacgtgcgctccggaggcttgcagaatgcggaacaccgcgcgggcaggaacagggcccacactac
cgccccacaccccgcctcccgcaccgccccttcccggccgctgctctcggcgcgccctgctgagcagccgctattggcca

cagcccatcgcggtcggcgcgctgccattgctccctggcgctgtccgtctgcgagggtactagtgagacgtgcggcttcc
gtttgtcacgtccggcacgccgcgaaccgcaaggaaccttcccgacttaggggcggagcaggaagcgtcgccgggggcc
cacaagggtagcggcgaagatccgggtgacgctgcgaacggacgtgaagaatgtgcgagacccagggtcggcgccgctgc
gtttcccggaaccacgcccagagcagccgcgtccctgcgcaaacccagggctgccttggaaaaggcgcaaccccaacccc
ggatccttagtggtggtggtggtggtggaccggacgcgtttacgcataatccggcacatcatacggataaccggtaacca
ctttgtacaagaaagctgggtcttattgagaaaaagtaccagaatcttgggtttcttcagtttcctcagggccctcttcc
tcaataagaatattgagcagagggtcctgaccagcttctacattttctatcatttgacaaatttaccatatgatatttc
actctgtaaaatttgcttccagtttttaatttaactagcagagcttgcagagcttcgggacccccaaattttcgctttc
ttgagaatggaggaggggtcttcggggtggtgaaggaggaggattcgtattcctcatctgtaaactgagatgacgaggcc
tcctcggcagaggaagacgggggctgccggggcgagcttcttgaggagggggctcctcaggctcctcagaggacgaggg
aggctcaggggaggaaagtgattcatcgcagaagagatcctcccaggtgccatcagttctggaagaatttctaggtacac
tggttccattgggtgtgctggattctcttcctgaattggtggtctcctctctgctactggatccagaggatgaggtgggt
tcctcattgtgttcgggaggtatatcgggtcctctggactgggagtctgaagcctgggacgctgagaaggacccataccc
agaggaagagctctggctgtggggtggtgagcttccactgggggctcccctggatgcattggaggaaggctttctggatc
ttgagttggtcccgtgtggattgggcccatattcgtatgccttcccgaagctgaatcctcctgatctccaccattctttg
aatttagtggtcccatatataggggcctcgtcaacctagatgggaaagtacagaaaatctgtcataaataacctttcttt
gatattttgccttatagacttttccatatctaatacttacagaggaaggaagtaggagtctagaaaaggtgcagatgcag
taagcagtagtcagtttcttctaaagtttttttgccaccagtcaaaactttcccaagtaggaggaaatccaaaccaaagaa
taaagcactgatagcaaaaacactctccccacgtcagacagtttttttgctttaaagtttttagactacaatgctggcga
gacaacttacagctaatacaagcgcacttagaatctctaagttgcttaagcatgcacccaggacctctgcaaaatctagc
attatatccactttgcatataatcctttaaagttccatattcttcccaaggaaattttgtactgacctcatcaaacatag
agaagtcacttctgagcttgtggatattttgctggaatttgctccaaagggtgttcaattccatcattataacaggattt
cccccttatcagggtgatgctttaagcagcttcttttgaaagcagctttcatcagagggatgttgccataacaattagg
agcaatctctaaaagcttgcagagagcctctctttctttcctatttaggactaaatccatgcctgcttttttgtacaaac
ttgttgatcaatcgatgctagccaattctccaggcgatctgacggttcactaaacgagctctgcttatataggcctccca
ccgtacacgcctacctcgacatacgttctctatcactgatagggagtaaactcgacatacgttctctatcactgataggg
ataaactcgacatacgttctctatcactgatagggagtaaactcgacatacgttctctatcactgatagggagtaaactc
gacatacgttctctatcactgatagggagtaaactcgacatcgttctctatcactgatagggagtaaactcgacatacgt
tctctatcactgatagggagtaaactcgacatatcgattcgcggccaaagtggatctctgctgtccctgtaataaacccg
aaaattttgaatttttgtaatttgtttttgtaattctttagtttgtatgtctgttgctattatgtctactattctttccc
ctgcactgtacccccaatccccccttttcttttaaaattgtggatgaatactgccatttgtctcgaggtcgagaattgt
cccctcggggttgggaggtgggtctgaaacgataatggtgaatatccctgcctaactctattcactatagaaagtacagc
aaaaactattcttaaacctaccaagcctcctactatcattatgaataattttatataccacagccaatttgttatgttaa
accaattccacaaacttgcccatttatctaattccaataattcttgttcattcttttcttgctggttttgcgattcttca
attaaggagtgtattaagcttgtgtaattgttaatttctctgtcccactccatccaggtcgtgtgattccaaatctgttc
cagagatttattactccaactagcattccaaggcacagcagtggtgcaaatgagttttccagagcaaccccaaatcccca
ggagctgttgatcctttaggtatctttccacagccaggattcttgcctggagctgcttgatgccccagactgtgagttgc
aacagatgctgttgcgcctcaatagccctcagcaaattgttctgctgctgcactataccagacaataattgtctggcctg
taccgtcagcgtcattgacgctgcgcccatagtgcttcctgctgctcccaagaacccaaggaacaaagctcctattccca

ctgctcttttttctctctgcaccactcttctctttgccttggtgggtgctactcctaatggttcaattttttactacttta
tatttatataattcacttctccaattgtccctcatatctcctcctccaggtctgaagatcagcggccgcttgctgtgcgg
tggtcttactttgttttgctcttcctctatcttgtctaaagcttccttggtgtcttttatctctatcctttgatgcaca
caatagagggttgctactgtattatataatgatctaagttcttctgatcctgtctgaagggatggttgtagctgtcccag
tatttgtctacagccttctgatgtttctaacaggccaggattaactgcgaatcgttctagctccctgcttgcccatacta
tatgttttaatttatatttttctttccccctggccttaaccgaattttttcccatcgcgatctaattctcccccgctta
atactgacgctctcgcacccatctctctccttctagcctccgctagtcaaaatttttggcgtactcaccagtcgccgccc
ctcgcctcttgccgtgcgcgcttcagcaagccgagtcctgcgtcgagagagctcctctggtttccctttcgctttcaagt
ccctgttcgggcgccactgctagagattttccacactgactaaaagggtctgagggatctctagttaccagagtcacaca
acagacgggcacacactacttgaagcactcaaggcaagctttattgaggcttaagcagtgggttccctagttagccagag
agctcccaggctcagatctggtctaaccagagagacccgtttatgtatcgagctaggcacttaaatacaatatctctgca
atgcggcaattcagtggttcgtccaatccatgtcagacccgtctgttgccttcctaataaggcacgatcgtaccacctta
cttccaccaatcggcatgcacggtgcttttctctccttgtaaggcatgttgctaactcatcgttaccatgttgcaagac
tacaagagtattgcataagactacattaagcttgcagctccagcttttgttccctttagtgagggttaattgcgcgcttg
gcgtaatcatggtcatagctgtttcctgtgtgaaattgttatccgctcacaattccacacaacatacgagccggaagcat
aaagtgtaaagcctggggtgcctaatgagtgagctaactcacattaattgcgttgcgctcactgcccgctttccagtcgg
gaaacctgtcgtgccagctgcattaatgaatcggccaacgcgcggggagaggcggtttgcgtattgggcgctcttccgct
tcctcgctcactgactcgctgcgctcggtcgttcggctgcggcgagcggtatcagctcactcaaaggcggtaatacggtt
atccacagaatcaggggataacgcaggaaagaacatgtgagcaaaaggccagcaaaaggccaggaaccgtaaaaaggccg
cgttgctggcgtttttccataggctccgcccccctgacgagcatcacaaaaatcgacgctcaagtcagaggtggcgaaac
ccgacaggactataaagataccaggcgtttccccctggaagctccctcgtgcgctctcctgttccgaccctgccgcttac
cggatacctgtccgcctttctcccttcgggaagcgtggcgctttctcatagctcacgctgtaggtatctcagttcggtgt
aggtcgttcgctccaagctgggctgtgtgcacgaaccccccgttcagcccgaccgctgcgccttatccggtaactatcgt
cttgagtccaacccggtaagacacgacttatcgccactggcagcagccactggtaacaggattagcagagcgaggtatgt
aggcggtgctacagagttcttgaagtggtggcctaactacggctacactagaagaacagtatttggtatctgcgctctgc
tgaagccagttaccttcggaaaaagagttggtagctcttgatccggcaaacaaaccaccgctggtagcggtggtttttt
gtttgcaagcagcagattacgcgcagaaaaaaaggatctcaagaagatcctttgatcttttctacggggtctgacgctca
gtggaacgaaaactcacgttaagggattttggtcatgagattatcaaaaaggatcttcacctagatccttttaaattaaa
aatgaagttttaaatcaatctaaagtatatatgagtaaacttggtctgacagttaccaatgcttaatcagtgaggcacct
atctcagcgatctgtctatttcgttcatccatagttgcctgactccccgtcgtgtagataactacgatacgggagggctt
accatctggccccagtgctgcaatgataccgcgagacccacgctcaccggctccagatttatcagcaataaaccagccag
ccggaagggccgagcgcagaagtggtcctgcaactttatccgcctccatccagtctattaattgttgccgggaagctaga
gtaagtagttcgccagttaatagtttgcgcaacgttgttgccattgctacaggcatcgtggtgtcacgctcgtcgtttgg
tatggcttcattcagctccggttcccaacgatcaaggcgagttacatgatcccccatgttgtgcaaaaaagcggttagct
ccttcggtcctccgatcgttgtcagaagtaagttggccgcagtgttatcactcatggttatggcagcactgcataattct
cttactgtcatgccatccgtaagatgcttttctgtgactggtgagtactcaaccaagtcattctgagaatagtgtatgcg
gcgaccgagttgctcttgcccggcgtcaatacgggataataccgcgccacatagcagaactttaaaagtgctcatcattg
gaaaacgttcttcggggcgaaaactctcaaggatcttaccgctgttgagatccagttcgatgtaacccactcgtgcaccc
aactgatcttcagcatcttttactttcaccagcgtttctgggtgagcaaaaacaggaaggcaaaatgccgcaaaaaaggg

aataagggcgacacggaaatgttgaatactcatactcttcctttttcaatattattgaagcatttatcagggttattgtc
tcatgagcggatacatatttgaaaggcctccaaaaaagcctcctcactacttctggaatagctcagaggccgaggcggcc
tcggcctctgcataaat

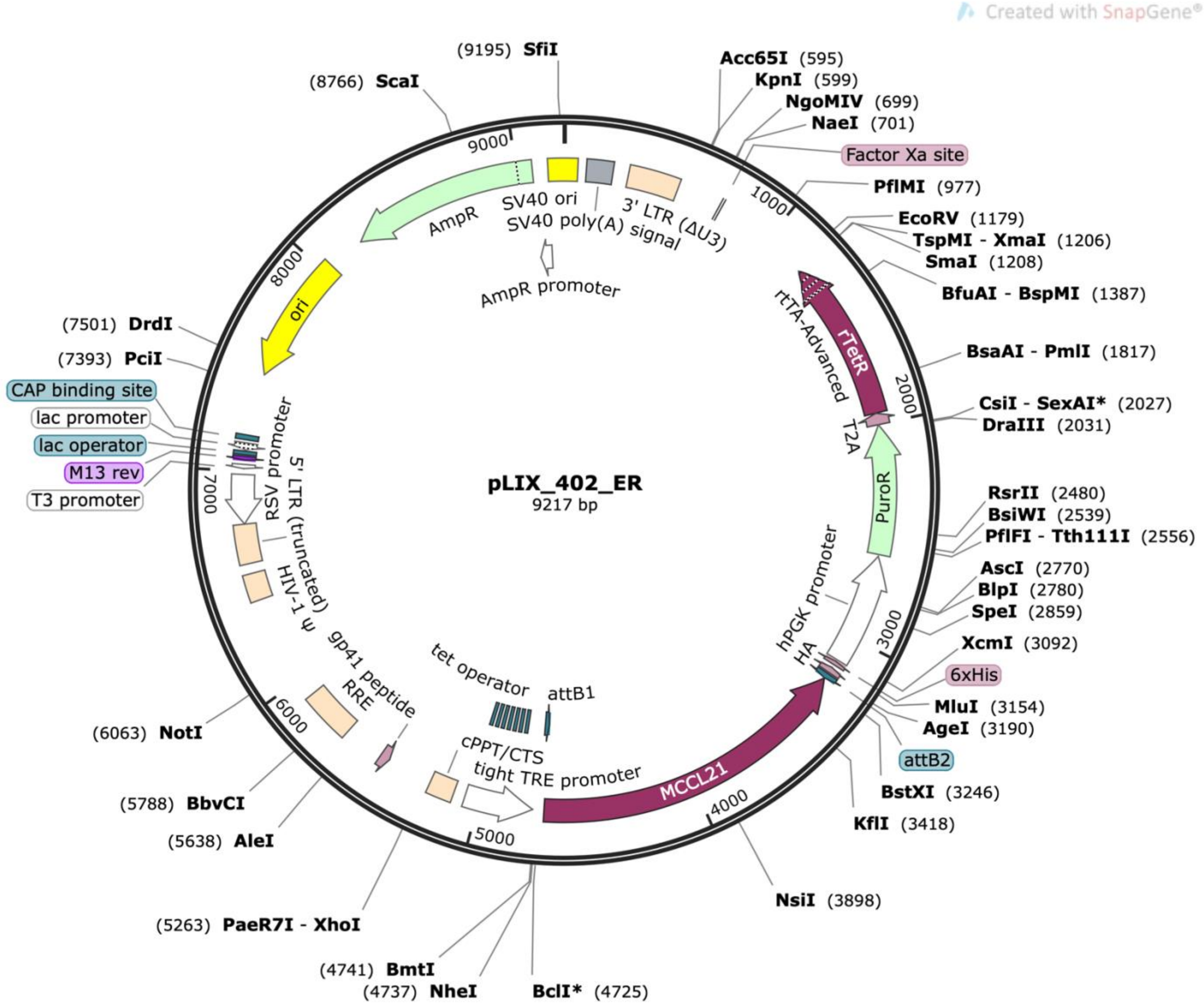

**>pLIX_402_GFP (8453 bp)**

gaaggcctcctggcgccgcaccggcccaaggagcccgcgtggttcctggccaccgtcggcgtctcgcccgaccaccaggg
caagggtctgggcagcgccgtcgtgctccccggagtggaggcggccgagcgcgccggggtgcccgccttcctggagacct
ccgcgccccgcaacctccccttctacgagcggctcggcttcaccgtcaccgccgacgtcgaggtgcccgaaggaccgcgc
acctggtgcatgacccgcaagcccggtgccgaaggtagaggttctctcctcacttgtggtgatgttgaagaaaaccctgg
tccaatgtctagactggacaagagcaaagtcataaacggagctctggaattactcaatggtgtcggtatcgaaggcctga
cgacaaggaaactcgctcaaaagctgggagttgagcagcctaccctgtactggcacgtgaagaacaagcgggccctgctc
gatgccctgccaatcgagatgctggacaggcatcatacccacttctgccccctggaaggcgagtcatggcaagactttct
gcggaacaacgccaagtcataccgctgtgctctcctctcacatcgcgacggggctaaagtgcatctcggcacccgcccaa
cagagaaacagtacgaaaccctggaaaatcagctcgcgttcctgtgtcagcaaggcttctccctggagaacgcactgtac
gctctgtccgccgtgggccactttacactgggctgcgtattggaggaacaggagcatcaagtagcaaaagaggaaagaga
gacacctaccaccgattctatgcccccacttctgagacaagcaattgagctgttcgaccggcagggagccgaacctgcct
tccttttcggcctggaactaatcatatgtggcctggagaaacagctaaagtgcgaaagcggcgggccgaccgacgccctt
gacgattttgacttagacatgctcccagccgatgcccttgacgactttgaccttgatatgctgcctgctgacgctcttga
cgattttgaccttgacatgctccccgggtaactaagtaaggatcgatccaagatatcgtattcttaactatgttgctcct
tttacgctatgtggatacgctgctttaatgcctttgtatcatgctattgcttcccgtatggctttcattttctcctcctt
gtataaatcctggttgctgtctctttatgaggagttgtggcccgttgtcaggcaacgtggcgtggtgtgcactgtgtttg
ctgacgcaacccccactggttggggcattgccaccacctgtcagctcctttccgggactttcgctttccccctccctatt
gccacggcggaactcatcgccgcctgccttgcccgctgctggacaggggctcggctgttgggcactgacaattccgtggt
gttgtcggggaagctgacgtcctttccatggctgctcgcctgtgttgccacctggattctgcgcgggacgtccttctgct
acgtcccttcggccctcaatccagcggaccttccttcccgcggcctgctgccggctctgcggcctcttccgcgtcttcgc
cttcgccctcagacgagtcggatctccctttgggccgcctccccgcctgtttcgcctcggcgtccggactagaggtacct
ttaagaccaatgacttacaaggcagctgtagatcttagccactttttaaaagaaaaggggggactggaagggctaattca
ctcccaacgaagacaagatctgctttttgcttgtactgggtctctctggttagaccagatctgagcctgggagctctctg
gctaactagggaacccactgcttaagcctcaataaagcttgccttgagtgcttcaagtagtgtgtgcccgtctgttgtgt
gactctggtaactagagatccctcagacccttttagtcagtgtggaaaatctctagcagtagtagttcatgtcatcttat
tattcagtatttataacttgcaaagaaatgaatatcagagagtgagaggaacttgtttattgcagcttataatggttaca
aataaagcaatagcatcacaaatttcacaaataaagcatttttttcactgcattctagttgtggtttgtccaaactcatc
aatgtatcttatcatgtctggctctagctatcccgcccctaactccgcccatcccgcccctaactccgcccagttccgcc
cattctccgccccatggctgactaattttttttatttatgcagaggccgaggccgcctcggcctctgagctattccagaa
gtagtgaggaggcttttttggaggcctttcaaatatgtatccgctcatgagacaataaccctgataaatgcttcaataat
attgaaaaaggaagagtatgagtattcaacatttccgtgtcgcccttattcccttttttgcggcattttgccttcctgtt
tttgctcacccagaaacgctggtgaaagtaaaagatgctgaagatcagttgggtgcacgagtgggttacatcgaactgga
tctcaacagcggtaagatccttgagagttttcgccccgaagaacgttttccaatgatgagcacttttaaagttctgctat
gtggcgcggtattatcccgtattgacgccgggcaagagcaactcggtcgccgcatacactattctcagaatgacttggtt
gagtactcaccagtcacagaaaagcatcttacggatggcatgacagtaagagaattatgcagtgctgccataaccatgag
tgataacactgcggccaacttacttctgacaacgatcggaggaccgaaggagctaaccgcttttttgcacaacatggggg
atcatgtaactcgccttgatcgttgggaaccggagctgaatgaagccataccaaacgacgagcgtgacaccacgatgcct
gtagcaatggcaacaacgttgcgcaaactattaactggcgaactacttactctagcttcccggcaacaattaatagactg

gatggaggcggataaagttgcaggaccacttctgcgctcggcccttccggctggctggtttattgctgataaatctggag
ccggtgagcgtgggtctcgcggtatcattgcagcactggggccagatggtaagccctcccgtatcgtagttatctacacg
acggggagtcaggcaactatggatgaacgaaatagacagatcgctgagataggtgcctcactgattaagcattggtaact
gtcagaccaagtttactcatatatactttagattgatttaaaacttcatttttaatttaaaaggatctaggtgaagatcc
tttttgataatctcatgaccaaaatcccttaacgtgagttttcgttccactgagcgtcagaccccgtagaaaagatcaaa
ggatcttcttgagatcctttttttctgcgcgtaatctgctgcttgcaaacaaaaaaaccaccgctaccagcggtggtttg
tttgccggatcaagagctaccaactctttttccgaaggtaactggcttcagcagagcgcagataccaaatactgttcttc
tagtgtagccgtagttaggccaccacttcaagaactctgtagcaccgcctacatacctcgctctgctaatcctgttacca
gtggctgctgccagtggcgataagtcgtgtcttaccgggttggactcaagacgatagttaccggataaggcgcagcggtc
gggctgaacggggggttcgtgcacacagcccagcttggagcgaacgacctacaccgaactgagatacctacagcgtgagc
tatgagaaagcgccacgcttcccgaagggagaaaggcggacaggtatccggtaagcggcagggtcggaacaggagagcgc
acgagggagcttccagggggaaacgcctggtatctttatagtcctgtcgggtttcgccacctctgacttgagcgtcgatt
tttgtgatgctcgtcaggggggcggagcctatggaaaaacgccagcaacgcggcctttttacggttcctggccttttgct
ggccttttgctcacatgttctttcctgcgttatcccctgattctgtggataaccgtattaccgcctttgagtgagctgat
accgctcgccgcagccgaacgaccgagcgcagcgagtcagtgagcgaggaagcggaagagcgcccaatacgcaaaccgcc
tctccccgcgcgttggccgattcattaatgcagctggcacgacaggtttcccgactggaaagcgggcagtgagcgcaacg
caattaatgtgagttagctcactcattaggcaccccaggctttacactttatgcttccggctcgtatgttgtgtggaatt
gtgagcggataacaatttcacacaggaaacagctatgaccatgattacgccaagcgcgcaattaaccctcactaaaggga
acaaaagctggagctgcaagcttaatgtagtcttatgcaatactcttgtagtcttgcaacatggtaacgatgagttagca
acatgccttacaaggagagaaaaagcaccgtgcatgccgattggtggaagtaaggtggtacgatcgtgccttattaggaa
ggcaacagacgggtctgacatggattggacgaaccactgaattgccgcattgcagagatattgtatttaagtgcctagct
cgatacataaacgggtctctctggttagaccagatctgagcctgggagctctctggctaactagggaacccactgcttaa
gcctcaataaagcttgccttgagtgcttcaagtagtgtgtgcccgtctgttgtgtgactctggtaactagagatccctca
gacccttttagtcagtgtggaaaatctctagcagtggcgcccgaacagggacttgaaagcgaaagggaaaccagaggagc
tctctcgacgcaggactcggcttgctgaagcgcgcacggcaagaggcgaggggcggcgactggtgagtacgccaaaaatt
ttgactagcggaggctagaaggagagagatgggtgcgagagcgtcagtattaagcgggggagaattagatcgcgatggga
aaaaattcggttaaggccagggggaaagaaaaaatataaattaaaacatatagtatgggcaagcagggagctagaacgat
tcgcagttaatcctggcctgttagaaacatcagaaggctgtagacaaatactgggacagctacaaccatcccttcagaca
ggatcagaagaacttagatcattatataatacagtagcaaccctctattgtgtgcatcaaaggatagagataaaagacac
caaggaagctttagacaagatagaggaagagcaaaacaaaagtaagaccaccgcacagcaagcggccgctgatcttcaga
cctggaggaggagatatgagggacaattggagaagtgaattatataaatataaagtagtaaaaattgaaccattaggagt
agcacccaccaaggcaaagagaagagtggtgcagagagaaaaaagagcagtgggaataggagctttgttccttgggttct
tgggagcagcaggaagcactatgggcgcagcgtcaatgacgctgacggtacaggccagacaattattgtctggtatagtg
cagcagcagaacaatttgctgagggctattgaggcgcaacagcatctgttgcaactcacagtctggggcatcaagcagct
ccaggcaagaatcctggctgtggaaagatacctaaaggatcaacagctcctggggatttggggttgctctggaaaactca
tttgcaccactgctgtgccttggaatgctagttggagtaataaatctctggaacagatttggaatcacacgacctggatg
gagtgggacagagaaattaacaattacacaagcttaatacactccttaattgaagaatcgcaaaaccagcaagaaaagaa
tgaacaagaattattggaattagataaatgggcaagtttgtggaattggtttaacataacaaattggctgtggtatataa
aattattcataatgatagtaggaggcttggtaggtttaagaatagtttttgctgtactttctatagtgaatagagttagg

cagggatattcaccattatcgtttcagacccacctcccaaccccgaggggacaattctcgacctcgagacaaatggcagt
attcatccacaattttaaaagaaaaggggggattggggggtacagtgcaggggaaagaatagtagacataatagcaacag
acatacaaactaaagaattacaaaaacaaattacaaaaattcaaaattttcgggtttattacagggacagcagagatcca
ctttggccgcgaatcgatatgtcgagtttactccctatcagtgatagagaacgtatgtcgagtttactccctatcagtga
tagagaacgatgtcgagtttactccctatcagtgatagagaacgtatgtcgagtttactccctatcagtgatagagaacg
tatgtcgagtttactccctatcagtgatagagaacgtatgtcgagtttatccctatcagtgatagagaacgtatgtcgag
tttactccctatcagtgatagagaacgtatgtcgaggtaggcgtgtacggtgggaggcctatataagcagagctcgttta
gtgaaccgtcagatcgcctggagaattggctagcatcgattgatcaacaagtttgtacaaaaaagttggcatggtgagca
agggcgaggagctgttcaccggggtggtgcccatcctggtcgagctggacggcgacgtaaacggccacaagttcagcgtg
tccggcgagggcgagggcgatgccacctacggcaagctgaccctgaagttcatctgcaccaccggcaagctgcccgtgcc
ctggcccaccctcgtgaccaccctgacctacggcgtgcagtgcttcagccgctaccccgaccacatgaagcagcacgact
tcttcaagtccgccatgcccgaaggctacgtccaggagcgcaccatcttcttcaaggacgacggcaactacaagacccgc
gccgaggtgaagttcgagggcgacaccctggtgaaccgcatcgagctgaagggcatcgacttcaaggaggacggcaacat
cctggggcacaagctggagtacaactacaacagccacaacgtctatatcatggccgacaagcagaagaacggcatcaagg
tgaacttcaagatccgccacaacatcgaggacggcagcgtgcagctcgccgaccactaccagcagaacacccccatcggc
gacggccccgtgctgctgcccgacaaccactacctgagcacccagtccgccctgagcaaagaccccaacgagaagcgcga
tcacatggtcctgctggagttcgtgaccgccgccgggatcactctcggcatggacgagctgtacaagccaactttcttgt
acaaagtggttaccggttatccgtatgatgtgccggattatgcgtaaacgcgtccggtccaccaccaccaccaccactaa
ggatccggggttggggttgcgccttttccaaggcagccctgggtttgcgcagggacgcggctgctctgggcgtggttccg
ggaaacgcagcggcgccgaccctgggtctcgcacattcttcacgtccgttcgcagcgtcacccggatcttcgccgctacc
cttgtgggccccccggcgacgcttcctgctccgcccctaagtcgggaaggttccttgcggttcgcggcgtgccggacgtg
acaaacggaagccgcacgtctcactagtaccctcgcagacggacagcgccagggagcaatggcagcgcgccgaccgcgat
gggctgtggccaatagcggctgctcagcagggcgcgccgagagcagcggccgggaaggggcggtgcgggaggcggggtgt
ggggcggtagtgtgggccctgttcctgcccgcgcggtgttccgcattctgcaagcctccggagcgcacgtcggcagtcgg
ctccctcgttgaccgaatcaccgacctctctccccagcaattcaccatgaccgagtacaagcccacggtgcgcctcgcca
cccgcgacgacgtccccagggccgtacgcaccctcgccgccgcgttcgccgactaccccgccacgcgccacaccgtcgat
ccggaccgccacatcgagcgggtcaccgagctgcaagaactcttcctcacgcgcgtcgggctcgacatcggcaaggtgtg
ggtcgcggacgacggcgccgcggtggcggtctggaccacgccggagagcgtcgaagcgggggcggtgttcgccgagatcg
gcccgcgcatggccgagttgagcggttcccggctggccgcgcagcaacagatg

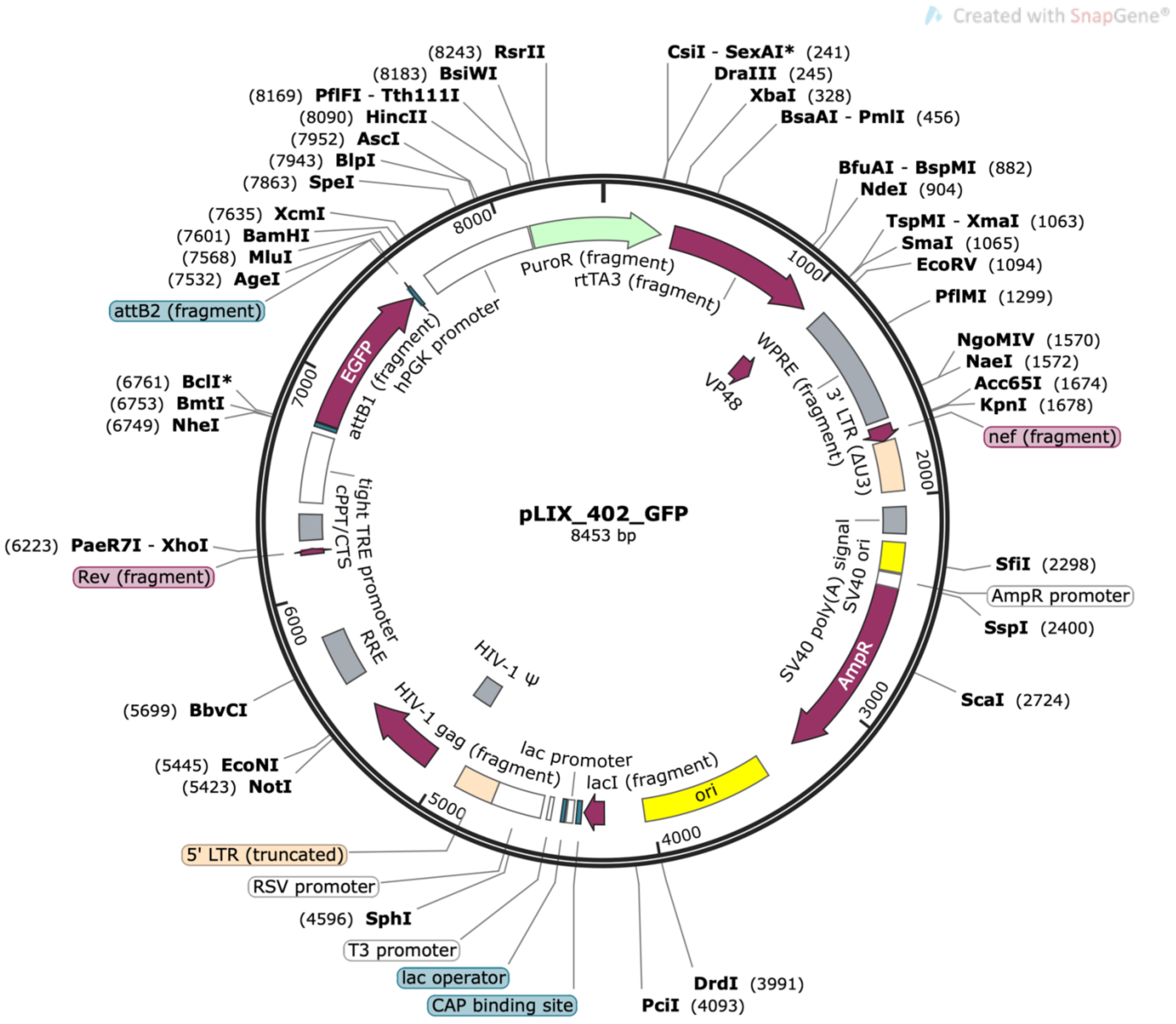
Created with SnapGene®
pLIX_402_GFP
8453 bp
(8243) RsrII
(8183) BsiWI
(8169) PflFI - Tth111I
(8090) HincII
(7952) AscI
(7943) BlpI
(7863) SpeI
(7635) XcmI
(7601) BamHI
(7568) MluI
(7532) AgeI
attB2 (fragment)
(6761) BclI*
(6753) BmtI
(6749) NheI
(6223) PaeR7I - XhoI
Rev (fragment)
(5699) BbvCI
(5445) EcoNI
(5423) NotI
5' LTR (truncated)
RSV promoter
(4596) SphI
T3 promoter
lac operator
CAP binding site
CsiI - SexAI* (241)
DraIII (245)
XbaI (328)
BsaAI - PmlI (456)
BfuAI - BspMI (882)
NdeI (904)
TspMI - XmaI (1063)
SmaI (1065)
EcoRV (1094)
PflMI (1299)
NgoMIV (1570)
NaeI (1572)
Acc65I (1674)
KpnI (1678)
nef (fragment)
SfiI (2298)
AmpR promoter
SspI (2400)
ScaI (2724)
DrdI (3991)
PciI (4093)
PuroR (fragment)
rtTA3 (fragment)
VP48
WPRE (fragment)
3' LTR (ΔU3)
SV40 poly(A) signal
SV40 ori
AmpR
ori
lacI (fragment)
lac promoter
HIV-1 gag (fragment)
HIV-1 Ψ
RRE
tight TRE promoter
cPPT/CTS
attB1 (fragment)
hPGK promoter
EGFP
1000
2000
3000
4000
5000
6000
7000
8000

**>pLIX_403_hWNT5B (9106 bp)**

ggcgatctgacggttcactaaacgagctctgcttatataggcctcccaccgtacacgcctacctcgacatacgttctcta
tcactgatagggagtaaactcgacatacgttctctatcactgatagggataaactcgacatacgttctctatcactgata
gggagtaaactcgacatacgttctctatcactgatagggagtaaactcgacatacgttctctatcactgatagggagtaa
actcgacatcgttctctatcactgatagggagtaaactcgacatacgttctctatcactgatagggagtaaactcgacat
atcgattcgcggccaaagtggatctctgctgtccctgtaataaacccgaaaattttgaatttttgtaatttgtttttgta
attctttagtttgtatgtctgttgctattatgtctactattctttcccctgcactgtacccccaatccccccttttctt
ttaaaattgtggatgaatactgccatttgtctcgaggtcgagaattgtcccctcggggttgggaggtgggtctgaaacga
taatggtgaatatccctgcctaactctattcactatagaaagtacagcaaaaactattcttaaacctaccaagcctccta
ctatcattatgaataattttatataccacagccaatttgttatgttaaaccaattccacaaacttgcccatttatctaat
tccaataattcttgttcattcttttcttgctggttttgcgattcttcaattaaggagtgtattaagcttgtgtaattgtt
aatttctctgtcccactccatccaggtcgtgtgattccaaatctgttccagagatttattactccaactagcattccaag
gcacagcagtggtgcaaatgagttttccagagcaaccccaaatccccaggagctgttgatcctttaggtatctttccaca
gccaggattcttgcctggagctgcttgatgccccagactgtgagttgcaacagatgctgttgcgcctcaatagccctcag
caaattgttctgctgctgcactataccagacaataattgtctggcctgtaccgtcagcgtcattgacgctgcgcccatag
tgcttcctgctgctcccaagaacccaaggaacaaagctcctattcccactgctctttttctctctgcaccactcttctc
tttgccttggtgggtgctactcctaatggttcaattttactactttatatttatataattcacttctccaattgtccct
catatctcctcctccaggtctgaagatcagcggccgcttgctgtgcggtggtcttactttgttttgctcttcctctatc
ttgtctaaagcttccttggtgtcttttatctctatcctttgatgcacacaatagagggttgctactgtattatataatga
tctaagttcttctgatcctgtctgaagggatggttgtagctgtcccagtatttgtctacagccttctgatgtttctaaca
ggccaggattaactgcgaatcgttctagctccctgcttgcccatactatatgttttaatttatatttttctttcccccct
ggccttaaccgaattttttcccatcgcgatctaattctcccccgcttaatactgacgctctcgcacccatctctctcctt
ctagcctccgctagtcaaaattttggcgtactcaccagtcgccgcccctcgcctcttgccgtgcgcgcttcagcaagcc
gagtcctgcgtcgagagagctcctctggtttccctttcgctttcaagtccctgttcgggcgccactgctagagattttcc
acactgactaaaagggtctgagggatctctagttaccagagtcacacaacagacgggcacacactacttgaagcactcaa
ggcaagctttattgaggcttaagcagtgggttccctagttagccagagagctcccaggctcagatctggtctaaccagag
agacccgtttatgtatcgagctaggcacttaaatacaatatctctgcaatgcggcaattcagtggttcgtccaatccatg
tcagacccgtctgttgccttcctaataaggcacgatcgtaccaccttacttccaccaatcggcatgcacggtgcttttc
tctccttgtaaggcatgttgctaactcatcgttaccatgttgcaagactacaagagtattgcataagactacattaagct
tgcagctccagcttttgttccctttagtgagggttaattgcgcgcttggcgtaatcatggtcatagctgtttcctgtgtg
aaattgttatccgctcacaattccacacaacatacgagccggaagcataaagtgtaaagcctggggtgcctaatgagtga
gctaactcacattaattgcgttgcgctcactgcccgctttccagtcgggaaacctgtcgtgccagctgcattaatgaatc
ggccaacgcgcggggagaggcggtttgcgtattgggcgctcttccgcttcctcgctcactgactcgctgcgctcggtcgt
tcggctgcggcgagcggtatcagctcactcaaaggcggtaatacggttatccacagaatcaggggataacgcaggaaaga
acatgtgagcaaaaggccagcaaaaggccaggaaccgtaaaaaggccgcgttgctggcgtttttccataggctccgcccc
cctgacgagcatcacaaaaatcgacgctcaagtcagaggtggcgaaacccgacaggactataaagataccaggcgtttcc
ccctggaagctccctcgtgcgctctcctgttccgaccctgccgcttaccggatacctgtccgcctttctcccttcgggaa
gcgtggcgctttctcatagctcacgctgtaggtatctcagttcggtgtaggtcgttcgctccaagctgggctgtgtgcac
gaaccccccgttcagcccgaccgctgcgccttatccggtaactatcgtcttgagtccaacccggtaagacacgacttatc

gccactggcagcagccactggtaacaggattagcagagcgaggtatgtaggcggtgctacagagttcttgaagtggtggc
ctaactacggctacactagaagaacagtatttggtatctgcgctctgctgaagccagttaccttcggaaaaagagttggt
agctcttgatccggcaaacaaaccaccgctggtagcggtggtttttttgtttgcaagcagcagattacgcgcagaaaaaa
aggatctcaagaagatcctttgatcttttctacggggtctgacgctcagtggaacgaaaactcacgttaagggattttgg
tcatgagattatcaaaaaggatcttcacctagatccttttaaattaaaaatgaagttttaaatcaatctaaagtatatat
gagtaaacttggtctgacagttaccaatgcttaatcagtgaggcacctatctcagcgatctgtctatttcgttcatccat
agttgcctgactccccgtcgtgtagataactacgatacgggagggcttaccatctggccccagtgctgcaatgataccgc
gagacccacgctcaccggctccagatttatcagcaataaaccagccagccggaagggccgagcgcagaagtggtcctgca
actttatccgcctccatccagtctattaattgttgccgggaagctagagtaagtagttcgccagttaatagtttgcgcaa
cgttgttgccattgctacaggcatcgtggtgtcacgctcgtcgtttggtatggcttcattcagctccggttcccaacgat
caaggcgagttacatgatcccccatgttgtgcaaaaaagcggttagctccttcggtcctccgatcgttgtcagaagtaag
ttggccgcagtgttatcactcatggttatggcagcactgcataattctcttactgtcatgccatccgtaagatgcttttc
tgtgactggtgagtactcaaccaagtcattctgagaatagtgtatgcggcgaccgagttgctcttgcccggcgtcaatac
gggataataccgcgccacatagcagaactttaaaagtgctcatcattggaaaacgttcttcggggcgaaaactctcaagg
atcttaccgctgttgagatccagttcgatgtaacccactcgtgcacccaactgatcttcagcatcttttactttcaccag
cgtttctgggtgagcaaaaacaggaaggcaaaatgccgcaaaaaagggaataagggcgacacggaaatgttgaatactca
tactcttcctttttcaatattattgaagcatttatcagggttattgtctcatgagcggatacatatttgaaaggcctcca
aaaaagcctcctcactacttctggaatagctcagaggccgaggcggcctcggcctctgcataaataaaaaaaattagtca
gccatggggcggagaatgggcggaactgggcggagttaggggcgggatgggcggagttaggggcgggatagctagagcca
gacatgataagatacattgatgagtttggacaaaccacaactagaatgcagtgaaaaaaatgctttatttgtgaaatttg
tgatgctattgctttatttgtaaccattataagctgcaataaacaagttcctctcactctctgatattcatttctttgca
agttataaatactgaataataagatgacatgaactactactgctagagattttccacactgactaaaagggtctgaggga
tctctagttaccagagtcacacaacagacgggcacacactacttgaagcactcaaggcaagctttattgaggcttaagca
gtgggttccctagttagccagagagctcccaggctcagatctggtctaaccagagagacccagtacaagcaaaaagcaga
tcttgtcttcgttgggagtgaattagcccttccagtccccccttttcttttaaaaagtggctaagatctacagctgcctt
gtaagtcattggtcttaaaggtacctctagtccggacgccgaggcgaaacaggcggggaggcggcccaaagggagatccg
actcgtctgagggcgaaggcgaagacgcggaagaggccgcagagccggcagcaggccgcgggaaggaaggtccgctggat
tgagggccgaagggacgtagcagaaggacgtcccgcgcagaatccaggtggcaacacaggcgagcagccatggaaaggac
gtcagcttccccgacaacaccacggaattgtcagtgcccaacagccgagcccctgtccagcagcgggcaaggcaggcggc
gatgagttccgccgtggcaatagggagggggaaagcgaaagtcccggaaaggagctgacaggtggtggcaatgccccaac
cagtggggggttgcgtcagcaaacacagtgcacaccacgccacgttgcctgacaacgggccacaactcctcataaagagac
agcaaccaggatttatacaaggaggagaaaatgaaagccatacgggaagcaatagcatgatacaaaggcattaaagcagc
gtatccacatagcgtaaaaggagcaacatagttaagaatacgatatcttggatcgatccttacttagttacccggggagc
atgtcaaggtcaaaatcgtcaagagcgtcagcaggcagcatatcaaggtcaaagtcgtcaagggcatcggctgggagcat
gtctaagtcaaaatcgtcaagggcgtcggtcggcccgccgctttcgcactttagctgtttctccaggccacatatgatta
gttccaggccgaaaaggaaggcaggttcggctccctgccggtcgaacagctcaattgcttgtctcagaagtgggggcata
gaatcggtggtaggtgtctctctttcctcttttgctacttgatgctcctgttcctccaatacgcagcccagtgtaaagtg
gcccacggcggacagagcgtacagtgcgttctccagggagaagccttgctgacacaggaacgcgagctgattttccaggg
tttcgtactgtttctctgttgggcgggtgccgagatgcactttagccccgtcgcgatgtgagaggagagcacagcggtat

gacttggcgttgttccgcagaaagtcttgccatgactcgccttccagggggcagaagtgggtatgatgcctgtccagcat
ctcgattggcagggcatcgagcagggcccgcttgttcttcacgtgccagtacagggtaggctgctcaactcccagctttt
gagcgagtttccttgtcgtcaggccttcgataccgacaccattgagtaattccagagctccgtttatgactttgctcttg
tccagtctagacattggaccagggttttcttcaacatcaccacaagtgaggagagaacctctaccttcggcaccgggctt
gcgggtcatgcaccaggtgcgcggtccttcgggcacctcgacgtcggcggtgacggtgaagccgagccgctcgtagaagg
ggaggttgcggggcgcggaggtctccaggaaggcgggcaccccggcgcgctcggccgcctccactccggggagcacgacg
gcgctgcccagacccttgccctggtggtcgggcgagacgccgacggtggccaggaaccacgcgggctccttgggccggtg
cggcgccaggaggccttccatctgttgctgcgcggccagccgggaaccgctcaactcggccatgcgcgggccgatctcgg
cgaacaccgcccccgcttcgacgctctccggcgtggtccagaccgccaccgcggcgccgtcgtccgcgacccacaccttg
ccgatgtcgagcccgacgcgcgtgaggaagagttcttgcagctcggtgacccgctcgatgtggcggtccggatcgacggt
gtggcgcgtggcggggtagtcggcgaacgcggcggcgagggtgcgtacggccctggggacgtcgtcgcgggtggcgaggc
gcaccgtgggcttgtactcggtcatggtgaattgctggggagagaggtcggtgattcggtcaacgagggagccgactgcc
gacgtgcgctccggaggcttgcagaatgcggaacaccgcgcgggcaggaacagggcccacactaccgccccacaccccgc
ctcccgcaccgccccttcccggccgctgctctcggcgcgccctgctgagcagccgctattggccacagcccatcgcggtc
ggcgcgctgccattgctccctggcgctgtccgtctgcgagggtactagtgagacgtgcggcttccgtttgtcacgtccgg
cacgccgcgaaccgcaaggaaccttcccgacttaggggcggagcaggaagcgtcgccgggggcccacaagggtagcggc
gaagatccgggtgacgctgcgaacggacgtgaagaatgtgcgagacccagggtcggcgccgctgcgtttcccggaaccac
gcccagagcagccgcgtccctgcgcaaacccagggctgccttggaaaaggcgcaaccccaaccccggatccgaccggttg
caaccactttgtacaagaaagctgaacgagaaacgtaaaatgatataaatatcaatatattaaattagattttgcataaa
aaacagactacataatactgtaaaacacaacatatccagtcactatggtcgacctgcagactggctgtgtataagggagc
ctgacatctatttacagatgtactggtccacgatctccgtgcacttcttacacctgacgaagcagcaccagtggaacttg
cagtggcagcgctccacctgcacgctcttgaactggttgtagccacgcccgcagcacatgagctcacagccatccatgcc
ctccgaggtcttgttgcagaggcggccctgcgtgcccagggagcccgtgctctcgttgcgcaggcagtagtcggggctgg
ggtccacatagaccaggtcctccggggtgggctgggtgaagcggctgttgaccagctccagccggcccttgcgggtgacg
cgcatggcggccgcgctgtcgtacttctccttcagccggtccccgaccttgcggaactcggccagctgcagccagcaggt
cttgaggctgcaggaccccgagacgccgtggcatttgcaggctacgtctgccatcttatacacagccctgcgaccggcct
cgttgttttgcaggttcatgagcacccggccctgctcctctgatcctttggcaaagttcttctctcgctcccgggcatcc
acaaactccttggcgaagcggtagccgtactccacgttgtccccacagccgccccacagccagtcccgggcaggtcctt
gggccgcgccgtccggctgcagccgcaggtggagagctcgccctcgcggcaggcccggctgatggcgttgaccacgcccg
cggcgctcaccgcgtgggtgaaggcggtctctcggctgcctatctgcatgactctcccaaagacagatgcgttgtccgct
gtgctgcaattccaccgccgctgccggaactggtgctggcattccttgatgccagtcttggctccctcccctatgtaggc
catgtgctcctggtacaattggcacagcttcctctggccaggggagagcccgggaagctgactgcacacgggctgggcac
cgatgataaacatctcgggtctctgcaccgggttcaaagctaatgaccaccaggagttggcgtctgtcagaagctgagcc
cagctggacagcagagcagccgtgaacagcagcagcaggctgggcatggtggctttagcttccttagctcctgaaaatct
cgacggatcctaactcaaaatccacacattatacgagccggaagcataaagtgtaaagcctggggtgcctaatgcggccg
ccatagtgactggatatgttgtgttttacagtattatgtagtctgtttttttatgcaaaatctaatttaatatattgatat
ttatatcattttacgtttctcgttcagctttttttgtacaaacttgttgatgctagccaattctcca

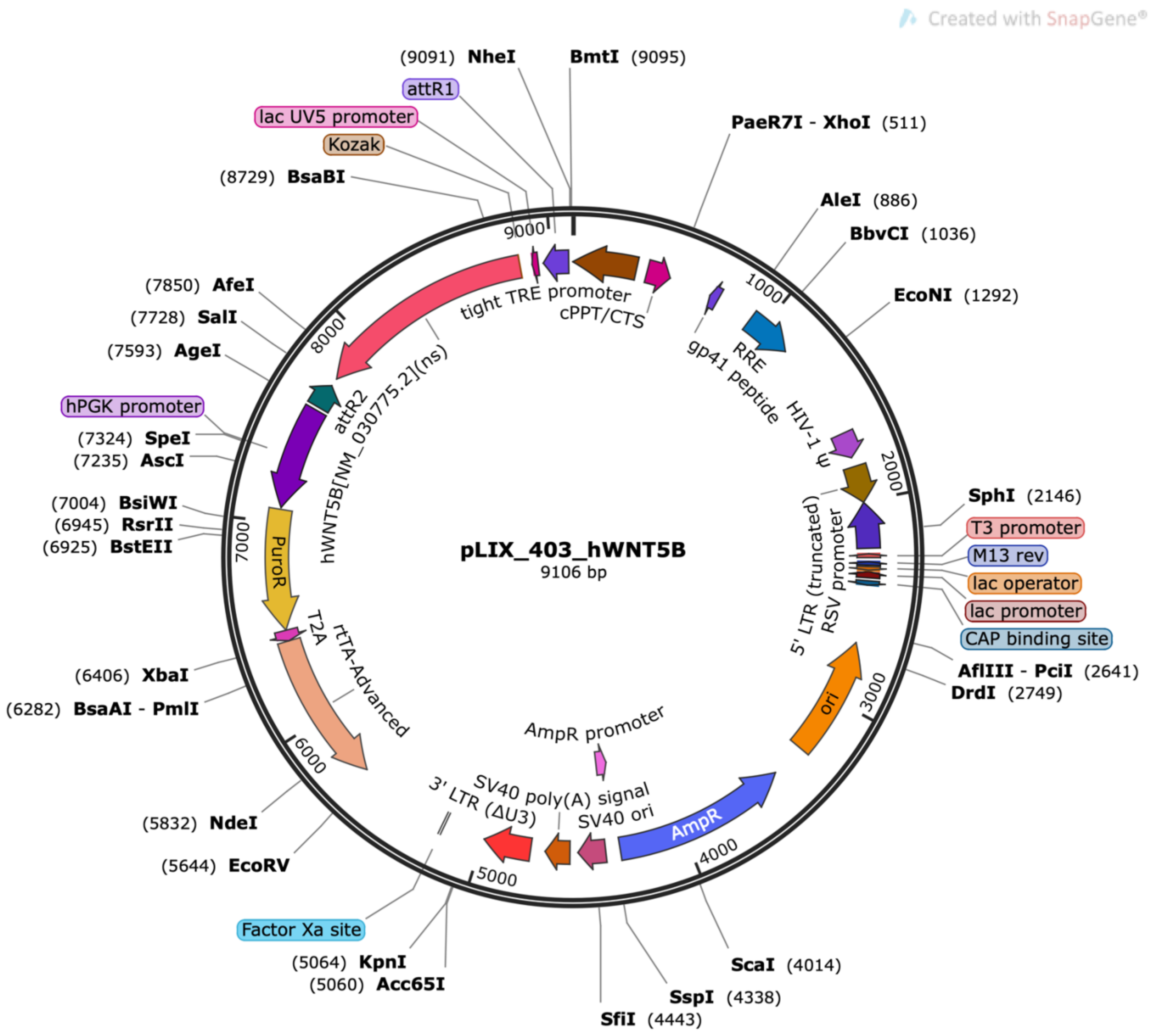
Created with SnapGene®
pLIX_403_hWNT5B
9106 bp
(9091) NheI
BmtI (9095)
attR1
lac UV5 promoter
Kozak
(8729) BsaBI
PaeR7I - XhoI (511)
AleI (886)
BbvCI (1036)
EcoNI (1292)
(7850) AfeI
(7728) SalI
(7593) AgeI
hPGK promoter
(7324) SpeI
(7235) AscI
(7004) BsiWI
(6945) RsrII
(6925) BstEII
(6406) XbaI
(6282) BsaAI - PmlI
(5832) NdeI
(5644) EcoRV
Factor Xa site
(5064) KpnI
(5060) Acc65I
SfiI (4443)
SspI (4338)
ScaI (4014)
SphI (2146)
T3 promoter
M13 rev
lac operator
lac promoter
CAP binding site
AflIII - PciI (2641)
DrdI (2749)
tight TRE promoter
cPPT/CTS
RRE
gp41 peptide
HIV-1 Ψ
5' LTR (truncated)
RSV promoter
ori
AmpR
AmpR promoter
SV40 ori
SV40 poly(A) signal
3' LTR (ΔU3)
rtTA-Advanced
T2A
PuroR
attR2
hWNT5B[NM_030775.2](ns)
1000
2000
3000
4000
5000
6000
7000
8000
9000

**siRNA sequences**

| siRNA target | Sequences |
|---|---|
| siCTRL | commercially available from IDT CAT#: 51-01-14-03 |
| siTCF3-1 | Sense: 5’-CGUUCUGCAUAGAAUUCAAACGAGA-3’<br>Anti-sense: 5’-UCUCGUUUGAAUUCUAUGCAGAACGCA-3’ |
| siTCF3-2 | Sense: 5’-GCAGAGUAAGAUAGAAGACCACCTG-3’<br>Anti-sense: 5’-CAGGUGGUCUUCUAUCUUACUCUGCAG-3’ |
| siTCF7-1 | Sense: 5’ -GGAGAAGCUCUGUUUAUAAAAACAA-3’<br>Anti-sense: 5’-UUGUUUUUAUAAACAGAGCUUCUCCAU-3’ |
| siTCF7-2 | Sense: 5’-GAAAAAGAAAUGCAUUCGGUACUTA-3’<br>Anti-sense: 5’-UAAGUACCGAAUGCAUUUCUUUUUCCU-3’ |

**References**


1. Cheng J, et al. Merkel cell polyomavirus recruits MYCL to the EP400 complex to promote oncogenesis. *Plos Pathog*. 2017;13(10):e1006668.

2. Varghese F, et al. IHC Profiler: An Open Source Plugin for the Quantitative Evaluation and Automated Scoring of Immunohistochemistry Images of Human Tissue Samples. *PLoS ONE*. 2014;9(5):e96801.

3. Alvarez MJ, et al. Functional characterization of somatic mutations in cancer using network-based inference of protein activity. *Nat Genet*. 2016;48(8):838–847.

4. Garcia-Alonso L, et al. Benchmark and integration of resources for the estimation of human transcription factor activities. *Genome Res*. 2019;29(8):1363–1375.

5. Lachmann A, et al. ARACNe-AP: gene network reverse engineering through adaptive partitioning inference of mutual information. *Bioinformatics*. 2016;32(14):2233–2235.
